# Age-dependent changes in the putamen transcriptome and proteome of a transgenic minipig model of Huntington’s disease

**DOI:** 10.64898/2026.11.13.766462

**Authors:** Eva Kamenná, Kateryna Nemesh, Jakub Červenka, David Sekáč, Christian Landles, Anastasiia Kolesnikova, Ivona Valeková, Štefan Juhás, Jana Juhásová, Georgina Osborne, Helena Kupcová Skalníková, Petr Vodička, Jan Motlík, Jiří Klempíř, Zdeňka Ellederová, Jiří Klíma

## Abstract

Huntington’s disease is an inherited neurodegenerative disorder with no disease-modifying treatment available. Preclinical trials of gene therapies rely on large animal models and the transgenic Huntington’s disease (TgHD) minipig is one of them. Minipigs live more than a decade, so the effect of a single dose of gene therapy can be followed for years. The TgHD transgene carries a mixed CAG/CAA repeat that is not expected to expand somatically. However, its putamen, part of the striatum that degenerates first in patients, had not been examined by genome-wide methods, nor beyond 4 years of age. We compared the putamen of transgenic and wild-type minipigs at 4 and 8 years by RNA sequencing and label-free mass spectrometry proteomics, and measured mutant huntingtin by immunoassay. Seven genes were differentially expressed at 4 years and 3,324 at 8 years, and the pathway changes of the transcriptome and the proteome were concordant at that age. The D2 medium spiny neuron marker gene set was reduced at 8 years in both the transcriptome and the proteome. Mitochondrial and oxidative phosphorylation pathways were positively enriched at 4 years and negatively enriched at 8 years. Immune and complement pathways were positively enriched at 8 years, and the number of altered splicing events increased almost fivefold between 4 and 8 years. We detected no age-dependent accumulation of soluble mutant huntingtin. Since the repeat does not expand, these changes developed without somatic expansion, a major driver of progression in human Huntington’s disease. We propose that the reduced neuronal markers reflect a loss of neuronal identity rather than neuronal death. The TgHD minipig would then recapitulate the slow early phase of the disease, when treatment is arguably most likely to help.

## 1. Introduction

Huntington’s disease (HD) is an autosomal dominant neurodegenerative disorder caused by an expanded CAG repeat in the first exon of the huntingtin gene (*HTT*). Inheritance of 40 or more repeats is fully penetrant, and longer alleles cause earlier onset (Bates et al., 2015). The disease affects approximately 1 in 7,300 individuals of European descent (Bates et al., 2015), and although the cause was identified in 1993 (MacDonald et al., 1993), no disease-modifying treatment has reached the clinic. Degeneration is uneven across the brain. The striatum is affected the earliest and most severely, and its medium spiny neurons (MSNs) are selectively vulnerable (Bates et al., 2015). MSNs of the indirect pathway (D2 MSNs) are lost before those of the direct pathway (D1 MSNs) (Dunah et al., 2002; Lee et al., 2020), which is thought to underlie the early hyperkinetic phenotype (Raymond et al., 2011). Single-nucleus studies have recently narrowed the vulnerability further, to indirect-pathway MSNs of the striosome compartment (Matsushima et al., 2023).

How a single expanded repeat produces this pattern of loss is not fully understood, although the biology of the healthy huntingtin protein offers some clues. Huntingtin is a large scaffolding protein with more than 350 binding partners, and it participates in transcriptional regulation, intracellular transport, proteostasis and DNA repair (Donaldson et al., 2026). It is also needed at the synapse for vesicle transport, neurotransmitter release and receptor recycling, so a loss of its normal function may contribute to the phenotype (Barron et al., 2021). However, the expanded polyglutamine tract in mutant huntingtin (mHTT) adds a toxic gain of function that disturbs these processes. Expansion also makes the protein prone to cleavage into N-terminal fragments that form oligomers, aggregates and inclusions in neurons (Bates et al., 2015). Overall, mHTT has been linked to disturbed gene transcription and RNA processing, mitochondrial function, calcium handling, synaptic transmission, DNA-damage repair and immune responses (Bates et al., 2015; Donaldson et al., 2026).

More recently, somatic expansion of the CAG repeat has been proposed as a further disease mechanism. Although it was described in 2007 (Shelbourne et al., 2007), it gained attention only after genome-wide association studies first identified genetic modifiers of age at onset in 2015. Most genes implicated are DNA-repair genes, and CAA interruptions have also been identified as modifiers of age at onset (Lee et al., 2019, 2015). Post-mitotic neurons cannot use replication-coupled repair, and the repeat continues to expand in these cells throughout life, reaching well over a hundred CAG repeats in individual striatal projection neurons. Expansion up to roughly 150 repeats appears to have little effect on the cell, and neurons lose their identity and die only above that threshold (Handsaker et al., 2025; Mätlik et al., 2024). HD is therefore monogenic in origin, but it is now understood to be driven both by protein toxicity and by somatic expansion in the DNA of specific cell types.

Of the processes linked to mHTT, transcriptional dysregulation is one of the earliest (Seredenina and Luthi-Carter, 2012) and is already present in the caudate nucleus of human carriers before the clinical diagnosis (Agus et al., 2019). Mitochondria are affected in several ways. Huntingtin supports mitochondrial transport, fusion and fission, and the mutant protein has been associated with bioenergetic deficits, oxidative stress, disturbed calcium handling and mitochondrial DNA abnormalities (Joshi et al., 2025). Microglial reactivity and altered astrocyte function accompany these changes and have been proposed as both contributors to degeneration and sources of biomarkers in cerebrospinal fluid (CSF) or blood (Blusch and Björkqvist, 2025). Alternative splicing is also disturbed in both the HD cortex and the human and mouse striatum (Lin et al., 2016; Tano et al., 2023).

Animal models play a key role in studying disease mechanisms, as presymptomatic and early symptomatic human striatal tissue is scarce, and late symptomatic tissue is already too degenerated for meaningful analysis (Agus et al., 2019). They provide material across disease stages at the cost of an imperfect phenotype. Animals need repeat lengths well above the human threshold of 40 CAG repeats to reach a phenotype within their lifetime, and they differ from humans in brain anatomy, lifespan and immune biology (Howland et al., 2020; Pouladi et al., 2013). The TgHD Libechov minipig was created as a large-animal model with a gyrencephalic brain for gene-therapy testing, and it was used in the preclinical work on AAV5-miHTT, later AMT-130 (Evers et al., 2018). It expresses a truncated N-terminal fragment of human huntingtin (amino acids 1 to 548) with 124 glutamines. The transgene is a human *HTT* cDNA with a mixed CAG/CAA repeat, under the human *HTT* promoter and integrated as a single copy on chromosome 1 (Baxa et al., 2013). This mixed repeat is stable across generations (Askeland et al., 2018b) and is not expected to expand in somatic cells, so the model represents proteinopathy without somatic expansion, 26 units below the proposed threshold of 150 CAG repeats (Handsaker et al., 2025).

The TgHD minipig has been characterised before with targeted analyses (Askeland et al., 2018b; Baxa et al., 2013). However, the model has not been examined by exploratory, genome-wide profiling, and no characterisation has continued beyond 4 years of age. With a minipig lifespan of 10 to 12 years, this model offers a rare opportunity to monitor gene therapy for years after delivery, yet data on the model at older ages have been missing. Here we characterise the putamen of TgHD and wild-type minipigs at 4 and 8 years of age by bulk RNA sequencing and label-free mass spectrometry proteomics, with qPCR validation, homogeneous time-resolved fluorescence (HTRF) assays for mHTT and assays of nuclear and mitochondrial DNA integrity.

## 2. Materials and methods

### 2.1. Animals and tissue collection

The transgenic Huntington’s disease (TgHD) minipig model was generated by lentiviral transduction of porcine embryos as previously described (Baxa et al., 2013). The transgene is a human huntingtin (*HTT*) complementary DNA encoding the N-terminal fragment of the protein, amino acids 1 to 548, and is driven by the human *HTT* promoter. The polyglutamine tract of 124 glutamines is encoded by a mixed CAG/CAA repeat with the repeating unit CAGCAACAACAGCAGCAA. A single transgene copy is integrated at chromosome 1q24 to q25, and heterozygous animals were used in this study.

Putamen, pre-frontal cortex, motor cortex and mixed cortex were collected from 27 animals at three ages (Supplementary Table S1). There were 4 wild-type (WT) and 4 TgHD animals at 4 years of age, 5 WT and 7 TgHD at 7 years, and 4 WT and 3 TgHD at 8 years. These age groups are referred to as the 4-year, 7-year and 8-year cohorts throughout.

Animals were euthanised and brains were processed by one of two protocols. The 7-year animals were perfused with cold phosphate-buffered saline (PBS) before dissection, whereas 4-year and 8-year brains were removed and dissected directly without perfusion, to shorten the time between euthanasia and brain collection. Tissue was fresh-frozen in liquid nitrogen and stored at −80 °C. All procedures were carried out under approved project protocols 53/2015 and 109/2020 (see Ethics statement).

Of these 27 animals, 12 provided putamen tissue for RNA sequencing (RNA-seq), 3 WT and 3 TgHD at each of 4 and 8 years. All 27 animals were used for proteomics (putamen) and for the huntingtin immunoassays (cortex and putamen, Section 2.2). Quantitative PCR validation was performed on the 7-year and 8-year cohorts (putamen), and DNA integrity and mitochondrial DNA copy number were measured in the 7-year cohort (cortex and putamen).

### 2.2. HTT protein quantification by HTRF

Soluble mutant huntingtin (mHTT) was quantified by homogeneous time-resolved fluorescence (HTRF) in putamen (4, 7 and 8 years), pre-frontal cortex and motor cortex (4 and 8 years) of WT and TgHD animals. Tissues were lysed and the assays were run as described previously (Landles et al., 2024; Landles et al., 2021), with the modifications for minipig, including a lysate concentration of 20% (w/v) (Kolesnikova et al., 2026). Two antibody pairs were used, 2B7-Tb with MW1-d2 for total soluble mHTT and MAB2166-Tb with MAB5490-d2 for the transgenic human HTT protein. Neither HTRF assay detected endogenous *Sus scrofa* HTT. The HTRF signal was normalised to WT after subtraction of the background signal. Genotype and age effects were tested by two-way ANOVA with Bonferroni post hoc correction (Supplementary Table S18).

### 2.3. RNA extraction and sequencing

Total RNA was isolated from approximately 10 mg of frozen putamen tissue using TRI Reagent (Sigma, T9424) at 1.5 mL per 5–10 mg of tissue. Tissue was homogenised by passage through a 23G needle and syringe. Glycogen (25 µg) was added to enhance RNA precipitation, and the pellet was dissolved in RNase-free water. Genomic DNA was removed by dsDNase treatment (Thermo Fisher Scientific, EN0521).

RNA integrity was assessed on an Agilent Bioanalyzer using the RNA 6000 Nano Kit, with RNA integrity number (RIN) values of 4.0 to 8.3, and concentrations were measured with the Qubit RNA BR Assay Kit (Thermo Fisher Scientific, Q10210). Seven of the 12 samples had a RIN below 7, but all 12 yielded libraries that passed quality control.

Library preparation and sequencing were performed by SEQme s.r.o. (Dobříš, Czech Republic). Poly(A) mRNA was selected using the NEBNext Poly(A) mRNA Magnetic Isolation Module (New England Biolabs, E7490). Strand-specific libraries were prepared with the NEBNext Ultra II Directional RNA Library Prep Kit for Illumina (E7760) and indexed with NEBNext Multiplex Oligos for Illumina (96 Unique Dual Index Primer Pairs, E6440). Library quality was verified using the Agilent Bioanalyzer 2100 High Sensitivity DNA Kit, the Invitrogen Collibri Library Quantification Kit and the Qubit 1X dsDNA High-Sensitivity Assay Kit, and mean insert sizes ranged from 481 to 542 bp. Twelve libraries were pooled and sequenced on one lane of a 10B flow cell on an Illumina NovaSeq X Plus (paired-end, 2 × 150 bp). Sequencing produced 99.9 to 122.8 million read pairs per sample (mean 110.4 million, 2.65 billion reads in total).

### 2.4. Read processing and alignment

Adapters and low-quality bases were removed with Trimmomatic v0.39 (Bolger et al., 2014) using sample-specific adapter sequences (ILLUMINACLIP:2:30:10:2:True LEADING:3 TRAILING:3 SLIDINGWINDOW:4:15 MINLEN:36), and 97% of read pairs were retained.

Trimmed reads were checked with FastQC v0.12.1 and MultiQC v1.21 (Ewels et al., 2016), then aligned to the *Sus scrofa* reference genome (Sscrofa11.1) with STAR v2.5.2b (Dobin et al., 2013) in two-pass mode (--twopassMode Basic --alignSJoverhangMin 8 -- alignSJDBoverhangMin 1 --outFilterMismatchNmax 999 -- outFilterMismatchNoverReadLmax 0.04 --alignIntronMin 20 --alignIntronMax 20000 --alignMatesGapMax 5000 --outFilterIntronMotifs RemoveNoncanonical). Uniquely mapped reads averaged 91%.

Gene-level counts were generated with featureCounts v2.0.6 (Liao et al., 2014) in unstranded paired-end mode (-p --countReadPairs -B -C -s 0) against the *Sus scrofa* Ensembl annotation (Sscrofa11.1, release 109, 35,670 genes). On average 55% of read pairs were assigned to annotated genes (51 to 61% per sample).

### 2.5. Differential gene expression analysis

Differential expression analysis used DESeq2 v1.46.0 (Love et al., 2014) in R v4.4.3. Genes with 10 or more reads in at least 3 samples were retained, 21,961 genes in total (count matrix in Supplementary Table S3).

Five comparisons were made. The pooled genotype effect compared TgHD with WT across all animals, with age as a covariate (design ∼age_group + genotype). The genotype effect was tested separately at 4 years and at 8 years (design ∼genotype). WT ageing and TgHD ageing compared 8 years with 4 years in WT and in TgHD animals, respectively (design ∼age_group).

Normalisation, dispersion estimation and significance testing used default DESeq2 parameters, with the Wald test for all comparisons. Differentially expressed genes (DEGs) were identified at a false discovery rate (FDR) below 0.05 (Benjamini-Hochberg correction). The primary comparisons for downstream analyses were the genotype effects at 4 years and at 8 years.

### 2.6. Gene annotation

Gene annotations were retrieved from Ensembl BioMart (Sscrofa11.1) using the biomaRt R package v2.62.1 (Durinck et al., 2009). Symbols were assigned in order of confidence. Native pig symbols were available for 14,530 of the 21,961 tested genes. Where no pig symbol existed, we took the first token of the Ensembl description (132 genes), matched the description against human gene descriptions (395), and queried human orthologs directly, preferring one-to-one over one-to-many and many-to-many (196, 433 and 90, respectively). A further 58 genes were curated by hand. False-positive annotations were identified and reverted for non-protein-coding biotypes (lncRNA, pseudogenes). In total 15,834 genes have a symbol and the remaining 6,127 have none. The annotation source and confidence level were stored as a provenance tag for every gene (Supplementary Table S12). The 15,834 annotated genes correspond to 15,440 distinct symbols, 268 of which map to more than one Ensembl gene identifier. Where such a symbol is reported at the gene level, the entry with the highest mean normalised count was used. Ensembl identifiers for every gene are given in Supplementary Table S2.

### 2.7. Gene set enrichment analysis

Pre-ranked gene set enrichment analysis (GSEA) was performed with clusterProfiler v4.14.6 using the fgsea algorithm v1.32.4. RNA-seq genes were ranked by the DESeq2 Wald statistic and proteins by the limma v3.62.2 moderated t-statistic. Where a gene symbol appeared more than once, for example after protein-group expansion, the entry with the largest absolute ranking statistic was retained, one value per gene. Gene set size was restricted to 10 to 500 genes, or 5 to 500 genes for MitoCarta 3.0, and P values were corrected by the Benjamini-Hochberg method.

For the pig databases, GSEA was run directly on pig gene symbols, with org.Ss.eg.db for Gene Ontology (GO) and organism code ssc for KEGG. For the KEGG runs the symbols were first converted to Entrez identifiers by the bitr function of clusterProfiler. For the human databases, pig genes were mapped to hsapiens_homolog_associated_gene_name in the *Sus scrofa* Ensembl dataset using biomaRt (Durinck et al., 2009), and genes with no human ortholog were discarded. Where one pig gene had several human orthologs, the first one returned was kept, and where several pig genes shared one human symbol, their ranking values were averaged.

Seven pathway databases were used, KEGG, GO Biological Process, GO Cellular Component, Reactome, MSigDB Hallmark, WikiPathways and MitoCarta 3.0. KEGG and both GO domains were run on pig genes and on human orthologs, whereas Reactome, MSigDB Hallmark, WikiPathways and MitoCarta were run on human orthologs only. Supplementary Table S5. reports both the pig and the human results, with the number of gene sets tested per database and dataset. The figures show only the human ortholog results.

### 2.8. Cell-type marker gene sets

We compiled custom marker gene sets to test for changes in cell-type marker expression and in HD-relevant processes. Striatal cell-type markers were compiled from published single-cell and bulk transcriptomic studies (Anderson et al., 2023; Gokce et al., 2016; Jurga et al., 2020; Liddelow et al., 2017; Malaiya et al., 2021; Märtin et al., 2019; Matsushima et al., 2023; Saunders et al., 2018; Stanley et al., 2020) and the CellMarker 2.0 database (Hu et al., 2023). They cover medium spiny neurons with their D1, D2 and eccentric subtypes, the striosome and matrix compartments, cholinergic, parvalbumin-positive and somatostatin-positive interneurons, astrocytes with their reactive states, microglia with their homeostatic and disease-associated states, and oligodendrocytes. The reactive astrocyte sets, originally named A1, A2 and pan-reactive (Liddelow et al., 2017), are reported here as the inflammation-induced, the ischaemia-induced and the shared reactive set, following the consensus recommendation against binary astrocyte labels (Escartin et al., 2021). Microglial states are named following the consensus recommendations on microglial nomenclature (Paolicelli et al., 2022). Sets for the complement cascade, Toll-like receptor and NF-κB signalling, cytokines, chemokines and T cells were assembled from canonical pathway gene lists, and the disease-associated microglia set from Keren-Shaul et al. (2017) and Krasemann et al. (2017). Two functional sets were curated by hand, DNA repair (27 genes) (Handsaker et al., 2025) and heat-shock proteins and chaperones (23 genes) (Malla et al., 2021). One cross-species set contained the 23 genes reported as changed in the striatum of the OVT73 transgenic sheep model of HD (Handley et al., 2017).

Sources whose assignments conflicted for more than 80% of their genes were excluded. Genes assigned to several cell types within one study were removed, and conflicts between studies were resolved in favour of the cell type supported by the most independent sources. Each gene then received a composite score from citation frequency, marker specificity, validation confidence, expression in the pig putamen, and detection in the RNA-seq and proteomics data.

Each set contained the genes with a score of 8 or above, up to 60 genes per set. The marker database comprises 29 gene sets, 21 for cell types, subtypes, states and compartments, 5 immune pathway sets and 3 curated pathway sets, with 644 unique genes in total. These were tested with the GSEA function on TERM2GENE input, gene set size 5 to 500, and no P value cutoff. The complete marker database with provenance, scoring and per-set gene counts is provided in Supplementary Table S8.

### 2.9. Assignment of pathways to biological themes

To organise pathway results thematically, we assigned every pathway from the seven databases to a biological category by automated keyword matching. The curated vocabulary covers HD-relevant biology, such as synaptic transmission, mitochondrial metabolism, proteostasis, immune and inflammatory processes, myelination and neurodegeneration, alongside general categories such as signal transduction and the cell cycle.

Classification followed a fixed order, manual override rules first, then keyword scanning against the full vocabulary, with HD-relevant themes evaluated before general categories. Each pathway was assigned to the first matching theme, and pathways matching nothing were classified as Other. The complete pathway-to-theme mapping and the keyword definitions are provided in Supplementary Table S7.

### 2.10. Pathway selection and redundancy reduction for the figure panels

GSEA results are shown as dot heatmaps, one panel per theme (Section 2.9). More gene sets were significant than one panel can show, and many of them overlap, so the rows of each panel were selected in three steps.

First, redundant gene sets were removed. Gene sets with identical genes are shown once, under the shorter name. Gene Ontology is a hierarchy, so a broad term and a more specific term below it are often significant together. Within the biological process (GO:BP) and cellular component (GO:CC) ontologies, the most specific significant gene set was kept, and its broader parent terms were removed when their NES was concordant with that of the specific set, both positive or both negative. Gene sets from the other databases were not removed in this way, so a KEGG gene set and its Gene Ontology counterpart may both appear. Gene sets named in the Results were always kept.

Second, the remaining gene sets were ranked and the highest ranked were shown, 20 to 65 per panel depending on the theme. Gene sets were ranked by the number of datasets in which their FDR was below 0.05, and gene sets with equal counts were ordered by their mean minus log10 FDR in those datasets. Far more gene sets were significant in the 8-year transcriptome than in either proteomic comparison, so this ranking alone would have excluded gene sets significant only in the proteome. In the thematic panels, the count of every gene set with an FDR below 0.05 in either proteomic comparison was therefore increased by 4. These gene sets are ranked above all others, and the other rows of the panel follow in the order of the ranking.

Third, the gene sets shown in each panel were grouped by the genes they share, so that gene sets with overlapping genes are next to each other. The similarity of every pair was calculated as the mean of the Jaccard index and the overlap coefficient (Merico et al., 2010). Gene sets were then clustered by average linkage, and the tree was cut at a similarity of 0.375 (Reimand et al., 2019). The genes of each set were those used in the enrichment analysis. All significant gene sets, including those not shown in the panels, are listed in Supplementary Table S5.

The KEGG overview in Fig. 2C was selected differently. KEGG contains fewer and less nested gene sets than the other databases, so it was used for a compact overview. The panel shows the 20 pathways with the highest absolute NES in any one dataset among the 57 KEGG pathways significant in at least two of the four datasets, and the seven KEGG neurodegenerative disease pathways are shown separately in Fig. 2D.

### 2.11. Alternative splicing analysis

Alternative splicing was quantified with rMATS-turbo v4.3.0 (Wang et al., 2024) on the STAR-aligned BAM files, using the *Sus scrofa* gene annotation (Sscrofa11.1, Ensembl release 109) and a read length of 150 bp. Novel splice site detection was enabled, so events absent from the annotation were also tested. Five event types were quantified, skipped exon (SE), alternative 5’ splice site (A5SS), alternative 3’ splice site (A3SS), mutually exclusive exon (MXE) and retained intron (RI).

Four pairwise comparisons were analysed, each with three animals per group. These were the genotype effect at 4 years (WT against TgHD), the genotype effect at 8 years (WT against TgHD), TgHD ageing (TgHD at 4 against 8 years) and WT ageing (WT at 4 against 8 years). In every comparison the wild-type or younger group was the first group, so a positive inclusion level difference indicates higher inclusion in that group and reduced inclusion in the transgenic or older group. Both junction count (JC) and junction count plus exon count (JCEC) results were generated and are reported in Supplementary Table S9, and only the JC results were used for downstream analyses.

Significant events were defined at an FDR of 0.05 or below, with no minimum threshold on the inclusion level (PSI) difference. Events were then merged with the RNA-seq differential expression results by gene symbol, and events in genes with low expression (baseMean below 20) or without a gene symbol were excluded. Event counts before and after each step, for every comparison and event type, are given in Supplementary Table S9.

### 2.12. Positional RNA-binding-protein motif analysis

We asked whether the exons that change inclusion contain RNA-binding protein (RBP) motifs at particular positions relative to their splice sites. Window geometry and the choice of comparison exons follow the RNA maps of Park et al. (2016) and Hwang et al. (2020). Skipped-exon events with an FDR below 0.05 and an absolute inclusion difference above 0.10 were split into those with reduced and those with increased inclusion in TgHD. Each group was compared with unchanged events whose inclusion level was between 0.15 and 0.85, capped at 4,000. Four windows were defined per event, one at each splice site, each with 50 nucleotides of exon and 250 nucleotides of the adjacent intron. The four sites are the 5’ splice site of the upstream exon, the 3’ and 5’ splice sites of the skipped exon and the 3’ splice site of the downstream exon. Duplicate windows were removed within each position, because rMATS lists the same target exon once for each pair of flanking exons.

Sequence was scanned with two motif collections. The first was the human ATtRACT database (Giudice et al., 2016). It lists several matrices for many proteins and does not rank them, so one matrix per protein was retained by a fixed rule based on experiment type, motif length and agreement among that protein’s other entries. In all, 146 distinct matrices across 159 proteins were used. The second was the *Sus scrofa* set of CisBP-RNA (Rosado-Tristani et al., 2025), which lists one motif per protein and needed no such rule, with 171 matrices across 193 proteins. No motif in either collection was measured on pig protein, so conservation of binding preference is assumed.

Motif matches were counted in a 50-nucleotide window sliding in 1-nucleotide steps, which is the profile the RNA maps plot. For testing, each window was divided instead into six non-overlapping positions of 50 nucleotides, one exonic and five intronic. The exonic positions were not tested. A fixed 50-nucleotide position on a short exon includes intronic sequence, and exons with reduced inclusion are shorter than comparison exons (median 74 against 105 nucleotides, Wilcoxon P below 2e-16), so those positions would not be comparable between the groups. The intronic positions were tested even where the 250-nucleotide arm extends into a neighbouring exon, because flanking intron length did not differ between the two groups (Wilcoxon P = 0.094).

Each position was compared between a changed group and the comparison set by Fisher’s exact test, and positions where fewer than ten events contained the motif were not tested. P values were corrected by the Benjamini-Hochberg method within each comparison and each database, and results are reported at an FDR below 0.05. This differs from rMAPS, which applies a Wilcoxon rank sum test to each sliding window and reports uncorrected P values.

Matrices were grouped into families by Pearson correlation of their best ungapped alignment, joined at an r of 0.85 or above, into 76 families for ATtRACT and 56 for CisBP-RNA. Proteins in one family are reported as a single result. A consensus of two or fewer distinct bases, or one base at 75% or more, counts as low complexity.

### 2.13. Over-representation analysis of spliced genes

Genes with at least one significant alternative splicing event (Section 2.11) were tested for pathway over-representation, separately for the genotype comparisons at 4 and 8 years and for TgHD ageing. Pig gene symbols were matched to human Entrez identifiers through org.Hs.eg.db. Over-representation was tested with clusterProfiler, using enrichGO for Gene Ontology Biological Process in all three comparisons and enrichKEGG for human KEGG pathways in the 8-year and TgHD ageing comparisons. P values were corrected by the Benjamini-Hochberg method, and terms with an adjusted P value below 0.05 and a q value below 0.1 are reported in Supplementary Table S16.

### 2.14. Validation

#### 2.14.1. qPCR validation

Quantitative PCR (qPCR) was used to validate the RNA-seq results in two cohorts. The first was the sequenced 8-year cohort (3 TgHD and 3 WT), from which RNA was isolated anew, with several independent isolations per animal. RNA (500 ng) was reverse transcribed with the High-Capacity cDNA Reverse Transcription Kit (Thermo Fisher Scientific, 4368814).

Reactions were run on a CFX96 Touch Real-Time PCR Detection System (Bio-Rad) with SYBR Green (Thermo Fisher Scientific, A25780) and primers designed in-house (Supplementary Table S17). Expression was normalised to *ACTB* by the ΔCt method and compared between genotypes as 2^(−ΔΔCt). *ACTB* was chosen because it did not differ between genotypes in the RNA-seq data, with a log2 fold change of −0.010 at a mean normalised count of 63,695 in the 8-year comparison (Supplementary Table S2).

The second cohort, 6 TgHD and 5 WT animals at 7 years, had not been sequenced and was processed by the same protocol with a partly different gene panel. Fourteen genes were analysed in the 8-year cohort and 21 in the 7-year cohort (Supplementary Table S11). For each cohort, the log2 fold change between TgHD and WT from qPCR was correlated with that from the 8-year RNA-seq comparison by Pearson correlation.

#### 2.14.2. RNA-protein concordance

Agreement between the transcriptome and the proteome was assessed in three ways. First, across individual genes, DESeq2 log2 fold changes from the 8-year comparison were correlated with limma log2 fold changes from the 7- to 8-year proteomic comparison, for the genes detected in both datasets. Second, within each of the 12 animals that had both RNA-seq and proteomic data, variance-stabilised RNA counts were correlated with log2 protein abundances across the genes detected in both. Third, across pathways, GSEA normalised enrichment scores (NES) from the human ortholog KEGG, Gene Ontology and Reactome results were correlated (Pearson) between the 8-year transcriptome and the 7- to 8-year proteome for all pathways tested in both datasets, and the proportion of concordant pathways, positively enriched in both or negatively enriched in both, was counted among the pathways significant in both datasets (FDR below 0.05). The first two comparisons were Pearson correlations as well, and the underlying values are given in Supplementary Table S11.

### 2.15. DNA damage assay and mtDNA copy number estimation

DNA was isolated from frozen cortex and putamen of the 7-year cohort (5 WT and 7 TgHD) with the DNeasy Blood and Tissue Kit (Qiagen, 69506) according to the manufacturer’s protocol. The eluted DNA was quantified on a NanoDrop 1000 (Thermo Fisher Scientific), and the concentration was adjusted to 10 ng/µL for nuclear DNA (nDNA) and 2 ng/µL for mitochondrial DNA (mtDNA) integrity measurements. DNA integrity was measured by real-time qPCR analysis of damage frequency (RADF) (Wang et al., 2016). The method is based on the ability of a lesion on one strand to inhibit digestion of double-stranded DNA by the restriction enzyme TaqI (New England Biolabs, R0149L). TaqI recognises the palindromic sequence 5’-TCGA-3’ and cuts between T and C, leaving a 5’ overhang of two nucleotides. According to the protocol, 30 ng of nDNA or 6 ng of mtDNA template was added to reactions with and without TaqI. The DNA damage level was calculated as 2^−(CtTaq − Ctnt), where CtTaq is the threshold cycle of the sample with TaqI and Ctnt that of the sample without TaqI. Primers for *MT-RNR1* (product length 108 bp) and *NDUFAS* (product length 148 bp) were used to quantify mtDNA and nDNA damage, respectively (Supplementary Table S17). The mtDNA copy number relative to nDNA was calculated from the real-time PCR data as 2^(CtnDNA − CtmtDNA).

### 2.16. Protein extraction

Approximately 60 mg of putamen tissue per sample was mixed 1:3 (w/v) with PBS (4 °C) containing 1× HALT protease and phosphatase inhibitor (Thermo Fisher Scientific) and homogenised with a stainless-steel ball in a TissueLyser II (Qiagen) at 30 Hz for 2 min. Sodium dodecyl sulphate (SDS) in PBS was added to a final concentration of 1% (w/v). Samples were vortexed for 15 s and homogenised with the immersed 2 mm miniprobe of a Q500 sonicator (Qsonica) three times at 20% power for 10 s at room temperature.

Homogenates were diluted 10-fold with 1% SDS in PBS containing 1× HALT inhibitor and boiled at 95 °C for 5 min. Sample lysis continued for 10 cycles of centrifugation at 2,350 × *g* for 15 s followed by vortexing for 20 s in a Multi-Spin MSC-6000 (Biosan) at room temperature. Lysates were centrifuged (20,000 × *g*, 15 min, 20 °C), and supernatants were collected and stored at −80 °C. Protein concentrations were determined using the Pierce BCA Protein Assay Kit (Thermo Fisher Scientific) according to the manufacturer’s instructions.

### 2.17. Sample preparation for mass spectrometry

Proteins were prepared by a modified filter-aided sample preparation (FASP) protocol as we described previously (Vymola et al., 2024). Briefly, 30 µg of protein per sample was loaded onto Microcon 30K centrifugal ultrafiltration units (Merck) and washed three times with wash buffer (8 M urea and 5 mM EDTA in 50 mM ammonium bicarbonate) by centrifugation at 12,000 × *g* for 15 min at 22 °C. Proteins were reduced with 10 mM tris(2-carboxyethyl)phosphine in wash buffer (60 min, 32 °C) and alkylated with 40 mM iodoacetamide in wash buffer (30 min, 25 °C, in the dark). Ultrafiltration units were washed once with wash buffer and three times with 50 mM ammonium bicarbonate. On-filter digestion was performed with 1 µg of rLys-C protease (Promega) in 50 mM ammonium bicarbonate containing 0.02% (w/v) ProteaseMAX (Promega) for 80 min at 37 °C, followed by digestion with 1 µg of Trypsin Platinum protease (Promega) for 12 h at 37 °C. Peptides were collected by centrifugation, acidified with formic acid to 2% (v/v) and centrifuged (14,000 × *g*, 20 min, 4 °C).

Supernatants were desalted using Pierce peptide desalting spin columns (Thermo Fisher Scientific) according to the manufacturer’s instructions, except that formic acid replaced trifluoroacetic acid. Peptides were dried by vacuum centrifugation at 45 °C and resuspended in 0.1% formic acid, and their concentrations were measured at 280 nm (Synergy HTX, BioTek). Samples were diluted to 0.2 µg/µL in QuanRecovery with MaxPeak HPS vials (Waters) and spiked with indexed retention time (iRT) peptides (Biognosys) at an iRT to sample ratio of 1:30 (v/v).

### 2.18. Liquid chromatography-mass spectrometry

A liquid chromatography-tandem mass spectrometry (LC-MS/MS) system consisting of a nanoElute 2 coupled to a timsTOF HT (both Bruker Daltonics) was operated as described previously (Výmolová et al., 2026). Briefly, peptides were loaded onto a trap column (PepMap Neo C18, 5 × 0.3 mm, 5 µm, Thermo Fisher Scientific) and then separated on an analytical column (PepSep Ultra C18, 25 cm × 75 µm, 1.5 µm, Bruker Daltonics) using a linear gradient of 2 to 35% acetonitrile with 0.1% formic acid over 90 min at 200 nL/min and 50 °C.

For data-dependent acquisition (DDA) with parallel accumulation-serial fragmentation (PASEF), 0.6 µg of peptides per sample was analysed with the default DDA PASEF-standard_1.1_sec_cycletime method (1/K_0_ range 0.6 to 1.6 V s/cm^2^, ramp/accumulation time 100 ms, 10 PASEF ramps per cycle, total cycle time 1.17 s) in positive ion mode with an MS scan range of 300 to 1700 *m/z*.

For data-independent acquisition (DIA), 0.8 µg of peptides per sample was analysed using the default dia-PASEF long gradient method (1/K_0_ range 0.6 to 1.6 V s/cm^2^, ramp/accumulation time 100 ms, 16 MS/MS ramps, 32 MS/MS windows, cycle time 1.8 s) in positive ion mode with an MS scan range of 300 to 1700 *m/z*.

### 2.19. Mass spectrometry data processing

All raw data were analysed in Spectronaut (v19.5.241126.62635, Biognosys) as we described previously (Cervenka et al., 2026), and the protein group quantities used here were subsequently exported with Spectronaut v21.0.260703.94842 (Section 2.20). DIA data were processed with the directDIA workflow and the default BGS Factory Settings, with the following modifications. The corresponding DDA data were used as library extension runs to create a hybrid spectral library. The protein FASTA database was compiled by merging the UniProtKB *Sus scrofa* reference proteome (UP000008227, release 2024_06, 22,833 proteins, one sequence per gene) (UniProt Consortium, 2025) with the chicken protein OVAL (P01012), yeast ADH1 (P00330), the human N-terminal HTT fragment (amino acids 1 to 548, P42858), 381 universal contaminant proteins (Frankenfield et al., 2022) and the iRT sequence (Biognosys).

A custom cleavage rule was applied for the combination of Lys-C and trypsin, with cleavage after every lysine and after every arginine not followed by proline, and up to two missed cleavages allowed. Carbamidomethylation of cysteine was set as a fixed modification, and protein N-terminal acetylation and methionine oxidation as variable modifications. The Hybrid (DDA + DIA) Library option was selected in the DIA Analysis Workflow. Data were normalised using Spectronaut’s cross-run local normalisation with no filter type and automatic row selection. Proteins identified by a single stripped peptide sequence were excluded.

### 2.20. Statistical analysis of proteomics data

Protein group quantities were normalised in Spectronaut (Section 2.19) and exported in two forms, one with all 27 samples processed and normalised together, and one with each age group processed and normalised separately (4 TgHD and 4 WT at 4 years, 10 TgHD and 9 WT at 7 to 8 years). The joint export contained 8,216 protein groups. The age-separated export was the primary dataset and was used for every genotype comparison.

Both exports were processed in R in the same way. Protein groups made up only of contaminants or of proteins of a species other than *Sus scrofa* were removed; where a pig protein shared a group with a contaminant, only the contaminant was removed. The human HTT fragment (P42858) was exported separately to confirm genotype and then removed, because it is present only in TgHD animals. As a protein group can list several accessions, each such group was expanded to one entry per gene, and every entry was assigned the quantity of its group.

Quantities were log2-transformed, and zero values were set to missing without imputation. Entries quantified in at least 50% of the samples in at least one age-by-genotype group were retained, 8,496 in the age-separated dataset, 8,168 of them with a gene symbol. Differential abundance was tested with limma v3.62.2 by moderated t-tests, with each entry fitted on the samples in which it was quantified, and P values were corrected by the Benjamini-Hochberg method. TgHD was compared with WT at 4 years and in the 7- to 8-year group. The two ages were compared within each genotype in the joint export only, because quantities normalised separately for each age cannot be compared between ages.

The proteomic data were interpreted through GSEA, run on the limma moderated t-statistic with the same databases and settings as for the RNA-seq data (Section 2.7).

### 2.21. Transcription factor enrichment analysis

Transcription factors (TFs) whose known targets were overrepresented among the DEGs were ranked with ChEA3 (Keenan et al., 2019). The genes upregulated and the genes downregulated in TgHD at 8 years were submitted as two separate lists of gene symbols to the ChEA3 web server (https://maayanlab.cloud/chea3/). The target sets are defined for human genes, so pig gene symbols were used as human symbols, which assumes that TF targets are conserved. The 20 highest-ranked TFs for each list are given in Supplementary Table S13.

### 2.22. Software

Statistical analysis and plotting were performed in R v4.4.3 with Bioconductor v3.20. The main packages were DESeq2 v1.46.0, limma v3.62.2, clusterProfiler v4.14.6, fgsea v1.32.4, ReactomePA v1.50.0, msigdbr v25.1.1, biomaRt v2.62.1, org.Hs.eg.db and org.Ss.eg.db v3.20.0, Biostrings v2.74.1, ComplexHeatmap v2.22.0 and ggplot2 v4.0.1. All R package versions are listed in Supplementary Table S14. Tools run outside R, including rMATS-turbo v4.3.0 and Spectronaut, are described in the sections that use them. Figures were assembled in Canva Pro.

## 3. Results

### 3.1. Experimental design, validation and overview of the molecular changes

#### 3.1.1. The TgHD Libechov minipig and experimental design

We characterised the putamen of the TgHD Libechov minipig at two ages by bulk RNA sequencing and label-free mass spectrometry proteomics. RNA-seq was performed on putamen tissue from TgHD and WT minipigs at 4 and 8 years of age (n = 3 per genotype and age, Fig. 1), approximately one-third and two-thirds of the expected lifespan of this breed. Label-free proteomics was conducted on a larger, partially overlapping cohort of 27 animals (4 WT and 4 TgHD at 4 years, 9 WT and 10 TgHD at 7 to 8 years, Fig. 1).

**Figure 1.**
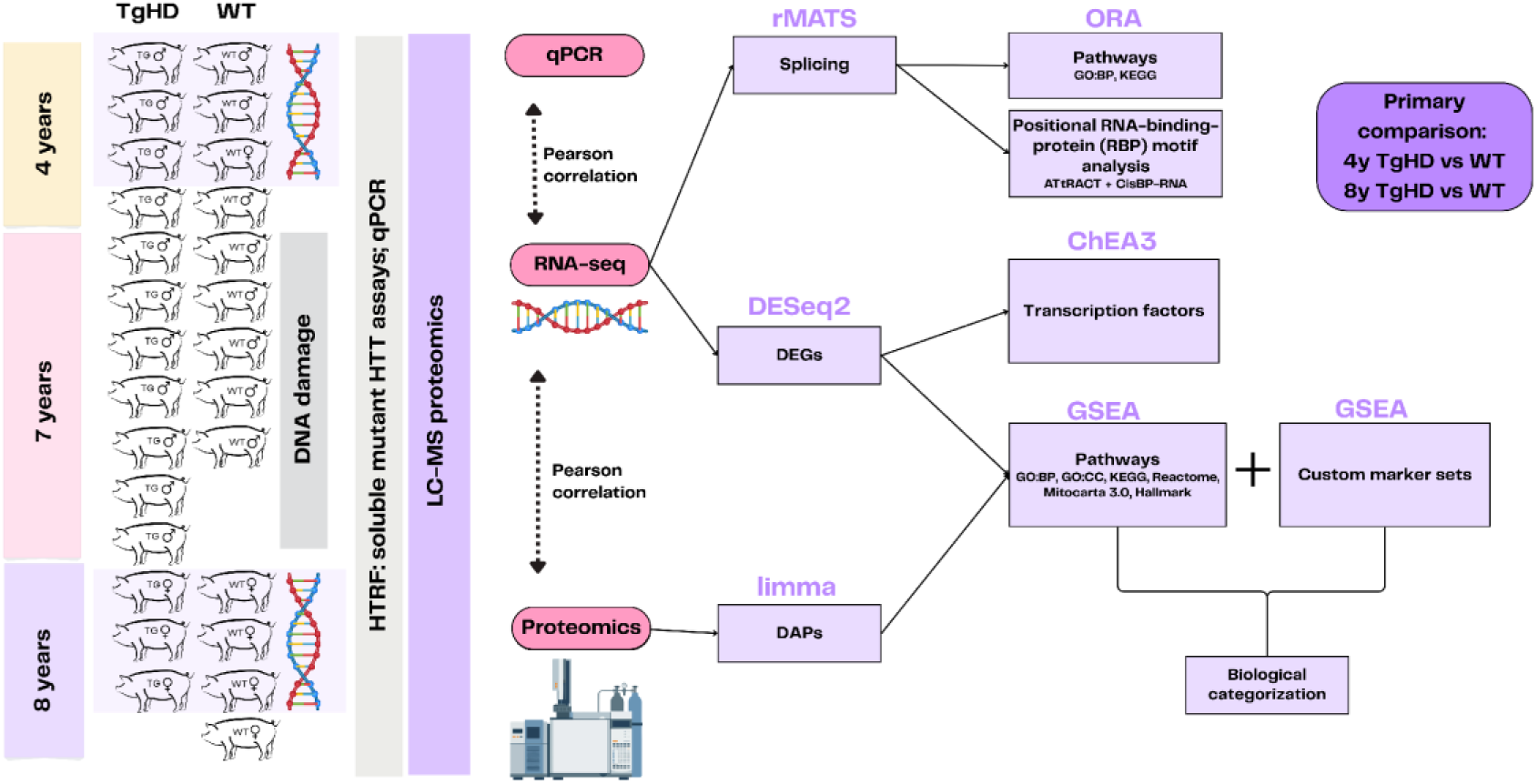
Study design. Schematic of the animals, tissues and analyses. Bulk RNA sequencing was performed on putamen tissue from TgHD and WT Libechov minipigs at 4 and 8 years of age (n = 3 per genotype and age). Label-free mass spectrometry proteomics was performed on a larger, partially overlapping cohort at 4 years (n = 4 per genotype) and at 7 to 8 years (n = 10 TgHD, n = 9 WT). All datasets were analysed for pathway enrichment and for cell-type marker enrichment by GSEA. Alternative splicing was quantified from the RNA sequencing data with rMATS, and the differentially spliced genes were tested by ORA for pathway enrichment. Differentially spliced genes were also tested by positional RBP motif analysis. Soluble mutant huntingtin was measured by HTRF in cortex and putamen. Cortex and putamen of the 7-year cohort (Methods 2.1) were also tested for DNA damage by RADF. The full animal overview is in Supplementary Table S1. Abbreviations. DAPs, differentially abundant proteins. DEGs, differentially expressed genes. GSEA, gene set enrichment analysis. HTRF, homogeneous time-resolved fluorescence. LC-MS, liquid chromatography mass spectrometry. ORA, over-representation analysis. RADF, real-time qPCR analysis of damage frequency. RBP, RNA-binding protein. TgHD, transgenic Huntington’s disease minipig. WT, wild type.

**Figure 2.**
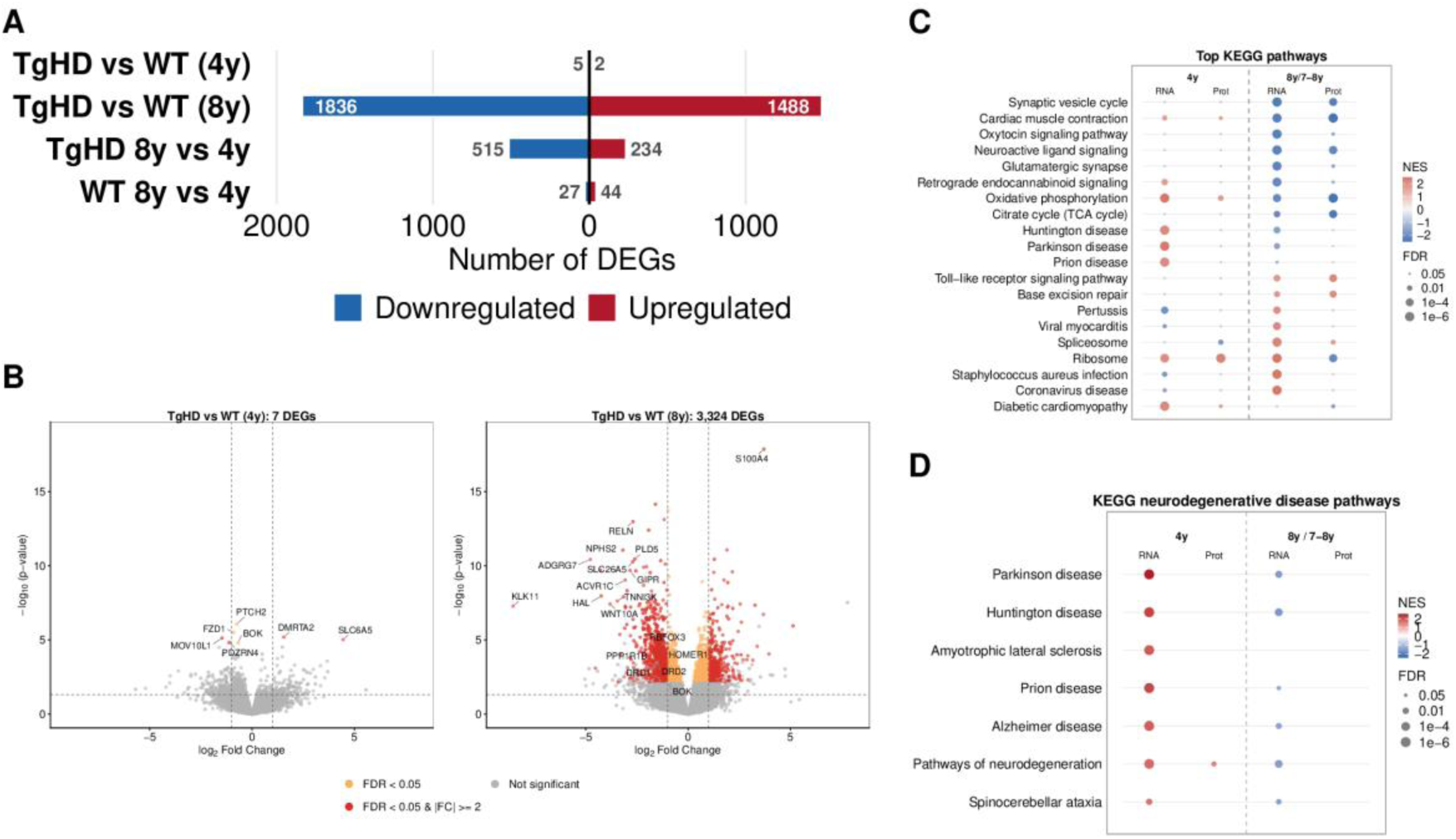
Transcriptomic and proteomic changes in the TgHD minipig putamen at 4 and 8 years. **(A)** Counts of DEGs in four comparisons. Bars extend left for genes lower and right for genes higher in the first-named group of each comparison. **(B)** Volcano plots of TgHD versus WT at 4 years (left) and 8 years (right). Up to ten genes per panel are labelled, those that are annotated in the genome and have the largest absolute log2 fold change multiplied by minus log10 nominal P, together with a curated set of genes with established roles in striatal identity and Huntington disease, labelled at 8 years only. **(C)** Dot heatmap of GSEA NES of KEGG pathways. The 20 pathways shown are those with the highest single-dataset absolute NES among the 57 KEGG pathways significant in at least two of the four datasets. The cardiac pathways appear because their gene sets contain calcium-handling and dystrophin-glycoprotein complex genes expressed in the brain (*CACNA1C*, *CACNG2*, *RYR2*, *ATP2A2*, *SLC8A1* and *DMD*), several of which encode synaptic proteins. **(D)** Dot heatmap of GSEA NES of the seven KEGG neurodegenerative disease pathways. In (C) and (D), columns are RNA-seq (RNA) and age-normalised proteomics (Prot), the later age being 8 years for RNA-seq and 7 to 8 years for proteomics (8y/7-8y), and all comparisons are TgHD relative to WT. Grey dots mark comparisons not reaching an FDR below 0.05. The full pathway list is in Supplementary Table S5. Abbreviations. DEG, differentially expressed gene. FDR, false discovery rate. GSEA, gene set enrichment analysis. NES, normalised enrichment score. TgHD, transgenic Huntington’s disease minipig. WT, wild type.

#### 3.1.2. Transcriptomic and proteomic changes in TgHD minipigs at 4 and 8 years

The exploratory principal component analysis (PCA) of transcriptomic data clearly separated TgHD and WT animals at 8 years, whereas at 4 years the two genotypes were intermixed (Supplementary Fig. S1A-C). In the proteome the first principal component separated the two age groups, as expected for data normalised separately for each age, and the genotypes were intermixed within each age group (Supplementary Fig. S1D). The differential expression analysis (DESeq2, FDR below 0.05) identified only 7 DEGs at 4 years (TgHD against WT), compared with 3,324 DEGs at 8 years (1,488 upregulated, 1,836 downregulated, Fig. 2A). Far fewer genes changed with age alone. WT animals differed by 71 genes between 4 and 8 years (44 up, 27 down), against 749 in TgHD animals (234 up, 515 down, Supplementary Table S2).

Several genes significantly downregulated in TgHD (Fig. 2B), including the dopamine receptors *DRD1* and *DRD2*, the MSN marker *PPP1R1B* (DARPP-32), the synaptic scaffold *HOMER1* and the extracellular matrix glycoprotein *RELN*, have established roles in striatal neuron identity and HD pathophysiology (Bates et al., 2015; Malla et al., 2021; Matsushima et al., 2023). *BOK*, a pro-apoptotic BCL-2 family member, was the only gene significant at both 4 and 8 years in the TgHD against WT comparison (log2 fold change −0.68 and −0.89), but it has not been linked to HD in the literature. The complete list of DEGs and the volcano plots for all age and genotype comparisons are given in Supplementary Table S2 and Supplementary Fig. S2.

Despite the significant transcriptomic changes in the TgHD putamen, no individual protein differed between the genotypes at an FDR below 0.05 at either age in the proteome (135 proteins at 4 years and 160 at 7 to 8 years had an unadjusted P below 0.05, Supplementary Fig. S3, Supplementary Table S4).

#### 3.1.3. qPCR validation and agreement between the datasets

To test technical stability, we quantified 14 transcripts by qPCR on RNA re-isolated from the same 8-year-old animals. The qPCR fold changes agreed with the RNA-seq fold changes (Pearson r = 0.89, 13 of 14 genes concordant in direction, Supplementary Fig. S4A). To test whether the findings extend to animals outside the sequenced cohort, we measured 21 genes in an independent cohort of 7-year-old animals (n = 6 TgHD and n = 5 WT). Their fold changes were moderately concordant with the 8-year RNA-seq data (r = 0.65, P = 0.0015, 13 of 21 genes concordant in direction, Supplementary Fig. S4B).

Next we examined the agreement between the RNA sequencing and the mass spectrometry data. Across the 12 animals with matched transcriptomic and proteomic data, transcript and protein abundances of the genes quantified in both datasets were weakly positively correlated within each animal (mean Pearson r = 0.41, SD = 0.02, Supplementary Table S11). We then correlated the pathway analyses of the transcriptome and the proteome. The GSEA NES of the 8-year transcriptome and the 7- to 8-year proteome correlated at r = 0.46 across the 6,253 human ortholog KEGG, GO and Reactome pathways tested in both datasets. Of the 337 pathways significant in both, 310 (92%) were concordant, positively enriched in both datasets or negatively enriched in both (Supplementary Fig. S4C, Supplementary Table S11). The 27 discordant pathways formed two groups, 20 cilium and flagellum gene sets with negative NES in the transcriptome and positive NES in the proteome, and seven with the opposite pattern, five of them ribosome and translation gene sets.

#### 3.1.4. Soluble mutant huntingtin

Putamen and cortical tissues from the whole cohort were analysed by HTRF. Using assays specific for soluble mHTT, the N548-124Q transgenic protein was readily detected in TgHD but not in WT animals. We detected no consistent age-dependent accumulation in either putamen or cortex, although single comparisons reached significance (Supplementary Fig. S5A and B, Supplementary Table S18).

#### 3.1.5. Pathway enrichment overview and neurodegeneration pathways

For an overview of the pathways that differ between TgHD and WT animals, we applied GSEA to the transcriptome and the proteome at each age, with the same pathway databases for both.

GSEA of the 8-year transcriptome identified hundreds of significantly enriched gene sets in every database, 1,094 in GO biological process alone (Supplementary Table S5). A positive NES means that the genes of a set are enriched among the genes with higher expression or protein abundance in TgHD than in WT animals of the same age, and a negative NES means they are enriched among genes with lower expression or abundance. Fig. 2C gives a compact overview of the KEGG pathways, which are fewer and less nested than those of the other databases (47 significant at 4 years and 136 at 8 years in the transcriptome), and Fig. 2D shows the KEGG neurodegenerative disease pathways separately.

Seven KEGG neurodegenerative disease pathways, including the Huntington disease pathway, were significantly and positively enriched in TgHD relative to WT in the transcriptome at 4 years (NES +1.64 to +2.48, FDR below 0.05), and Parkinson disease had the highest NES (Fig. 2D). In the proteome at 4 years, only one of them, pathways of neurodegeneration (hsa05022), was significant (NES = +1.50, FDR = 0.030). At 8 years six of these seven pathways were significantly but negatively enriched in the transcriptome, including the Huntington disease pathway (NES = −1.64, FDR = 0.0010). No neurodegenerative disease pathway was significant in the proteome at 7 to 8 years.

To identify the genes that contribute to the enrichment of the Huntington disease pathway, we mapped gene-level TgHD versus WT fold changes onto the KEGG Huntington disease pathway (hsa05016, Supplementary Fig. S6A-C) with the Pathview package (Luo and Brouwer, 2013). At 4 years the genes with higher expression in TgHD were mostly genes encoding oxidative phosphorylation (OXPHOS) complex subunits, while at 8 years the genes with lower expression included both OXPHOS subunit genes and neuronal genes. The seven neurodegenerative disease pathways share many of these genes (Supplementary Table S15), which explains why disease pathways other than Huntington disease are also enriched in this model.

Among the 20 pathways in Fig. 2C, the most negatively enriched pathway in TgHD against WT at 8 years in the transcriptome was the synaptic vesicle cycle (NES = −2.37), followed closely by cardiac muscle contraction, oxytocin signalling, neuroactive ligand signalling and the glutamatergic synapse (NES −2.20 to −2.29). The cardiac pathways are negatively enriched because their gene sets are built from calcium-handling and dystrophin-glycoprotein complex genes expressed in the brain, several of which are synaptic.

The ribosome pathway (KEGG) was positively enriched in the transcriptome at both ages (NES = +2.10 at 4 years and +2.75 at 8 years), while in the proteome it was positively enriched at 4 years (NES = +2.28) and negatively enriched at 7 to 8 years (NES = −2.10). Ribosomal protein transcripts were therefore higher and ribosomal proteins lower in TgHD at the older age. Given the changes in synaptic, mitochondrial, immune, splicing and DNA-repair pathways in this overview, we examined each of these in more detail in the following sections.

### 3.2. Negative enrichment of neuronal marker sets and synaptic pathways at 8 years

#### 3.2.1. Cell-type marker set enrichment in the TgHD putamen

In human HD striatum, single-nucleus transcriptomics has confirmed that indirect-pathway medium spiny neurons, which express D2 receptors (D2 MSN), are the most affected cell type, followed by direct-pathway D1 neurons (D1 MSN) (Lee et al., 2020). To determine which striatal cell-type marker sets change in our bulk data, we curated neuronal (D1 MSN, D2 MSN, pan-MSN and interneuron) and glial (oligodendrocyte, microglia, disease-associated microglia, astrocyte and reactive astrocyte) marker gene sets (Methods 2.8, Supplementary Table S8). We tested them by GSEA in the TgHD versus WT comparison at 4 and 8 years, in both the transcriptome and the proteome (Fig. 3A), and in the 8-year versus 4-year comparison within each genotype (Supplementary Table S6).

**Figure 3.**
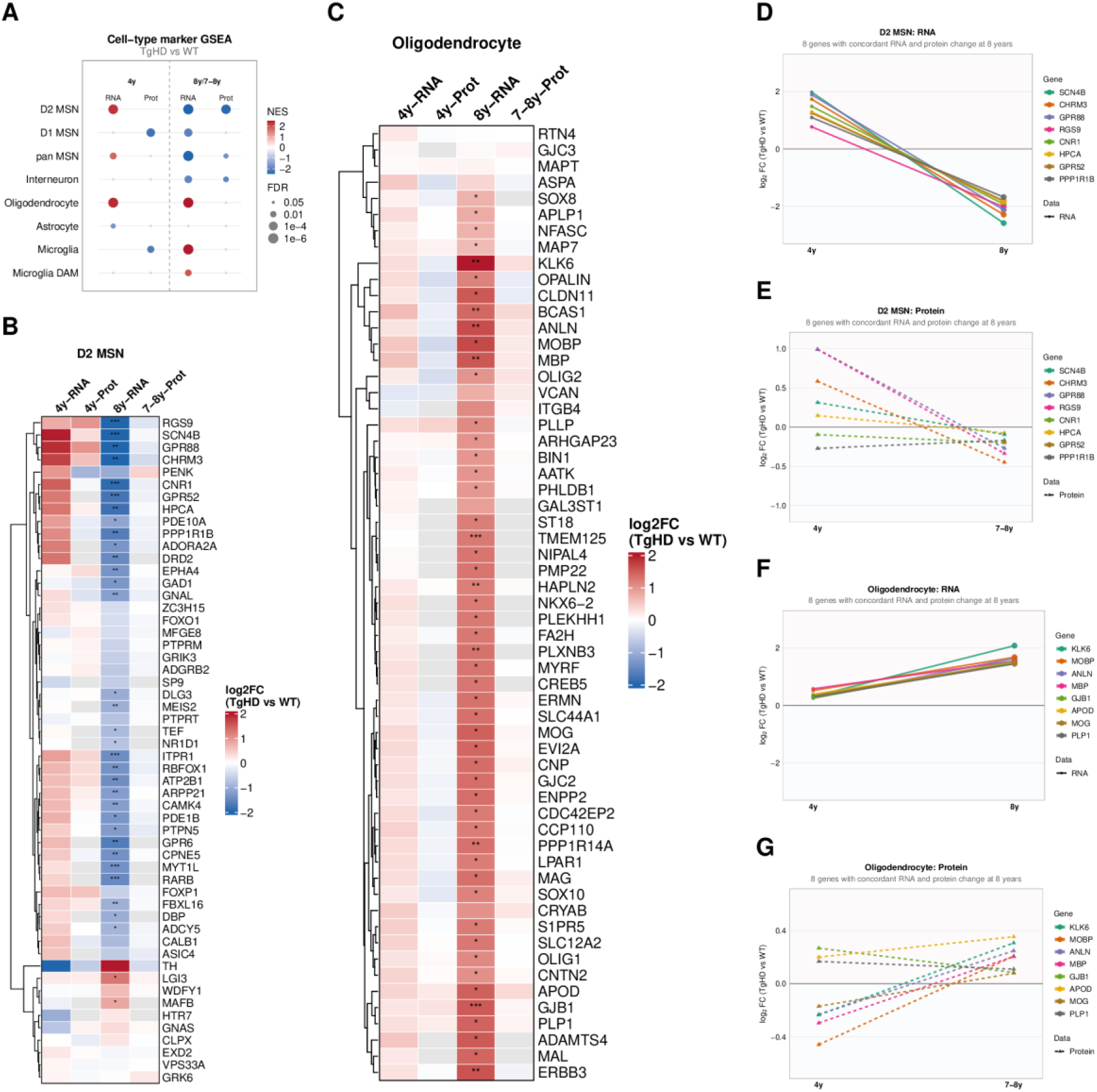
Cell-type marker enrichment and changes in single candidate genes between 4 and 8 years. **(A)** Dot heatmap of GSEA NES of curated cell-type marker gene sets. **(B, C)** Heatmaps of log2 fold change of the single genes used as markers in the GSEA, for the D2 medium spiny neuron (D2 MSN) marker set (B) and the oligodendrocyte marker set (C), with rows clustered by Euclidean distance. Grey cells mark genes not detected in that dataset. Asterisks mark RNA-seq FDR significance, * below 0.05, ** below 0.01, *** below 0.001. **(D-G)** Log2 fold change of TgHD relative to WT at 4 and at 8 years for the candidate genes that change concordantly in RNA and protein at 8 years. Eight genes are shown per panel, those with the largest root of the summed squared log2 fold changes in RNA-seq and proteomics, after requiring an absolute log2 fold change of at least 0.3 in one of them. D2 MSN set in RNA-seq (D) and proteomics (E), oligodendrocyte set in RNA-seq (F) and proteomics (G). Large points mark an FDR below 0.05 in the RNA-seq data. Throughout, columns are RNA-seq (RNA) and age-normalised proteomics (Prot), the later age being 8 years for RNA-seq and 7 to 8 years for proteomics (8y/7-8y), and all comparisons are TgHD relative to WT. Grey dots in (A) mark comparisons not reaching an FDR below 0.05. Marker gene sets and their provenance are in Supplementary Table S8. Abbreviations. D2 MSN, dopamine receptor D2 medium spiny neuron. FDR, false discovery rate. GSEA, gene set enrichment analysis. NES, normalised enrichment score. TgHD, transgenic Huntington’s disease minipig. WT, wild type.

The D2 MSN marker set was negatively enriched in both the transcriptome at 8 years and the proteome at 7 to 8 years (NES = −3.02, FDR = 8.6e-10 and NES = −2.45, FDR = 1.3e-05, respectively, Supplementary Table S6), consistent with reduced expression of D2 MSN identity genes. Of the 54 genes in the set, 34 were significant DEGs at 8 years, 32 of them downregulated in TgHD (Fig. 3B). Individual marker proteins had smaller log2 fold changes than their transcripts, yet the set as a whole was significantly negatively enriched (Fig. 3A).

Unexpectedly, the D2 MSN marker set was positively enriched in the transcriptome at 4 years (NES = +2.30, FDR = 6.9e-06, Fig. 3A, Supplementary Table S6), even though no individual gene was a significant DEG (Fig. 3B). The change between the two ages was confirmed in the direct comparison within TgHD animals (8 years versus 4 years, NES = −2.92, FDR = 9.4e-10). For single genes such as *SCN4B*, *CHRM3* and *GPR88*, the log2 fold change differed by 4.0 to 4.6 units between the two ages, from positive to negative, which corresponds to a 16- to 23-fold difference in the TgHD to WT ratio.

The D1 MSN marker set was also negatively enriched in TgHD relative to WT in the transcriptome at 8 years (NES = −2.00, FDR = 0.00034, Supplementary Table S6), but not in the 7- to 8-year proteome (NES = −0.71, FDR = 0.92). At 4 years the D1 and D2 MSN sets differed. While the D2 MSN set was positively enriched in the transcriptome and positive but not significant in the proteome (NES = +1.44), the D1 MSN set was significantly negatively enriched in the proteome (NES = −2.27, FDR = 0.00018) and negative but not significant in the transcriptome (NES = −1.30). The pan-MSN marker set was concordant with the D2 MSN set in every dataset (NES = +1.78 and −3.10 in the transcriptome at 4 and 8 years, respectively, and −1.78 in the 7- to 8-year proteome).

Interneurons are generally described as relatively spared in HD (Donaldson et al., 2026), yet in the TgHD model at 8 years the interneuron marker set was significantly negatively enriched relative to WT in both the transcriptome and the proteome (NES = −1.97 and −1.99).

The oligodendrocyte marker set was positively enriched in the transcriptome at both ages (NES = +2.38, FDR = 2.5e-06 at 4 years and NES = +3.88, FDR = 8.6e-10 at 8 years), but not in the proteome (NES = +0.89, FDR = 0.74 at 7 to 8 years, Fig. 3A). At 8 years 51 of the 60 marker genes were significant DEGs, while none was at 4 years (Fig. 3C). All 51 had a positive log2 fold change, that is, higher gene expression in TgHD than in WT, and they were more tightly grouped than the markers of other cell types (Supplementary Fig. S7).

The astrocyte and microglial marker sets are reported with the immune findings in Section 3.4.

#### 3.2.2. Single candidate genes at 4 and 8 years

We then examined single genes of the cell-type marker sets, comparing their TgHD versus WT fold changes at 4 and at 8 years. For each cell type we selected the eight genes with the largest concordant change in the 8-year transcriptome and the 7- to 8-year proteome, and plotted their values at both ages (Fig. 3D and E for D2 MSN, Fig. 3F and G for oligodendrocytes). All eight D2 MSN genes had positive transcript log2 fold changes at 4 years, none of them significant, and were significantly downregulated at 8 years (Fig. 3D). *SCN4B* had the largest transcript change. In the proteome GPR88 and RGS9 had the largest change between the two ages (Fig. 3E). The widely used marker *PPP1R1B* was significantly downregulated in the transcriptome at 8 years, with a log2 fold change (log2FC) of −1.67 (FDR = 0.003), while its protein abundance changed little (log2FC = −0.17, FDR = 0.997).

Among the D1 markers, the pattern was less uniform. The clearest case was *NXPH4*, a ligand of alpha-neurexins. It was significantly downregulated in the transcriptome at 8 years (log2FC = −2.08, FDR = 0.03, Supplementary Fig. S8). It was also lower in TgHD than in WT in the 7- to 8-year proteome (log2FC = −0.59, FDR = 0.997), where its nominal P value was the second lowest of any protein (P = 7.8e-04).

Among the oligodendrocyte markers, all eight selected genes were significantly upregulated in TgHD compared with WT in the transcriptome at 8 years, with the largest change in *KLKc* (log2FC = +2.08), followed by *MOBP*, *ANLN*, *MBP*, *GJB1*, *APOD*, *MOG* and *PLP1* (Fig. 3F and G).

#### 3.2.3. Negative enrichment of synaptic pathways at 8 years

The pathways of the synaptic-transmission and synaptic-structure themes (Methods 2.9, Supplementary Table S7) were compared in TgHD relative to WT animals at 4 and 8 years (Fig. 4A and B).

**Figure 4.**
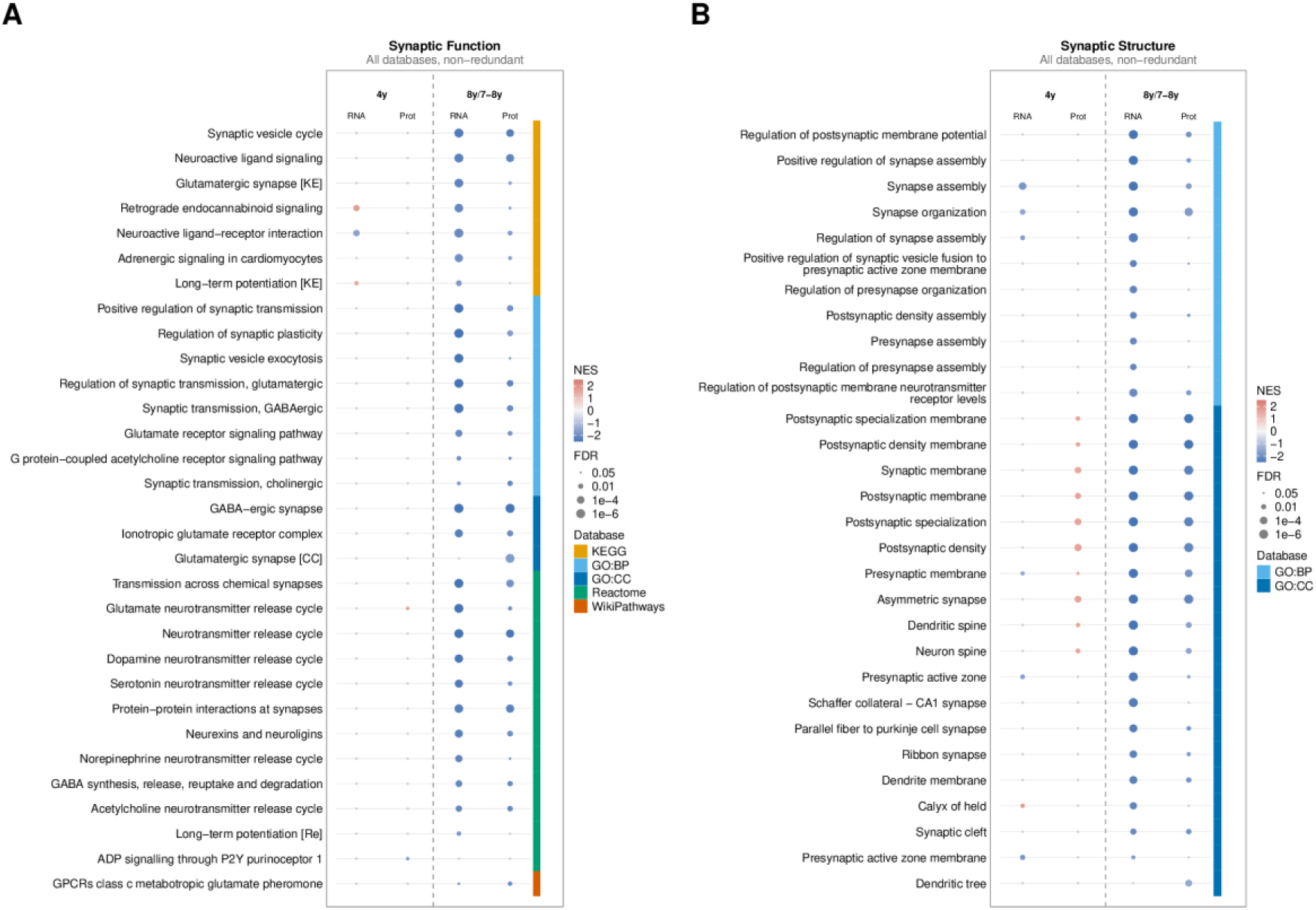
Synaptic pathway enrichment across the transcriptome and proteome. **(A, B)** Dot heatmaps of GSEA NES of pathways assigned by keyword matching (Methods 2.9, Supplementary Table S7) to the synaptic-transmission theme (A) and to the synaptic-structure theme (B), from GO:BP, GO:CC, KEGG, Reactome and WikiPathways. The colour strip marks the source database. Pathways significant in at least one dataset were ranked by the number of datasets in which they reached significance, with pathways significant in the proteome ranked first (Methods 2.10), and the top 30 are shown. Redundant GO:BP and GO:CC parent pathways were removed and pathways with identical gene membership are shown once (Methods 2.10). In both panels, columns are RNA-seq (RNA) and age-normalised proteomics (Prot), the later age being 8 years for RNA-seq and 7 to 8 years for proteomics (8y/7-8y), and all comparisons are TgHD relative to WT. Grey dots mark comparisons not reaching an FDR below 0.05. The full pathway list is in Supplementary Table S5. Abbreviations. FDR, false discovery rate. GO:BP, Gene Ontology biological process. GO:CC, Gene Ontology cellular component. GSEA, gene set enrichment analysis. NES, normalised enrichment score. TgHD, transgenic Huntington’s disease minipig. WT, wild type.

At 4 years synaptic pathways were barely affected in TgHD compared with WT. Of the 98 transmission and 80 structure pathways tested, 4 and 9 were significant, with no consistent direction of enrichment. By 8 years nearly every significant synaptic pathway was negatively enriched, 79 of 98 transmission and 70 of 80 structure pathways in the transcriptome and 39 of 93 and 29 of 76 in the proteome (Fig. 4A and B, Supplementary Table S5). The strongest negative enrichment was in the postsynaptic compartment, where the postsynaptic specialisation membrane and the postsynaptic density membrane had the two lowest NES in the transcriptome (−3.22 and −3.15) and again in the proteome (−2.75 and −2.65). Among functional pathways the GABAergic synapse, glutamatergic synaptic transmission, neurotransmitter secretion and the synaptic vesicle cycle were the most affected, in line with the negative enrichment of the D2 MSN marker set described above.

A few pathways, all related to excitatory synaptic signalling, had a significantly positive NES at 4 years and a significantly negative one at 8 years, in each case TgHD relative to WT animals of that age. Long-term potentiation and retrograde endocannabinoid signalling did so in the transcriptome, and the glutamate neurotransmitter release cycle in the proteome (Fig. 4A). In the proteome a group of nested synaptic compartment gene sets also changed from positive to negative, including the postsynaptic density, the postsynaptic and presynaptic membranes and the dendritic spine (Fig. 4B).

#### 3.2.4. Transcription factors associated with the downregulated genes

To identify potential upstream regulators of the transcriptional changes, we performed transcription factor enrichment analysis with ChEA3 (Keenan et al., 2019) on the genes downregulated in TgHD at 8 years (Supplementary Table S13). The top-ranked factor was MYT1L (rank 1, score 3.33), followed by SCRT1 (rank 2), DPF1 (rank 8), NEUROD6 (rank 10), RORB (rank 12) and FOXG1 (rank 14), all of which have established roles in neuronal identity.

### 3.3. Opposite enrichment of mitochondrial and oxidative phosphorylation pathways at 4 and 8 years

#### 3.3.1. Mitochondrial and OXPHOS pathways are positively enriched at 4 years and negatively at 8 years

Mitochondrial and OXPHOS pathways were positively enriched in both the transcriptome and the proteome in the putamen of 4-year-old TgHD minipigs compared with age-matched WT animals (Fig. 5A). At 8 years the same pathways were negatively enriched relative to age-matched WT, often with a more negative NES in the proteome. Of the 82 pathways of the mitochondrial theme tested in the transcriptome, 57 were significant at 4 years, all positively enriched, and 33 at 8 years, all negatively enriched. In the proteome 60 of the 77 tested pathways were significant at 7 to 8 years, all negatively enriched (Supplementary Fig. S9, Supplementary Table S5). The most negative NES values in the proteome were those of the mitochondrial protein-containing complex (−3.04) and the mitochondrial ribosome (−2.92). These counts do not include the MitoCarta gene sets, which are reported below.

**Figure 5.**
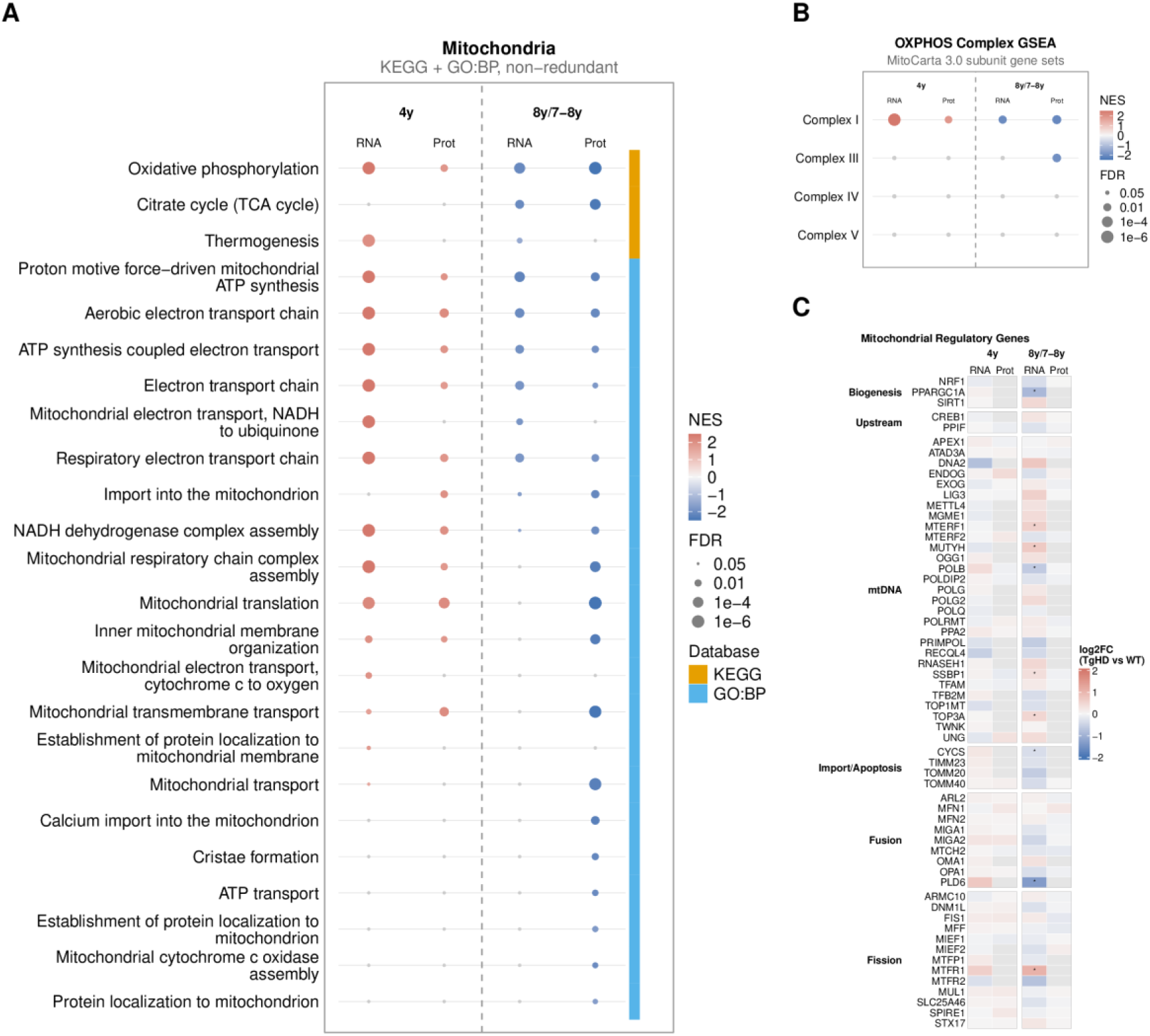
Mitochondrial and oxidative phosphorylation pathway enrichment and mitochondrial regulatory genes. **(A)** Dot heatmap of GSEA NES of pathways assigned to the mitochondrial and OXPHOS theme by keyword matching (Methods 2.9, Supplementary Table S7), from KEGG and GO:BP. The colour strip marks the source database. Pathways significant in at least one dataset were ranked by the number of datasets in which they reached significance, with pathways significant in the proteome ranked first (Methods 2.10), and the top 25 are shown. Redundant GO:BP parent pathways were removed and pathways with identical gene membership are shown once (Methods 2.10). **(B)** Dot heatmap of GSEA NES of the individual OXPHOS complex subunit gene sets of MitoCarta 3.0 (Rath et al., 2020). Complex II is absent because its gene set of four subunits is below the minimum of five genes required for GSEA. **(C)** Heatmap of log2 fold change of selected regulators of mitochondrial biogenesis, mitochondrial DNA maintenance, protein import, fusion and fission, with rows grouped by function. Grey cells in (C) mark genes not detected in that dataset. Asterisks mark RNA-seq FDR significance, * below 0.05, ** below 0.01, *** below 0.001. Throughout, columns are RNA-seq (RNA) and age-normalised proteomics (Prot), the later age being 8 years for RNA-seq and 7 to 8 years for proteomics (8y/7-8y), and all comparisons are TgHD relative to WT. Grey dots in (A) and (B) mark comparisons not reaching an FDR below 0.05. All significant mitochondrial pathways are in Supplementary Fig. S9 and the full pathway list in Supplementary Table S5. Abbreviations. FDR, false discovery rate. GO:BP, Gene Ontology biological process. GSEA, gene set enrichment analysis. NES, normalised enrichment score. OXPHOS, oxidative phosphorylation. TgHD, transgenic Huntington’s disease minipig. WT, wild type.

Oxidative phosphorylation (KEGG) was significantly enriched in TgHD against WT in all four datasets, positively at 4 years and negatively at the later age (NES in the transcriptome +2.95 at 4 years and −2.08 at 8 years, in the proteome +1.90 at 4 years and −2.53 at 7 to 8 years, Fig. 5A). The aerobic electron transport chain (GO:BP) reversed between the two ages as well (transcriptome +2.88 and −2.04, proteome +2.11 and −2.09).

We also applied GSEA to the curated mitochondrial gene sets of MitoCarta 3.0 (Rath et al., 2020). We found the same reversal in the TgHD versus WT comparison, positive at 4 years and negative at the later age (Supplementary Fig. S10). Three gene sets were significant in all four datasets, the OXPHOS subunits (NES in the transcriptome +2.84 at 4 years and −2.18 at 8 years, in the proteome +1.99 at 4 years and −2.35 at 7 to 8 years), the complex I subunits (transcriptome +2.80 and −2.04, proteome +1.89 and −2.09) and the inner mitochondrial membrane set (transcriptome +2.08 and −1.56, proteome +2.10 and −2.57). The tricarboxylic acid (TCA) cycle set was positive but not significant in the 4-year transcriptome (NES = +1.79, FDR = 0.051) and was significantly negatively enriched in the 7- to 8-year proteome (NES = −2.45).

The mitochondrial ribosome set differed between the transcriptome and the proteome at the later age. It was positively enriched in TgHD relative to WT at 4 years in both (transcriptome +2.29, proteome +2.07). At 8 years it was not significant in the transcriptome (NES = −0.87), whereas in the 7- to 8-year proteome it was the most negative of all MitoCarta sets (NES = −2.88).

The subunit sets of whole complexes were significantly enriched in opposite directions at the two ages (Fig. 5B), while most individual subunit genes were not significant at either age. In the transcriptome most OXPHOS subunit genes had a positive log2 fold change (TgHD versus WT) at 4 years and a negative one at 8 years, neither of them significant (Supplementary Fig. S11, Supplementary Table S2). Only two of the subunit transcripts had an FDR below 0.05, both downregulated in the transcriptome at 8 years, *NDUFA5* (complex I, log2FC = −0.82, FDR = 0.0016) and *ATP5MG* (complex V, log2FC = −0.54, FDR = 0.047). For whole complexes, the complex I subunit set changed from positive to negative enrichment in both the transcriptome and the proteome, as described above, and complex III was negatively enriched in the 7- to 8-year proteome (NES = −1.96, Fig. 5B).

#### 3.3.2. Mitochondrial regulators, mtDNA integrity and calcium pathways

Next we examined regulators of mitochondrial biogenesis, dynamics and mtDNA maintenance (Fig. 5C). *PPARGC1A* encodes PGC-1α, a master regulator of mitochondrial biogenesis (Fernandez-Marcos and Auwerx, 2011), and its reduced striatal expression is one of the established mitochondrial findings in HD (Cui et al., 2006; Weydt et al., 2006). In our data *PPARGC1A* was unchanged at 4 years and significantly downregulated in TgHD at 8 years in the transcriptome (log2FC = −0.91, FDR = 0.029). PGC-1α was not detected in the proteome. Further biogenesis regulators had negative but not significant log2 fold changes in the transcriptome at 8 years, *PPARGC1B* (log2FC = −0.72, FDR = 0.091), *ESRRA* (−0.65, FDR = 0.092) and *NRF1* (−0.36, FDR = 0.088), while *TFAM* and *CREB1* did not change.

The parallel changes in biogenesis regulators and OXPHOS pathway enrichment raise the question of whether altered mitochondrial homeostasis affects the mitochondrial genome, either through oxidative damage or through mitochondrial biogenesis. We therefore measured mtDNA damage by RADF (Methods 2.15) and mtDNA copy number by qPCR at 7 years. Neither nuclear nor mitochondrial DNA damage differed between genotypes, and mtDNA copy number was slightly but not significantly higher in TgHD cortex, with no change in putamen (Supplementary Fig. S12B and C).

Because oxidative phosphorylation pathways were negatively enriched at 8 years, we examined whether glycolysis pathways were enriched in the opposite direction. No glycolysis pathway was positively enriched at 8 years. Canonical glycolysis (NES = −1.75, FDR = 0.046) and the response to glucose (NES = −1.76, FDR = 0.0022) were significantly negatively enriched in TgHD relative to WT in the 8-year transcriptome, and no glycolysis pathway was significant in the proteome. The only positively enriched carbohydrate pathways were the pentose phosphate pathway (NES = +1.79, FDR = 0.034) and carbohydrate phosphorylation (NES = +1.89, FDR = 0.044), both in the 7- to 8-year proteome (Supplementary Fig. S13A). In the 8-year transcriptome, both the oxidative and the glycolytic pathways were therefore negatively enriched.

Mitochondrial calcium handling is reported to be impaired in HD models (Milakovic et al., 2006; Panov et al., 2002), so we also examined the calcium-handling pathways. Like the mitochondrial pathways, the calcium-related pathways were positively enriched in TgHD relative to WT at 4 years and negatively enriched at 8 years (Supplementary Fig. S13B, Supplementary Table S5). At 4 years every significant calcium pathway was positively enriched, 5 of 89 tested in the transcriptome and 14 of 85 in the proteome. At the later age every significant pathway was negatively enriched, 64 of 89 in the transcriptome and 25 of 87 in the proteome.

The KEGG calcium signalling pathway was positively enriched in the 4-year proteome (NES = +1.61, FDR = 0.047) and negatively enriched in the 8-year transcriptome (NES = −1.85, FDR = 1.9e-05) and in the 7- to 8-year proteome (NES = −1.77, FDR = 0.0022). Calcium ion import into the cytosol was negatively enriched in both the 8-year transcriptome (NES = −2.04, FDR = 0.0023) and the 7- to 8-year proteome (NES = −2.02, FDR = 0.021). Mitochondrial calcium ion transmembrane transport was negatively enriched in the 7- to 8-year proteome only (NES = −2.29, FDR = 0.0023, 20 genes). The most negative calcium pathway overall was the calcium channel complex in the 8-year transcriptome (GO:CC, NES = −2.71, FDR = 2.4e-09), a set of neuronal channel genes rather than mitochondrial ones. This agrees with the negative enrichment of the cardiac pathways in Fig. 2C, whose gene sets are built from the same calcium-handling genes (Section 3.1.5).

### 3.4. Immune pathways, microglial markers and complement

#### 3.4.1. Immune and inflammation pathways are positively enriched at 8 years

Toll-like receptor signalling was among the 20 KEGG pathways with the highest absolute NES (Fig. 2C), with positive enrichment in both the transcriptome and the proteome of the older cohort, so we examined the immune and inflammation pathways in more detail. Of the 508 immune pathways tested in the transcriptome for TgHD against WT animals, 317 were significant at 8 years, and all but four were positively enriched (Fig. 6A). At 4 years only 39 were significant, 16 positively and 23 negatively enriched. In the 7- to 8-year proteome, 24 of 424 tested pathways were significant, 23 of them positively enriched, and 14 of the 24 were also significant in the 8-year transcriptome (Supplementary Table S5).

**Figure 6.**
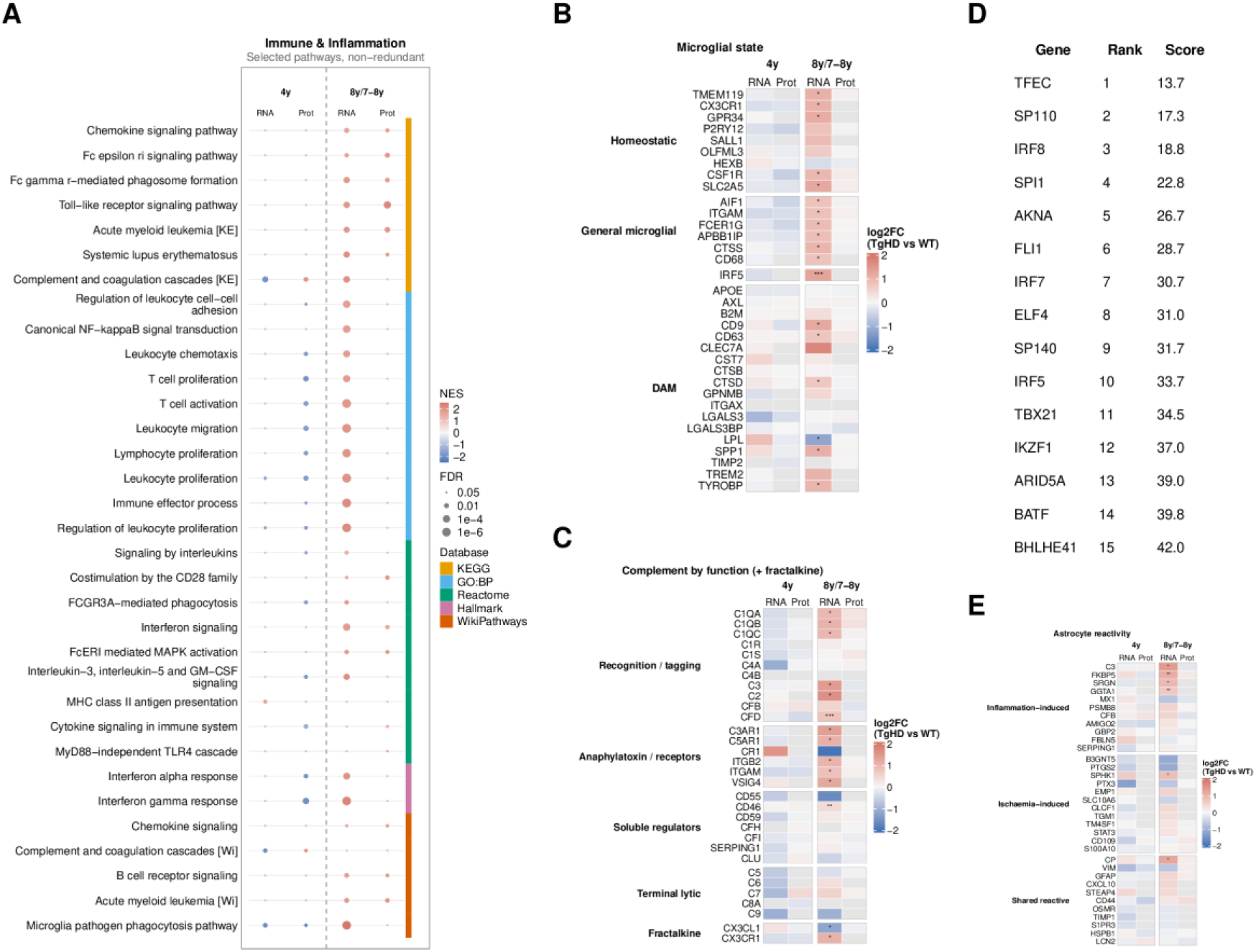
Immune pathway enrichment, microglial markers, complement, transcription factors and astrocyte markers. **(A)** Dot heatmap of GSEA NES of pathways assigned to the immune and inflammation theme by keyword matching (Methods 2.9, Supplementary Table S7), from GO:BP, KEGG, Reactome, Hallmark and WikiPathways. The colour strip marks the source database. Pathways named after a specific infectious disease were excluded by a keyword match on the pathway name. Of the remainder, a pathway was kept if it reached significance in a proteomics dataset, if it reached significance in both RNA-seq and proteomics at the same age with concordant NES, positive in both or negative in both, or if its mean NES reversed between 4 and 8 years. Pathways significant in at least one dataset were ranked by the number of datasets in which they reached significance, with pathways significant in the proteome ranked first (Methods 2.10), and the top 30 are shown. Canonical NF-κB signal transduction, MHC class II antigen presentation and the MyD88-independent TLR4 cascade were manually retained and are shown in addition to the top 30. Redundant GO:BP parent pathways were removed and pathways with identical gene membership are shown once (Methods 2.10). **(B)** Heatmap of log2 fold change of microglial genes, with rows grouped into homeostatic genes, general microglial genes expressed in both homeostatic and disease-associated states (Paolicelli et al., 2022), the interferon regulatory factor gene *IRF5*, and the genes of the DAM signature (Keren-Shaul et al., 2017; Krasemann et al., 2017). The general microglial group is our grouping and not a published classification. **(C)** Heatmap of log2 fold change of the genes of the complement cascade and of the fractalkine pair, *CX3CL1* and *CX3CR1*, with rows grouped by function into recognition and tagging components, anaphylatoxin and phagocytic receptors, soluble regulators and the terminal lytic components. **(D)** Table of the 15 transcription factors ranked highest by ChEA3 (Keenan et al., 2019) for the genes upregulated at 8 years in TgHD, with the factor, its rank and its integrated mean-rank score. ChEA3 calculates a rank without a significance test, and a lower rank indicates stronger enrichment. **(E)** Heatmap of log2 fold change of the inflammation-induced, ischaemia-induced and shared reactive astrocyte marker sets (Supplementary Table S8). Grey cells in (B), (C) and (E) mark genes not detected in that dataset. Asterisks mark RNA-seq FDR significance, * below 0.05, ** below 0.01, *** below 0.001. Throughout, columns are RNA-seq (RNA) and age-normalised proteomics (Prot), the later age being 8 years for RNA-seq and 7 to 8 years for proteomics (8y/7-8y), and all comparisons are TgHD relative to WT. Grey dots in (A) mark comparisons not reaching an FDR below 0.05. The all-database immune panel is Supplementary Fig. S14 and the full ChEA3 output is in Supplementary Table S13. Abbreviations. DAM, disease-associated microglia. FDR, false discovery rate. GO:BP, Gene Ontology biological process. GSEA, gene set enrichment analysis. NES, normalised enrichment score. [KE], KEGG. [Wi], WikiPathways. TgHD, transgenic Huntington’s disease minipig. WT, wild type.

The pathways positively enriched in TgHD compared with WT in the 8-year transcriptome included interferon-alpha and interferon-gamma signalling, Toll-like receptor signalling, Fc-gamma-receptor-mediated phagocytosis and canonical NF-κB signalling, the last with an NES of +1.75 (Fig. 6A, Supplementary Fig. S14). In the proteome Toll-like receptor signalling (NES = +2.18) and the MyD88-independent TLR4 cascade (NES = +1.64) were significant, while NF-κB signalling was positive but not significant (NES = +1.53, single genes in Supplementary Fig. S15). *TLR4* itself was higher in TgHD at 8 years but not significantly (log2FC = +0.75, FDR = 0.095, Supplementary Fig. S15), and the protein was not detected. At 4 years the complement and coagulation cascades were positively enriched in the proteome (NES = +1.97) and negatively enriched in the transcriptome (NES = −1.90).

#### 3.4.2. Homeostatic and general microglial markers are increased at 8 years

The microglial marker set was positively enriched in the 8-year transcriptome (NES = +2.54, Fig. 3A), so we examined the individual microglial genes (Fig. 6B). The homeostatic markers *TMEM11S*, *CX3CR1*, *GPR34*, *CSF1R* and *SLC2A5* were significantly upregulated in the TgHD putamen at 8 years, while expression of *P2RY12* was increased but not significantly (log2FC = +0.82, FDR = 0.119). The general microglial markers *CDc8*, *ITGAM*, *AIF1*, *FCER1G*, *APBB1IP* and *CTSS* were also significantly upregulated.

In mouse models, the disease-associated microglia (DAM) signature consists of lower expression of homeostatic genes, such as *CX3CR1* and *P2RY12*, with higher expression of *APOE*, *AXL*, *SPP1*, *TREM2*, *ITGAX*, *CLEC7A*, *LGALS3* and *LPL* (Keren-Shaul et al., 2017; Krasemann et al., 2017; Paolicelli et al., 2022). In the TgHD putamen at 8 years, the homeostatic genes were increased, not decreased. Of these eight DAM genes, only *SPP1* was significantly higher in TgHD (log2FC = +1.11, FDR = 0.013). *TREM2*, which is required for the transition to the DAM state (Paolicelli et al., 2022), was higher but not significantly (log2FC = +1.05, FDR = 0.056). *APOE*, *AXL* and *LGALS3* were unchanged; *LPL* was decreased (log2FC = −1.20, FDR = 0.018), and *ITGAX* was not detected. Other genes of the DAM set, *CDS*, *TYROBP*, *CDc3* and *CTSD*, were more highly expressed in TgHD. Of all the microglial genes examined, the interferon regulatory factor gene *IRF5* had the lowest FDR in the 8-year transcriptome (log2FC = +1.30, FDR = 8.4e-06).

In the 7- to 8-year proteome, 11 of the 15 homeostatic and general microglial proteins were quantified; none differed between genotypes (log2 fold changes within ±0.16, Supplementary Table S4), and the microglial marker set was not enriched (Fig. 3A). Bulk RNA sequencing cannot distinguish a larger number of microglia from higher expression per cell. At 4 years the microglial set was not enriched in the transcriptome and was negatively enriched in the proteome (NES = −2.11, Fig. 3A).

#### 3.4.3. Complement genes are upregulated and the fractalkine transcript is reduced

We examined the complement genes and fractalkine signalling (CX3CL1 and its receptor CX3CR1), because both regulate microglial engulfment of synapses (Saba et al., 2022) and because factors of the complement cascade are among the transcriptional targets of NF-κB (Oeckinghaus and Ghosh, 2009). The complement gene set was positively enriched in the 8-year transcriptome (NES = +2.37, FDR = 1.5e-05, Supplementary Table S6). The genes of the three C1q chains (*C1ǪA*, *C1ǪB*, *C1ǪC*), of the early components C2 and C3 (*C2*, *C3*), of the alternative-pathway factor D (*CFD*) and of the anaphylatoxin receptors C3aR and C5aR (*C3AR1*, *C5AR1*) were upregulated in TgHD relative to WT animals, while the terminal component genes *C5* to *CS* were unchanged (Fig. 6C). Of the complement inhibitors in Fig. 6C, only *CD4c* changed significantly, upregulated in TgHD at 8 years (log2FC = +0.41, FDR = 0.0077). None of the complement proteins detected in the proteome changed.

*ITGAM*, which encodes the CD11b subunit of the complement receptor CR3, and the microglial fractalkine receptor gene *CX3CR1* were upregulated in TgHD against WT at 8 years, while the neuronal ligand gene *CX3CL1* was downregulated. CX3CL1 protein was unchanged and CX3CR1 protein was not detected (Fig. 6C).

#### 3.4.4. Microglial and immune transcription factors are associated with the upregulated genes

We applied transcription factor enrichment analysis with ChEA3 to the 1,488 genes upregulated at 8 years in TgHD. All ten top-ranked factors are transcription factors of microglia, macrophages or other immune cells (Fig. 6D, Supplementary Table S13), in contrast to the neuronal-identity factors found for the downregulated genes (Section 3.2.4). The highest ranked was TFEC (rank 1, score 13.67), a MiT family factor expressed in macrophages and microglia (Rehli et al., 1999), followed by IRF8 (rank 3), which is required for microglial identity (Saeki et al., 2024), and SPI1 (rank 4). *SPI1* encodes PU.1, a transcription factor that determines the identity of microglia and macrophages. Mutant HTT was reported to induce an inflammatory transcriptional response in microglia through PU.1 (Crotti et al., 2014). Three interferon regulatory factors were among the top ten, IRF8, IRF7 and IRF5. The transcripts of seven of the 12 highest-ranked factors were themselves significantly higher in TgHD at 8 years, *IRF5*, *FLI1*, *IKZF1*, *ELF4*, *TFEC*, *SPI1* and *IRF8* (Supplementary Table S2).

#### 3.4.5. Classical astrocyte markers are unchanged, while the inflammation-induced marker set is positively enriched

Escartin et al. (2021) defined three reactive astrocyte marker sets (Methods 2.8). Only the inflammation-induced set was positively enriched in TgHD relative to WT in the 8-year transcriptome (NES = +1.73, FDR = 0.038). The ischaemia-induced set (NES = +0.81, FDR = 0.71) and the shared reactive set (NES = +1.11, FDR = 0.36) were not enriched, and none of the three was enriched in the proteome (Fig. 6E, Supplementary Table S6). The general astrocyte marker set was not enriched at 8 years either (NES = −1.17, FDR = 0.28), and the classical markers *GFAP* and *VIM* were unchanged. Four inflammation-induced genes, *C3*, *FKBP5*, *SRGN* and *GGTA1*, and the shared reactive marker *CP* were significantly upregulated in the 8-year transcriptome. However, *C3* is also expressed by microglia, so its increase cannot be specifically assigned to astrocytes.

### 3.5. Alternative splicing, RNA processing and DNA repair

#### 3.5.1. Alternative splicing changes are more frequent at 8 years in TgHD

We quantified five event types with rMATS (Shen et al., 2014), skipped exon, mutually exclusive exon, alternative 3’ and 5’ splice site and retained intron. Skipped exon was the most frequently changed event type in every comparison (Fig. 7A). Retained introns were the rarest type, but a larger share of them changed than of the other types (38 of 515 at 8 years, 7.4% against 3.8%), 25 of the 38 with more retention in TgHD (Supplementary Table S9). In the TgHD versus WT comparison, the number of significantly changed events increased almost fivefold between the two ages, from 509 at 4 years to 2,490 at 8 years (Fig. 7A). At 8 years 1,217 events gained and 1,273 lost inclusion in TgHD (Supplementary Fig. S16). Splicing also changed with age, more in TgHD than in WT animals, with 1,728 changed events between 4 and 8 years in TgHD and 916 in WT (Fig. 7A).

**Figure 7.**
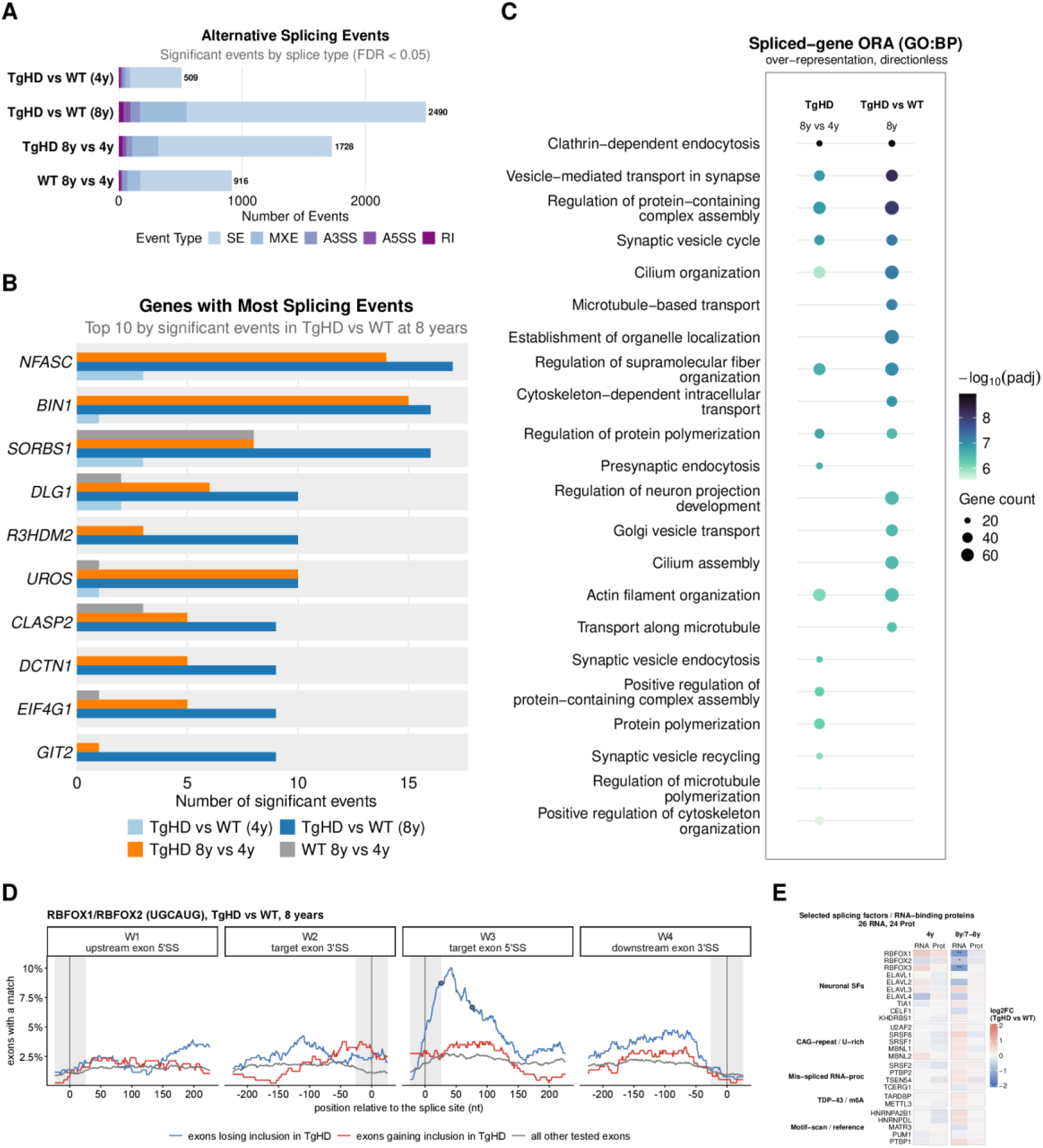
Alternative splicing events, differentially spliced genes, the RBFOX motif and splicing regulators. **(A)** Stacked bar chart of the significant alternative splicing events per comparison, coloured by event type. The comparisons are TgHD versus WT at 4 years and at 8 years, TgHD 8 years versus 4 years, and WT 8 years versus 4 years. **(B)** Grouped horizontal bar chart of the number of significant splicing events per gene. The ten genes with the most events in TgHD versus WT at 8 years are shown, ordered by that count, and the four bars of each gene are the four comparisons of (A). A pale grey slot marks a comparison in which the gene has no significant events. **(C)** Dot heatmap of the ORA of the differentially spliced genes (Methods 2.13), for TgHD 8 years versus 4 years and for TgHD versus WT at 8 years. **(D)** Motif map of the RBFOX motif UGCAUG around skipped exons, TgHD relative to WT at 8 years. The four sub-panels are the four splice sites of a skipped-exon event, with position 0 at the splice site. The y-axis shows the percentage of exons with a motif match in the 50-nucleotide window centred at that position, for the three exon groups in the legend. Grey bands cover windows that still contain exonic sequence, which were not tested. The test compares each group with the exons whose inclusion did not change (Methods 2.12), and a filled circle marks a position with an FDR below 0.05. RBFOX1 and RBFOX2 share the panel because their ATtRACT motifs have the same six-nucleotide consensus and identical values at every position. Supplementary Fig. S17 shows every protein tested. **(E)** Heatmap of log2 fold change of curated splicing regulators and RNA-binding proteins. Grey cells in (E) mark genes not detected in that dataset. Asterisks mark RNA-seq FDR significance, * below 0.05, ** below 0.01, *** below 0.001. The full event list is in Supplementary Table S9 and the full motif test results in Supplementary Table S10. Abbreviations. A3SS, alternative 3’ splice site. A5SS, alternative 5’ splice site. FDR, false discovery rate. MXE, mutually exclusive exon. ORA, over-representation analysis. RI, retained intron. SE, skipped exon. TgHD, transgenic Huntington’s disease minipig. WT, wild type.

#### 3.5.2. Differentially spliced genes are mainly synaptic and cytoskeletal

Altered splicing at 8 years affected mainly neuronal, synaptic and cytoskeletal genes, and for some of them it appeared only in the older TgHD animals. The ten genes with the most significant alternative splicing (AS) events in the 8-year TgHD versus WT comparison had 9 to 17 events each, headed by *NFASC* (17), *BIN1* (16) and *SORBS1* (16), whereas none of them had more than three events in the same comparison at 4 years (Fig. 7B). Splicing of *NFASC* and *BIN1* did not change during WT ageing, while both changed extensively during TgHD ageing (14 and 15 events). *SORBS1* changed in both genotypes, but largely at different events. Only three of its eight events found during WT ageing were also found during TgHD ageing, and *UROS*, another of the ten, shared none of its events between the two (one event in WT ageing, ten in TgHD ageing).

Over-representation analysis of the 1,437 genes differentially spliced between TgHD and WT animals at 8 years showed enrichment of vesicle-mediated transport in synapse, the synaptic vesicle cycle, clathrin-dependent endocytosis, cilium organisation and microtubule-based transport (Fig. 7C). Among KEGG pathways, lysosome biogenesis, endocytosis and spinocerebellar ataxia were enriched (Supplementary Table S16). At 4 years far fewer genes were differentially spliced (375), and only six GO terms were enriched, four of them cilium or spermatid terms that share their genes (Supplementary Table S16). The proteins of the ten most differentially spliced genes (Fig. 7B) did not differ in abundance between genotypes at 7 to 8 years (Supplementary Table S4).

#### 3.5.3. The RBFOX motif is enriched downstream of exons that lose inclusion

Altered splicing can result from changes in RBP abundance or availability, and expanded CAG repeat RNA has been reported to bind splicing factors (Sathasivam et al., 2013; Schilling et al., 2019). We therefore asked whether the exons whose inclusion changed in TgHD minipigs contain specific RBP motifs at particular distances from their splice sites. Each skipped-exon event was scanned at its four splice sites for motifs from the human ATtRACT database and the *Sus scrofa* set of CisBP-RNA (Methods 2.12).

At 8 years the RBFOX motif UGCAUG was enriched in the intron immediately downstream of the exons that lost inclusion in TgHD (Fig. 7D). The motif was present downstream of 8.7% of these exons against 2.6% of the unchanged exons (47 of 539 against 91 of 3,558), a 3.41-fold enrichment at 26 nucleotides into the intron (FDR = 2.9e-07). Enrichment decreased with distance from the splice site, to 2.73-fold at 76 nucleotides (FDR = 0.0015), and was not significant at 126 nucleotides. The motif was not enriched at the other three splice sites or among the exons that gained inclusion. It was also enriched at the same position among the exons that lost inclusion between 4 and 8 years in TgHD animals (2.80-fold, FDR = 0.040, Supplementary Table S10).

In the transcriptome all three RBFOX genes were downregulated in TgHD at 8 years (*RBFOX1* log2FC = −1.39, FDR = 0.0054, *RBFOX2* −0.68, FDR = 0.048, *RBFOX3* −1.41, FDR = 0.0016), and none was changed at 4 years (Fig. 7E, Supplementary Table S2). Fourteen further RBP tests were significant at 8 years (Supplementary Fig. S17, Supplementary Table S10). Two came from the RBFOX1 entry, whose motif is identical to that of RBFOX2. Ten were tests of short or low-complexity motifs, such as poly-U. Many proteins bind such motifs, so these tests cannot be attributed to one protein. The remaining two belonged to one motif family, ACUAAC, assigned to SF1 and QKI. The SF1 motif UACUAAC was enriched in the intron upstream of the exons that lost inclusion (12.0-fold, 13 of 542 exons against 7 of 3,580, FDR = 1.6e-04), at the branch point position. The QKI motif ACUAACA was enriched downstream of the exons that gained inclusion (8.04-fold, 9 of 400 exons against 10 of 3,558, FDR = 0.018, Supplementary Fig. S18). No test was significant at 4 years or in WT ageing.

#### 3.5.4. Enrichment of splicing and RNA-processing pathways

The pathways of the splicing and RNA-processing machinery were positively enriched in the TgHD transcriptome relative to WT at both ages. The number of significant pathways increased from 13 of 68 tested at 4 years to 58 at 8 years, with none negatively enriched at either age. At 8 years they included the spliceosome, mRNA splicing and nonsense-mediated decay. In the proteome 14 of the 15 significant pathways were negatively enriched at 4 years, whereas all 24 significant pathways were positively enriched at 7 to 8 years (Supplementary Fig. S19A and B, Supplementary Table S5).

#### 3.5.5. DNA-repair genes are upregulated at 8 years, while MSH3 and FAN1 are unchanged

Base excision repair was positively enriched in TgHD relative to WT in both the transcriptome and the proteome of the older cohort (Fig. 2C), so we examined the DNA-repair genes in more detail. In human HD, variants in DNA-repair genes, mainly those of mismatch repair, modify the age at onset and promote somatic expansion of the CAG repeat (Lee et al., 2019, 2015), whereas the mixed CAG/CAA repeat of this model is not expected to expand.

Of the 27 DNA-repair genes examined, seven were differentially expressed between TgHD and WT animals at 8 years and none at 4 years (Fig. 8A, Supplementary Table S2). *ATM* (log2FC = +0.83), *FEN1* (+0.78), *MUTYH* (+0.68), *PNKP* (+0.55) and *MLH1* (+0.46) were upregulated, and *POLB* (−0.63) and *NEIL2* (−1.22) were downregulated. *LIG3* (+0.58) and *XRCC1* (+0.37) had positive log2 fold changes and *PMS2* (−0.35) a negative one; none was significant. The somatic-instability modifiers *MSH3* and *FAN1* were unchanged (log2FC = +0.06 and −0.10). Fifteen of the 27 proteins were quantified and none differed significantly between genotypes, with log2 fold changes between −0.16 and +0.22 (Fig. 8A, Supplementary Table S4).

**Figure 8.**
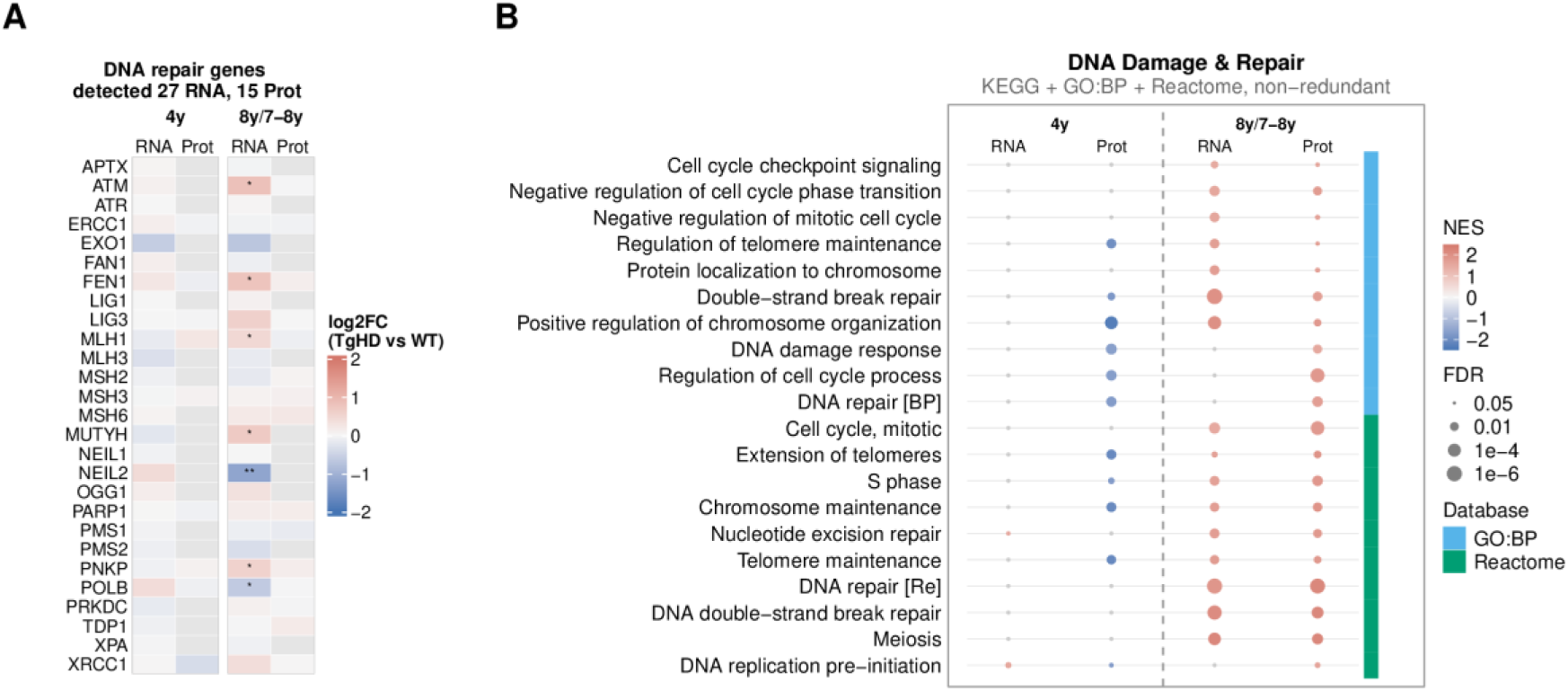
DNA-repair genes and cell-cycle and DNA-repair pathway enrichment. **(A)** Heatmap of log2 fold change of the 27 curated DNA-repair genes. Grey cells in (A) mark genes not detected in that dataset. Asterisks mark RNA-seq FDR significance, * below 0.05, ** below 0.01, *** below 0.001. **(B)** Dot heatmap of GSEA NES of pathways assigned to the cell cycle and DNA repair theme by keyword matching (Methods 2.9, Supplementary Table S7), from KEGG, GO:BP and Reactome. The colour strip marks the source database. Pathways significant in at least one dataset were ranked by the number of datasets in which they reached significance, with pathways significant in the proteome ranked first (Methods 2.10), and the top 20 are shown. Redundant GO:BP parent pathways were removed and pathways with identical gene membership are shown once (Methods 2.10). In both panels, columns are RNA-seq (RNA) and age-normalised proteomics (Prot), the later age being 8 years for RNA-seq and 7 to 8 years for proteomics (8y/7-8y), and all comparisons are TgHD relative to WT. Grey dots in (B) mark comparisons not reaching an FDR below 0.05. The full pathway list is in Supplementary Table S5. Abbreviations. FDR, false discovery rate. GO:BP, Gene Ontology biological process. GSEA, gene set enrichment analysis. NES, normalised enrichment score. TgHD, transgenic Huntington’s disease minipig. WT, wild type.

Pathways of the cell cycle and DNA repair theme were predominantly positively enriched in TgHD compared with WT in the 8-year transcriptome. Of the 213 pathways tested, 132 were significant and 131 of them positive (Fig. 8B, Supplementary Table S5). They included base excision repair, nucleotide excision repair and double-strand break repair. The KEGG and Reactome mismatch repair pathways were not significant.

DNA replication was among the most strongly enriched pathways of this theme (KEGG, NES = +2.39, FDR = 7.3e-05). Pathways of programmed cell death were also positively enriched in TgHD against WT in the 8-year transcriptome, among them apoptosis (KEGG, NES = +1.76, FDR = 0.0018) and the apoptotic execution phase (Reactome, NES = +2.12, FDR = 7.2e-04). Among the Gene Ontology terms, the leukocyte and the lymphocyte apoptotic process terms were significant, whereas the neuron apoptotic process term was not (NES = −0.90). No cell-death pathway was significant in the proteome (Supplementary Table S5). Nuclear DNA damage, measured by RADF at 7 years, did not differ between genotypes in cortex or putamen (Supplementary Fig. S12A).

## 4. Discussion

### 4.1. Age-dependent molecular changes of a model with a non-expanding polyQ

The transgene in the TgHD minipig is a mutant partial human *HTT* cDNA under the control of the human *HTT* promoter. It contains a mixed CAG/CAA repeat and is expressed throughout the animal’s life. Because it lacks intron 1, it cannot produce the *HTT1a* transcript or protein (Osborne et al., 2026). Soluble mutant HTT was detected in the putamen and cortex at 4 and 8 years of age and did not increase consistently over that interval (Section 3.1.4, Supplementary Fig. S5).

Differences between the TgHD and WT putamen were strongly age-dependent. At 4 years neither the transcriptome nor the proteome differed much from WT. At 8 years 3,324 genes, 15% of those expressed, were differentially expressed in the transcriptome, while no individual protein differed significantly. Ageing alone does not explain this difference, because far fewer genes changed between the two ages in WT animals than in TgHD animals. To capture coordinated changes not detectable at the single-gene level, we applied GSEA with curated cell-type marker sets and standard pathway databases, and both proteome and transcriptome datasets showed many enriched pathways at 8 years (Fig. 2).

### 4.2. Loss of neuronal identity without evidence of cell death

We interpret the 8-year state of the TgHD putamen as a loss of neuronal transcriptional identity. Markers of D2 medium spiny neurons and of other neuronal types were reduced at 8 years in both the transcriptome and the proteome. The complete set of 1,836 genes downregulated at that age was enriched for targets of neuronal transcription factors, headed by MYT1L and followed by SCRT1, DPF1, NEUROD6, RORB and FOXG1 (Section 3.2.4, Supplementary Table S13). MYT1L maintains neuronal identity by repressing non-neuronal genes (Mall et al., 2017). *MYT1L*, *FOXG1* and *NEURODc* were themselves significantly downregulated at 8 years, although our data cannot say whether their reduction precedes or follows the loss of marker expression. This finding supports the hypothesis of a loss of neuronal identity and is consistent with the loss of neuronal transcriptional identity described in human and mouse HD striatum (Langfelder et al., 2016; Lee et al., 2020; Matsushima et al., 2023).

Bulk tissue cannot distinguish a loss of identity from a loss of the neurons themselves, because dead neurons that have already been cleared would produce the same loss of marker expression; this question needs single-nucleus or spatial methods. The small changes of individual marker proteins also argue against a pronounced loss of neurons, because cleared cells would be expected to lose their proteins together with their transcripts, whereas a transcriptional change in surviving neurons would reach the proteome later and more weakly. Apoptosis pathways were positively enriched at 8 years, but the neuronal apoptosis terms were not (Section 3.5.5), which supports a loss of identity without traces of neuronal death.

The negative enrichment of the D2 MSN marker set (NES = −3.02 in the transcriptome and −2.45 in the proteome) is notable because MSNs make up only about 15% of nuclei in the pig striatum (Li et al., 2026). This may be partly explained by the RNA content of neurons. Neuronal nuclei contain several times more RNA than glial nuclei, about five times in the human nucleus accumbens (Huuki-Myers et al., 2023; Sosina et al., 2021), and neurons are also generally larger cells, so they contribute more to the RNA of bulk tissue than their number suggests. Even so, an enrichment of that size implies a large change in the neurons, whether in their marker expression or in their number. Individual D2 MSN marker proteins changed less than their transcripts, yet the marker set was still significantly negatively enriched in the proteome.

At 8 years the D2 and the pan-MSN marker sets were negatively enriched in both datasets. The D1 set was negatively enriched in the transcriptome but not in the proteome, which matches the loss of D2 identity before D1 described by Matsushima et al. (2023).

Interneuron markers were also negatively enriched in both datasets, although interneurons are a smaller population and are usually described as relatively spared in early HD (Donaldson et al., 2026).

At 4 years the same D2 marker set was significantly positively enriched in the transcriptome, which is compatible with an early compensatory response. However, we found no report of the same phenomenon in the literature. It could be specific to the species, but with three animals per group we cannot make a definitive statement. The D1 set was significantly negatively enriched in the proteome at that age, so the two neuronal populations already differed at 4 years.

A second change at 4 years may be protective. Retrograde endocannabinoid signalling, which reduces glutamate release from corticostriatal terminals onto MSNs, was positively enriched in the 4-year transcriptome and negatively enriched at 8 years in both the transcriptome and the proteome. Endocannabinoid signalling protects neurons against excitotoxicity (van der Stelt et al., 2002), and cannabinoid receptors have a protective role in HD pathology as their downregulation in the basal ganglia is an early event in HD (Blázquez et al., 2011). The negative enrichment at 8 years in the TgHD putamen thus matches the early human loss, while the positive enrichment at 4 years has no reported counterpart in human HD. A transcriptional pattern attributed to excitotoxicity without overt neuronal loss has been reported in the OVT73 transgenic sheep model of HD at 5 years of age (Jiang et al., 2024; Jiang et al., 2023), and the glutamate neurotransmitter release cycle was positively enriched in our 4-year proteome. We therefore speculate that endocannabinoid signalling contributes to the minipig’s ability to compensate for the disease until a later age.

Among the D2 MSN genes, *SCN4B* had the largest transcript change between 4 and 8 years and is of particular interest, since overexpression of *SCN4B* was recently reported to improve the phenotype in HD mice (Sathitloetsakun et al., 2026). The widely used marker *PPP1R1B* appears less suitable for tracking progression in this model, since its change was limited to the transcript. *GPR88*, which is among the genes repeatedly reported in transcriptomic HD studies (reviewed in Malla et al., 2021), may be a better candidate, since its protein abundance was also reduced, although not significantly.

Oligodendrocyte markers were positively enriched in the transcriptome at both ages. Among the eight selected markers, the protease KLK6 has been reported to impair myelination in mice (Yoon et al., 2022), while the increase in *MBP* and *MOG* is more consistent with active myelination, and *APOD* encodes a stress-responsive lipid transporter with a proposed neuroprotective role in neurological disorders including Alzheimer’s disease (Dassati et al., 2014). Myelination defects and oligodendrocyte loss are well described in HD (Ferrari Bardile et al., 2023; Osipovitch et al., 2019), yet human post-mortem work reported a raised oligodendrocyte contribution in the striatum at early grades (Myers et al., 1991). Oligodendrocytes make up about half of striatal cells in pigs and humans, compared with about a quarter in mice (Li et al., 2026), so the pig may model the human oligodendrocyte contribution to the phenotype more closely than the mouse. Further experiments would be needed to explain why the oligodendrocyte markers were enriched only in the transcriptome and not in the proteome.

One possible explanation is that the loss of neuronal markers inflates the apparent oligodendrocyte proportion. This is unlikely here, because the oligodendrocyte markers were already significantly enriched at 4 years, when the D2 markers were also positively enriched. Myers et al. (1991) reported increased oligodendrocyte numbers alongside reduced neuronal numbers and no astrogliosis in Vonsattel grades 0 to 2. We measured marker expression and not cell numbers, so the two cannot be equated, but both are increases of the oligodendrocyte component, and that places the 8-year pig closer to the early grades of the human disease than to late HD.

### 4.3. Opposite enrichment of mitochondrial and oxidative phosphorylation pathways at 4 and 8 years

We interpret the mitochondrial changes as an early compensatory increase followed by a later decline of mitochondrial gene and protein expression. At 4 years mitochondrial and oxidative phosphorylation pathways were positively enriched in both the transcriptome and the proteome, with no individual pathway standing out (Fig. 5). This could be a compensatory increase in mitochondrial content in response to the impaired energy metabolism of cells expressing mutant HTT (Seong et al., 2005). Such an interpretation is supported by the higher mitochondrial DNA copy number previously reported in the basal ganglia of TgHD minipigs (Askeland et al., 2018a).

At 8 years the same pathways were negatively enriched. This partly resembles late human HD, where mitochondrial mass and the subunits of respiratory complexes III and IV are reduced together with the biogenesis regulators PGC-1α and TFAM (Cui et al., 2006; Kim et al., 2010; Squitieri et al., 2010). *PPARGC1A* expression was reduced in our 8-year TgHD animals, while its protein, PGC-1α, was not detected in the proteome. However, mitochondrial DNA copy number was not lower in TgHD animals at 7 years, which does not support a pronounced loss of mitochondria. This biphasic pattern, positive enrichment at 4 years and negative at 8 years, agrees with HD mouse models, in which an early increase and a later decrease of mitochondrial gene or protein expression in the striatum were reported (Andrzejewski et al., 2014; Garcia et al., 2022; Lopes et al., 2022).

Complex I showed the biphasic pattern most clearly. Its subunit set was significant in all four datasets, positively enriched at 4 years and negatively enriched at the later age in TgHD (Fig. 5B), and *NDUFA5* was one of only two OXPHOS subunit genes significantly reduced at 8 years. Complex I activity is also increased in a transgenic HD mouse and in mice treated with the complex II inhibitor 3-nitropropionic acid, where it is interpreted as a compensatory mechanism (Oláh et al., 2008), and oxidative phosphorylation proteins are more abundant in the early-stage R6/2 mouse (Zabel et al., 2009). The decrease at 8 years would then be the loss of this compensation, and it is supported by the reduction of *PPARGC1A*.

The pentose phosphate pathway (PPP) was positively enriched in the 7- to 8-year proteome, at the age when oxidative phosphorylation pathways were negatively enriched (Supplementary Fig. S13A). A change in glucose metabolism away from oxidative phosphorylation could explain this (Del Prado et al., 2024; Garcia et al., 2022). Unlike the PPP, the glycolysis gene sets were not positively enriched. Glycolysis was negatively enriched in TgHD relative to WT in the 8-year transcriptome and not significant in the proteome, which agrees with the selective reduction in striatal glycolysis reported in patients, alongside unchanged oxidative metabolism (Powers et al., 2007). In the 8-year transcriptome both oxidative and glycolytic pathways were negatively enriched, which could reflect reduced metabolic capacity or energy demand. We would need to measure respiration in this tissue directly, since expression data alone cannot establish a bioenergetic deficit. We also found no sign of increased mitochondrial DNA damage in the 108 bp *MT-RNR1* sequence measured by RADF, although damage elsewhere in the mitochondrial genome cannot be excluded.

Calcium-handling pathways, like the mitochondrial pathways, were positively enriched at 4 years and negatively enriched at 8 years (Supplementary Fig. S13B). Impaired mitochondrial calcium handling is described in HD models (Milakovic et al., 2006; Panov et al., 2002). In our data the most negative calcium pathways were neuronal channel complexes in the transcriptome. In contrast, mitochondrial calcium transport was negatively enriched in the proteome only, so the calcium result may reflect the reduction of neuronal transcripts as much as altered mitochondrial calcium handling.

In the proteome the ribosome sets also reversed between the two ages. The KEGG ribosome pathway was positively enriched in the transcriptome at both ages, while in the proteome it was positively enriched at 4 years and negatively enriched at 7 to 8 years (Fig. 2C), and the mitochondrial ribosome set was the most negative MitoCarta set in the 7- to 8-year proteome (Section 3.3.1). Higher transcript and lower protein abundance of ribosomal proteins at the older age could point to reduced synthesis of ribosomal proteins, for example through ribosome stalling by mHTT, as reported in mouse HD striatal neuronal cells (Eshraghi et al., 2021).

### 4.4. A microglial response and a hypothesis of synapse pruning

Our data are consistent with a microglia-centred inflammatory response. The broad positive enrichment of immune and inflammatory pathways at 8 years (Fig. 6A) and the coordinated increase in homeostatic and general microglial markers may reflect microgliosis. The increase in *IRF5* (log2FC = +1.30) and in the complement genes *C1ǪA* and *C1ǪC* indicates a change in microglial state in addition to any potential increase in cell number (Fig. 6B and C). The homeostatic genes increased together with part of the DAM genes, which fits a larger number of microglia, or more transcription per cell, better than a transition to the DAM state, in which the homeostatic genes decrease (Paolicelli et al., 2022). The transcription factor profile of the genes upregulated at 8 years was dominated by transcription factors of microglia, macrophages and other immune cells, headed by TFEC, IRF8 and PU.1, encoded by *SPI1* (Fig. 6D). Such a profile may be linked to a cell-autonomous inflammatory response through PU.1 and C/EBP, which was described as intrinsically triggered by mHTT (Crotti et al., 2014).

Toll-like receptor and NF-κB signalling were positively enriched at 8 years. Toll-like receptor signalling and the MyD88-independent (TRIF) TLR4 cascade were significantly enriched in the 7- to 8-year proteome, and canonical NF-κB signalling, the downstream node of the MyD88-dependent branch, in the 8-year transcriptome (Fig. 6A). In human HD, TLR4 protein is increased in the putamen and *TLR4* and *TREM2* variants modify clinical progression (Vuono et al., 2020). In contrast, we detected neither protein in our proteome, and both transcripts increased but not significantly, so we can neither confirm nor exclude that finding.

In human and mouse HD, microglia prune complement-tagged corticostriatal synapses before neuronal loss (Saba et al., 2022; Wilton et al., 2023), and we propose that a similar process may operate in the TgHD minipig. Two signals that control pruning were changed at 8 years. First, transcripts of the complement components that tag synapses for microglial engulfment (C1q, C3) were increased (Fig. 6C). Complement inhibitors did not decrease, and *CD4c* was upregulated, so the data do not show a release of complement inhibition. Second, the transcript of the neuronal chemokine fractalkine (*CX3CL1*), whose signalling to microglia has been linked to synapse pruning (Saba et al., 2022), was reduced, while the transcript of its microglial receptor (*CX3CR1*) was increased. In bulk tissue these two changes may partly reflect fewer neuronal transcripts and more microglia. If they reflect regulation, more CX3CR1 protein could either compensate for less CX3CL1 or itself promote engulfment. In the proteome CX3CL1 was unchanged and CX3CR1 was not detected. Together with the possible microgliosis and the transcription factor profile described above, more complement tagging and less fractalkine signalling from neurons could explain the reduction in synaptic genes and proteins, even though neurons are a minority of striatal cells. Synapse removal may also contribute to the decline in D2 markers described above while the neurons themselves survive.

By contrast, we found little evidence (Fig. 6E) of the classical reactive astrogliosis of late-stage human HD (Vonsattel et al., 1985). *GFAP* and *VIM* were unchanged, and only the inflammation-induced set of reactive astrocyte markers was enriched. This agrees with astrocyte data from HD mouse models and human caudate, in which marked astrocyte reactivity appeared only at neuropathological grade 3 and was absent at grades 0 and 1 (Diaz-Castro et al., 2019). *CP*, which encodes the ferroxidase ceruloplasmin and is one of the two reactivity genes with the largest increase in those human data, was also increased in the TgHD putamen, which may relate to the iron accumulation reported in the basal ganglia of HD patients (Liang et al., 2025). This asymmetry supports an inflammatory response centred on microglia rather than on astrocytes in the TgHD minipig putamen at 8 years.

### 4.5. Altered splicing and DNA-repair gene changes as possible downstream events

Recent work in cell and mouse models of HD detected splicing dysregulation before symptom onset (Elorza et al., 2021; Tano et al., 2023). Here we found little change in alternative splicing at 4 years and many more changes at 8 years, both between TgHD and WT animals and between 8-year and 4-year TgHD animals (Fig. 7A). If altered splicing were an intrinsic effect of mutant HTT, we would expect it to be present already in the TgHD versus WT comparison at 4 years.

A direct effect, for example the sequestration of splicing factors by expanded CAG repeat RNA reported in cell and human studies (Sathasivam et al., 2013; Schilling et al., 2019), should leave a footprint, the binding motifs of the sequestered RBPs enriched around the affected exons. When we tested RBP motifs around skipped exons, only a few motif families were enriched, and only at 8 years. Most of them were short, of low complexity or shared between proteins, and could not be attributed to one protein (Section 3.5.3). The exception was the motif of RBFOX, a family of splicing regulators of which RBFOX1 and RBFOX3 (NeuN) are neuronal. It was enriched in the intron downstream of exons that lost inclusion, the position from which RBFOX proteins promote exon inclusion (Lovci et al., 2013; Yeo et al., 2009), which is consistent with reduced RBFOX activity. Indeed, transcripts of all three RBFOX genes were reduced at 8 years but not at 4 years, while RBFOX1 and RBFOX3 proteins were unchanged and RBFOX2 protein was not detected (Fig. 7D and E).

The one attributable motif thus belongs to a regulator that is itself neuronal. We therefore propose that the altered splicing is a downstream consequence of a change in cell-type composition or of a loss of neuronal identity. Neurons have a splicing programme distinct from that of other striatal cell types (Gokce et al., 2016), and the differentially spliced genes were mostly neuronal and cytoskeletal. They were enriched for vesicle transport and endocytosis at the synapse, processes in which huntingtin itself is involved (Barron et al., 2021). Elorza et al. (2021) used only the splicing changes shared between human post-mortem striatum and early symptomatic R6/1 mouse striatum to filter out changes driven by altered neuronal and glial composition and pro-inflammatory signals. An HD-specific set of splicing changes remained after that intersection, and even though our design cannot provide the same separation, three of the six effector genes defined by Elorza et al. (2021), namely *SYNJ1*, *KCTD17* and *VPS13C*, were differentially spliced at 8 years in the TgHD minipig.

Ciliary genes and pathways were prominent among the differentially spliced genes at both ages, and the ciliary gene sets were also the main source of disagreement between the transcriptome and the proteome, with negative NES in the 8-year transcriptome and positive NES in the 7- to 8-year proteome (Supplementary Fig. S4C). Ciliary gene regulation in this model therefore differs from WT in splicing, transcript abundance and protein abundance. This is notable given the sperm defects described in this model (Macakova et al., 2016) and the role of huntingtin in ciliogenesis (Keryer et al., 2011), and it warrants further investigation.

The CAG/CAA repeat of this model is not expected to expand and we detected no nuclear DNA damage, yet several DNA-repair pathways were positively enriched at 8 years (Fig. 8B). We therefore examined the individual genes in these pathways, including those identified as genetic modifiers of age at onset in HD. *ATM*, *FEN1*, *MUTYH*, *PNKP* and *MLH1* were upregulated; *NEIL2* and *POLB* were downregulated; and *MSH3* and *FAN1* were unchanged (Fig. 8A). The increase in *FEN1* is the opposite of the low-FEN1 state that favours striatal expansion in HD mice (Goula et al., 2009). Ooi et al. (2019) also found stable repeats together with an active DNA damage response in an isogenic panel of human pluripotent stem cell lines, and iPSCs derived from this TgHD minipig have *FAN1* upregulated without detectable DNA damage (Rysankova et al., 2026). However, in the adult bulk putamen, we did not detect changes in *FAN1* expression.

We see two possible interpretations of the DNA-repair enrichment. The first is a general genotoxic or cellular stress response, since ATM has a broad role and not all repair genes increased in expression. Nuclear DNA damage was not increased in the TgHD cortex or putamen at 7 years (Supplementary Fig. S12A), hence the repair pathways were enriched without detected damage. However, the assay would not report a small number of lesions or breaks that are repaired quickly. Mutant HTT is reported to interact with both ATM and PNKP, associating with ATM in a DNA-repair complex (Maiuri et al., 2017) and reducing PNKP activity without changing PNKP protein levels (Gao et al., 2019). We therefore cannot interpret the transcript changes as higher repair capacity.

The second interpretation is cell proliferation. DNA replication and cell-division pathways were also strongly positively enriched at 8 years (Fig. 8B), and the excision-repair pathways share many genes with DNA replication, including the genes of the replicative polymerases as well as *PCNA* and *FEN1*. Part of the DNA-repair enrichment may therefore reflect dividing cells and not a response to DNA damage. Proliferating microglia are the explanation best supported by the rest of our data, since the immune pathways were enriched at the same age and the microglial markers were increased; cells of the oligodendrocyte lineage may contribute as well. However, we cannot exclude a contribution of other cells, including neurons. Aberrant cell-cycle re-entry in neurons has been proposed as a route to cell death in HD (Liu et al., 2015). Oxidative stress (Klein and Ackerman, 2003) or DNA damage can push quiescent neurons from G0 into G1 and S phase, after which they undergo apoptosis rather than divide (Kruman et al., 2004). This would be consistent with the increase in *ATM* expression and with the positive enrichment of apoptosis pathways at 8 years, although the neuronal apoptosis terms were not enriched (Section 3.5.5).

### 4.6. What a non-expanding, fragment-based TgHD minipig can and cannot model

Taken together, our data describe two different states at the two ages. At 4 years the TgHD putamen showed only modest changes from WT. The changes we highlight at 4 years, in D2 MSN markers, mitochondrial pathways and retrograde endocannabinoid signalling, were all increases, which we interpret as a very early, compensated state. At 8 years the same markers and pathways were reduced, and immune changes, altered splicing and changes in DNA-repair gene expression appeared. This corresponds to Stage 1 of the Huntington’s Disease Integrated Staging System, biomarker evidence of pathogenesis without clinical signs (Tabrizi et al., 2022). We cannot tell whether the 8-year minipig would already be in Stage 2, which adds definite motor or cognitive signs, since we did not measure them in these animals. With two time points we cannot say when the change starts or which molecular changes appear first, but the order suggests that the neuronal and mitochondrial changes come before the others.

The genetic design of this model excludes three molecular mechanisms of human HD pathophysiology. The transgene is an *HTT* cDNA lacking intron 1, so it cannot produce the *HTT1a* transcript or the highly toxic HTT1a protein (Osborne et al., 2026). The CAG repeat is mixed with CAA units throughout, so it does not expand somatically, and its transcript does not form the slippery hairpins described for pure CAG tracts (Sobczak and Krzyzosiak, 2005). This may seem a disadvantage, but it offers an opportunity to isolate the molecular consequences of expressing an N-terminal mHTT fragment from the other disease mechanisms.

The most important of these excluded mechanisms is somatic expansion, which is now understood to be a major driver of progression in human HD. The inherited repeat expands over decades, and the slow molecular dysregulation that it initiates precedes the loss of neuronal identity and the cell death that follow once the repeat exceeds roughly 150 units in vulnerable striatal neurons (Handsaker et al., 2025; Mätlik et al., 2024). The TgHD minipig has a stable repeat, so it can model only the earliest part of this process. From this perspective, the mildness of the phenotype is not a shortcoming of the model but a predictable consequence of its genetic design. The animal stays in the premanifest stage that precedes the critical expansion threshold in human medium spiny neurons, which is arguably the most relevant window for therapeutic intervention. This is also why the model has served preclinical gene-therapy studies (Evers et al., 2018).

Because the model lacks an easily assessed behavioural phenotype, molecular readouts are needed to follow its progression. The molecular description provided here can serve to select candidate markers in cerebrospinal fluid, which can be collected in minipigs (Evers et al., 2018).

### 4.7. Limitations

The RNA sequencing cohort was three animals per group, which limits the power to detect small transcriptional differences, particularly at 4 years, so the contrast between the two ages may appear larger than it is. The sequencing results were supported in an independent cohort of 7-year-old animals, in which 21 genes measured by qPCR were moderately concordant with the 8-year RNA-seq data (r = 0.65, P = 0.0015, Supplementary Fig. S4B). Bulk sequencing and proteomics cannot assign a change in expression or abundance to a cell type, cannot separate cell number from expression per cell, and cannot demonstrate synapse removal, which requires single-nucleus or spatial methods. Transcripts removed by nonsense-mediated decay are invisible to steady-state RNA sequencing, so splicing changes that produce such transcripts are probably undercounted. The incomplete annotation of the *Sus scrofa* genome and proteome limits the confirmation of some transcript changes at the protein level. Transcripts of the mitochondrial genome were not quantified, so the OXPHOS result at the RNA level is based on nuclear-encoded subunits only, while the proteome includes 9 of the 13 mitochondrially encoded subunits. The design was cross-sectional, with two ages and different animals at each age, and no assay in this study tested aggregation of the transgenic protein. This would require immunohistochemistry, since N171-82Q mice, which also have a cDNA transgene and produce no HTT1a, show diffuse nuclear HTT aggregation (Osborne et al., 2026). The 4-year and 8-year animals were not perfused before dissection, so blood cells contribute to these samples, in both genotypes alike. Finally, the interpretation depends on the cell-type marker sets curated from the literature and on the grouping of pathways into biological themes. We document both with their sources in Supplementary Tables S7 and S8 and in the analysis code.

## 5. Conclusions

We characterised the putamen of the TgHD minipig at 4 and 8 years by transcriptomics and proteomics, and found that the model changes substantially between those two ages. At 4 years the TgHD putamen showed only modest changes in both datasets. At 8 years it showed widespread transcriptomic change, a reversal of mitochondrial and oxidative phosphorylation pathway enrichment, reduced markers of neuronal identity and increased expression of microglial marker genes. We propose that microglial engulfment of synapses links the microglial changes to the loss of synaptic and neuronal identity markers, which could explain a molecular phenotype without evidence of neuronal death. The splicing changes were about five times as frequent at 8 years as at 4 years and are consistent with reduced RBFOX transcript abundance, so they are more likely a downstream consequence of altered cell composition than a primary effect of mutant huntingtin. None of this required repeat expansion, which makes the TgHD minipig a model of the premanifest rather than the late stage of the disease.

## CRediT authorship contribution statement

**Eva Kamenná:** Conceptualization, Data curation, Formal analysis, Investigation, Methodology, Project administration, Software, Validation, Visualization, Writing – original draft, Writing – review & editing. **Kateryna Nemesh:** Conceptualization, Data curation, Funding acquisition, Investigation, Methodology, Writing – review & editing. **Jakub Červenka:** Data curation, Investigation, Methodology, Writing – review & editing. **David Sekáč:** Formal analysis, Methodology, Writing – original draft. **Christian Landles:** Data curation, Funding acquisition, Investigation, Methodology, Project administration, Resources, Supervision, Writing – review & editing. **Anastasiia Kolesnikova:** Writing – review & editing. **Ivona Valeková:** Writing – original draft, Writing – review & editing. **Štefan Juhás:** Investigation, Resources. **Jana Juhásová:** Investigation, Resources. **Georgina Osborne:** Data curation, Investigation, Methodology. **Helena Kupcová Skalníková:** Funding acquisition, Resources, Supervision, Writing – review & editing. **Petr Vodička:** Validation, Writing – review & editing. **Jan Motlík:** Funding acquisition, Project administration. **Jiří Klempíř:** Funding acquisition, Writing – review & editing. **Zdeňka Ellederová:** Conceptualization, Funding acquisition, Supervision, Writing – review & editing. **Jiří Klíma:** Conceptualization, Supervision, Writing – review & editing.

## Acknowledgements

We thank Mgr. Lucie Hošková for help with data visualisation and Prof. Gillian Bates (University College London) for her comments on the manuscript.

Computational resources were provided by the e-INFRA CZ project (ID:90254), supported by the Ministry of Education, Youth and Sports of the Czech Republic.

## Ethics

All animal experiments were conducted in accordance with Directive 2010/63/EU, Czech Act No. 246/1992 Coll., on the Protection of Animals Against Cruelty, and Decree No. 419/2012 Coll., under approved project protocols No. 109/2020 and 53/2015. The Institute of Animal Physiology and Genetics of the Czech Academy of Sciences is authorised by the Czech Ministry of Agriculture to conduct animal research (Authorisation No. MZE-24154/2021-18134).

## Funding

This work was supported by grants from the CHDI Foundation (ZE, CL), the Czech Academy of Sciences, Strategy AV21 (VP29), the Ministry of Education, Youth and Sports of the Czech Republic, grant EXREGMED, Excellent Research in Regenerative Medicine (CZ.02.01.01/00/22_008/0004562), and by the Innovative Health Initiative Joint Undertaking (IHI JU) under grant agreement No 101165643. The JU receives support from the European Union’s Horizon Europe research and innovation programme and COCIR, EFPIA, EuropaBio, MedTech Europe and Vaccines Europe, and Breakthrough T1D and VeriSIM Life.

K.N. was supported by Institutional Research Concept (IGA)[RVO67985904]. J.C. and H.K.S. were supported by the Ministry of Education, Youth and Sports project (LX22NPO5102) funded by the European Union – Next Generation EU, Ministry of Health project (NW26A-NUVR) and Charles University Cooperatio program Oncology and Haematology. E.K. and J. Klempíř were supported by the EU Joint Programme – Neurodegenerative Disease Research (delCAA-HD Project, JPND2023-1822-096) and the European Reference Network for Rare Neurological Diseases (Project ID No. 739510).

Disclaimer. Funded by the European Union, the private members, and those contributing partners of the IHI JU. Views and opinions expressed are however those of the author(s) only and do not necessarily reflect those of the aforementioned parties. Neither of the aforementioned parties can be held responsible for them.

## Declaration of competing interest

The authors declare that they have no known competing financial interests or personal relationships that could have appeared to influence the work reported in this paper.

## Data availability

The RNA sequencing data, the mass spectrometry proteomics data and the analysis scripts will be made publicly available after peer review. The processed results are provided in Supplementary Tables S1 to S18.

## Supplementary material

Supplementary material associated with this article consists of Supplementary Figures S1 to S19 with their captions, in one file, and Supplementary Tables S1 to S18, one spreadsheet file each.

## Supplementary figures

- Supplementary Fig. S1. Principal component analysis of the transcriptomic and proteomic data.
- Supplementary Fig. S2. Volcano plots for the five differential expression comparisons.
- Supplementary Fig. S3. Volcano plots for the proteomics comparisons.
- Supplementary Fig. S4. Validation of the transcriptomic data by qPCR and cross-platform correlation.
- Supplementary Fig. S5. Soluble mutant huntingtin measured by homogeneous time-resolved fluorescence.
- Supplementary Fig. S6. Gene-level changes mapped onto the KEGG Huntington disease pathway.
- Supplementary Fig. S7. Distribution of gene-level fold changes within each cell-type marker set.
- Supplementary Fig. S8. D1 medium spiny neuron candidate genes between 4 and 8 years.
- Supplementary Fig. S9. Mitochondrial pathway enrichment across all pathway databases.
- Supplementary Fig. S10. Mitochondrial pathway enrichment of MitoCarta 3.0 gene sets.
- Supplementary Fig. S11. Changes in the oxidative phosphorylation subunit genes.
- Supplementary Fig. S12. Nuclear and mitochondrial DNA damage and mitochondrial DNA copy number at 7 years.
- Supplementary Fig. S13. Carbohydrate metabolism and calcium-handling pathway enrichment.
- Supplementary Fig. S14. Immune and inflammatory pathway enrichment across all pathway databases.
- Supplementary Fig. S15. Toll-like receptor and NF-κB pathway genes.
- Supplementary Fig. S16. Direction of the alternative splicing changes.
- Supplementary Fig. S17. Positional RNA-binding protein motif enrichment around skipped exons at 8 years.
- Supplementary Fig. S18. Positional motif enrichment for SF1 and QKI around skipped exons at 8 years.
- Supplementary Fig. S19. Splicing and RNA-processing pathway enrichment.

## Supplementary tables

- Supplementary Table S1. Sample overview, animals, ages, sex, perfusion.
- Supplementary Table S2. Differential gene expression (DESeq2), the five comparisons.
- Supplementary Table S3. RNA-seq count matrix, raw and variance-stabilised, with sample information.
- Supplementary Table S4. Proteomics differential abundance (limma, age-normalised), 4-year, 8-year and 7- to 8-year comparisons.
- Supplementary Table S5. Gene set enrichment analysis, all databases and comparisons.
- Supplementary Table S6. Cell-type marker set enrichment.
- Supplementary Table S7. Pathway metadata and theme assignment.
- Supplementary Table S8. Cell-type marker gene sets, provenance and scoring.
- Supplementary Table S9. Alternative splicing events (rMATS).
- Supplementary Table S10. RNA-binding protein positional motif enrichment.
- Supplementary Table S11. Validation correlations, qPCR against RNA-seq and RNA against protein.
- Supplementary Table S12. Gene annotation and provenance.
- Supplementary Table S13. ChEA3 transcription factor enrichment.
- Supplementary Table S14. R package versions.
- Supplementary Table S15. Leading-edge gene overlap between pathways.
- Supplementary Table S16. Over-representation analysis of alternatively spliced genes.
- Supplementary Table S17. qPCR primer sequences.
- Supplementary Table S18. Two-way ANOVA statistics for the HTRF assays of Supplementary Fig. S5.

## Notes

### Competing Interest Statement

The authors have declared no competing interest.

