## Supplementary figures for "Age-dependent changes in the putamen transcriptome and proteome of a transgenic minipig model of Huntington’s disease"

##### Supplementary Figure S1

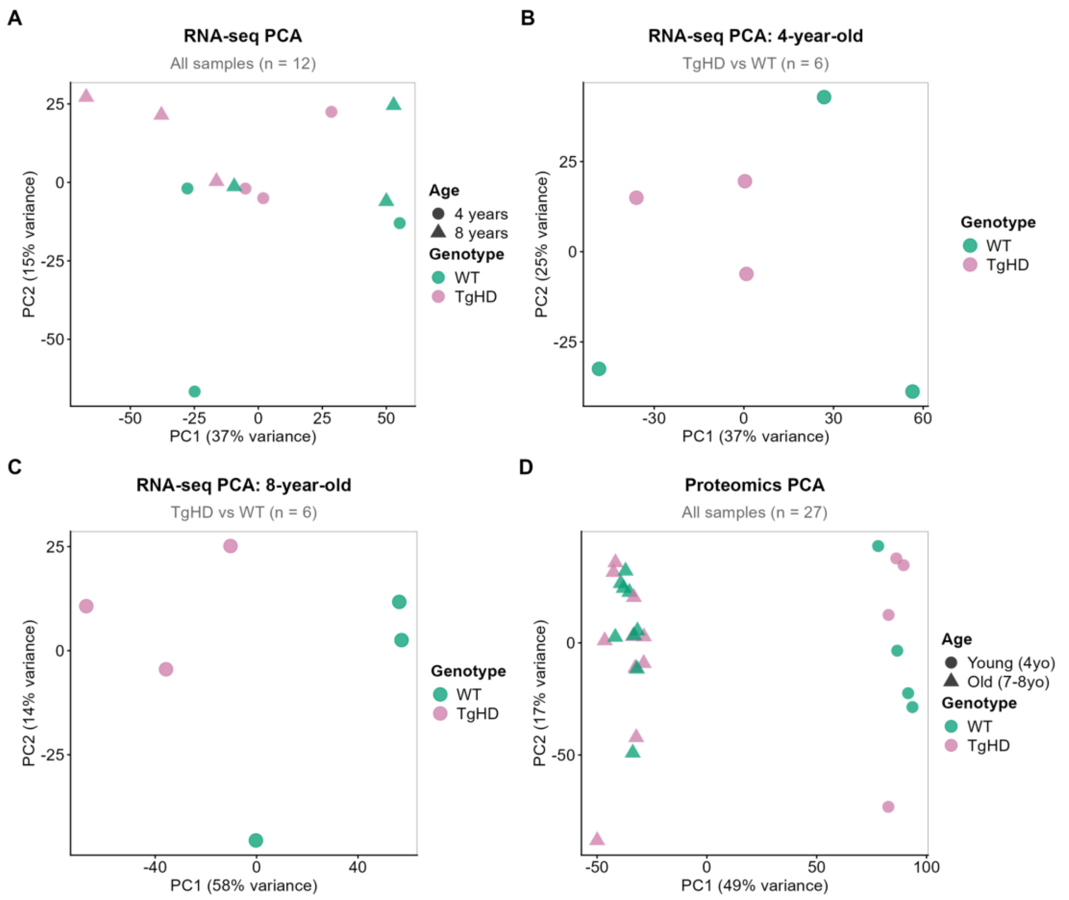


**Supplementary Fig. S1. Principal component analysis of the transcriptomic and proteomic data.** PCA of variance-stabilised RNA-seq counts for **(A)** all sequenced animals, **(B)** the 4-year animals and **(C)** the 8-year animals, and **(D)** of age-normalised protein abundances for all animals with proteomics. Each point is one animal, coloured by genotype and, in (A) and (D), shaped by age group. Abbreviations. PC, principal component. PCA, principal component analysis. TgHD, transgenic Huntington’s disease minipig. WT, wild type.

#### Supplementary Figure S2

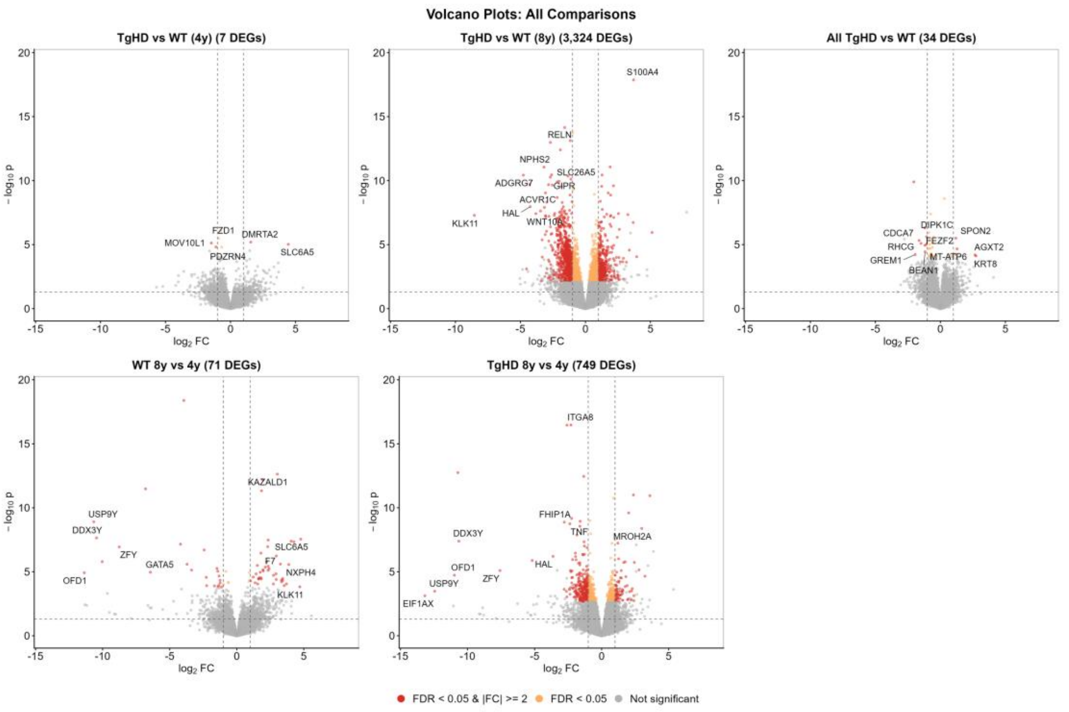


**Supplementary Fig. S2. Volcano plots for the five differential expression comparisons.** Volcano plots of log2 FC against minus log10 nominal p value, on shared axes, for TgHD versus WT at 4 years, TgHD versus WT at 8 years, TgHD versus WT with both ages pooled, WT at 8 versus 4 years and TgHD at 8 versus 4 years. The number of DEGs, those with a FDR below 0.05, is written in each panel title. The dashed lines mark a nominal p value of 0.05 and log2 fold changes of plus and minus 1. Labelled genes are those with FDR below 0.05 and the largest product of absolute log2 fold change and minus log10 p, up to five in the first panel and up to ten in each of the others. The first two comparisons are also shown in Fig. 2B, there with an additional curated set of labelled genes. Abbreviations. DEG, differentially expressed gene. FC, fold change. FDR, false discovery rate. TgHD, transgenic Huntington’s disease minipig. WT, wild type.

### Supplementary Figure S3

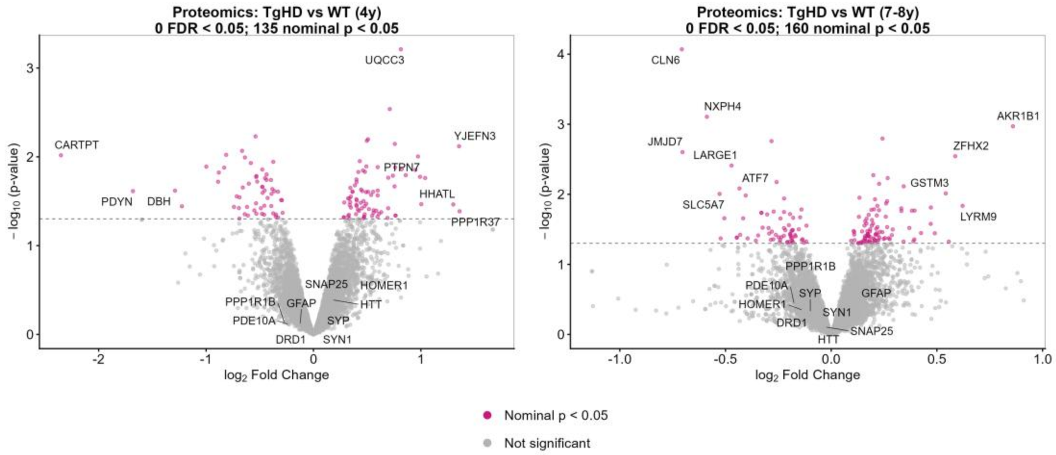


**Supplementary Fig. S3. Volcano plots for the proteomics comparisons.** Volcano plots of log2 fold change against minus log10 nominal p value for the age-normalised proteomics comparisons, TgHD versus WT at 4 years and at 7 to 8 years. The dashed line marks a nominal p value of 0.05. No protein reached a FDR below 0.05 in either comparison, and the number of proteins with a nominal p below 0.05 is written in each panel title. Labelled proteins are those with a nominal p below 0.05 and the largest product of absolute log2 fold change and minus log10 p, up to eight at 4 years and up to ten at 7 to 8 years, together with a fixed set of Huntington disease related proteins wherever quantified (HTT, PPP1R1B, DRD1, DRD2, GFAP, BDNF, HOMER1, PDE10A, SNAP25, SYN1, SYP). Abbreviations. FDR, false discovery rate. TgHD, transgenic Huntington’s disease minipig. WT, wild type.

#### Supplementary Figure S4

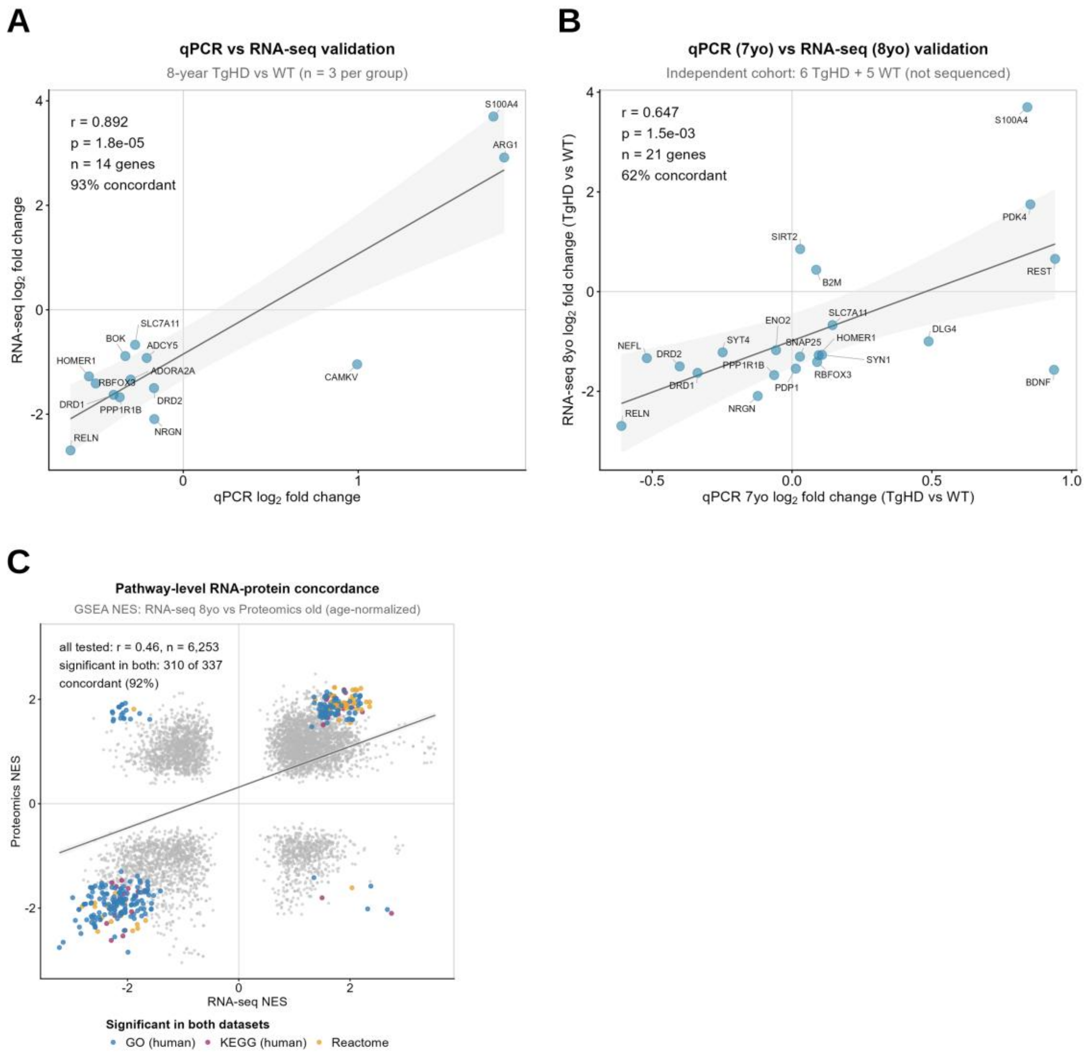


**Supplementary Fig. S4. Validation of the transcriptomic data by qPCR and cross-platform correlation. (A)** Quantitative polymerase chain reaction (qPCR) log2 fold change against RNA-seq log2 fold change for 14 genes measured on RNA re-isolated from the same 8-year animals. **(B)** As (A), for 21 genes measured in an independent cohort of 7-year animals that were not sequenced, plotted against the 8-year RNA-seq values. **(C)** NES from gene set enrichment analysis of the 8-year transcriptome against the age-normalised proteome at 7 to 8 years, one point per pathway, for the 6,253 human ortholog KEGG, Gene Ontology and Reactome pathways tested in both datasets (grey), with the 337 pathways significant in both coloured by source database. In (A) and (B) each point is one gene, and concordance is the share of genes increased in both datasets or decreased in both. In all panels the grey line is the linear fit with its 95% confidence band. Correlation coefficients, p values and the concordance counts are in Supplementary Table S11 and primer sequences in Supplementary Table S17. Abbreviations. NES, normalised enrichment score. qPCR, quantitative polymerase chain reaction. TgHD, transgenic Huntington's disease minipig. WT, wild type.

#### Supplementary Figure S5

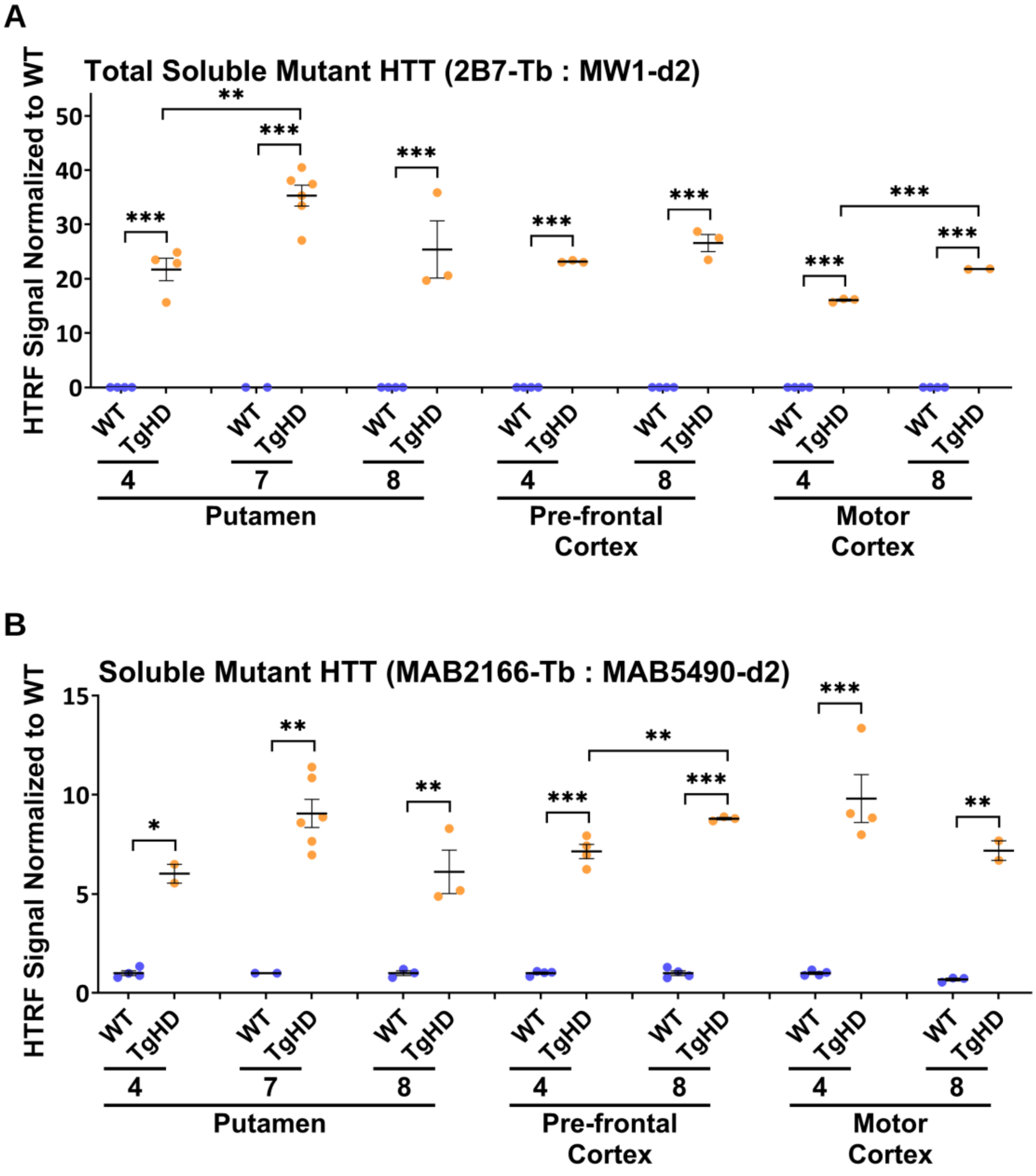


**Supplementary Fig. S5. Soluble mutant huntingtin measured by homogeneous time-resolved fluorescence.** HTRF assays in putamen (4, 7 and 8 years), pre-frontal cortex and motor cortex (4 and 8 years) of WT and TgHD animals. **(A)** Total soluble mutant huntingtin assay (2B7-Tb with MW1-d2), in which MW1 recognises the expanded polyglutamine tract. **(B)** Transgene-specific assay (MAB2166-Tb with MAB5490-d2). The MAB2166 epitope (amino acids 443 to 457) detected the transgenic human HTT protein only, and not the endogenous *Sus scrofa* HTT sequence. The signal was normalised to WT within each tissue and age (n = 2 to 6 per genotype and age). Data analysis was by two-way ANOVA with Bonferroni post hoc correction, and each data point represents one individual minipig. The test statistic, degrees of freedom and P values for the ANOVA are reported in Supplementary Table S18. Error bars are mean ± SEM. * P ≤ 0.05, ** P ≤ 0.01 and *** P ≤ 0.001. Brackets join the groups compared. Abbreviations. ANOVA, analysis of variance. HTRF, homogeneous time-resolved fluorescence. HTT, huntingtin. SEM, standard error of the mean. Tb, terbium cryptate. TgHD, transgenic Huntington's disease minipig. WT, wild type.

#### Supplementary Figure S6

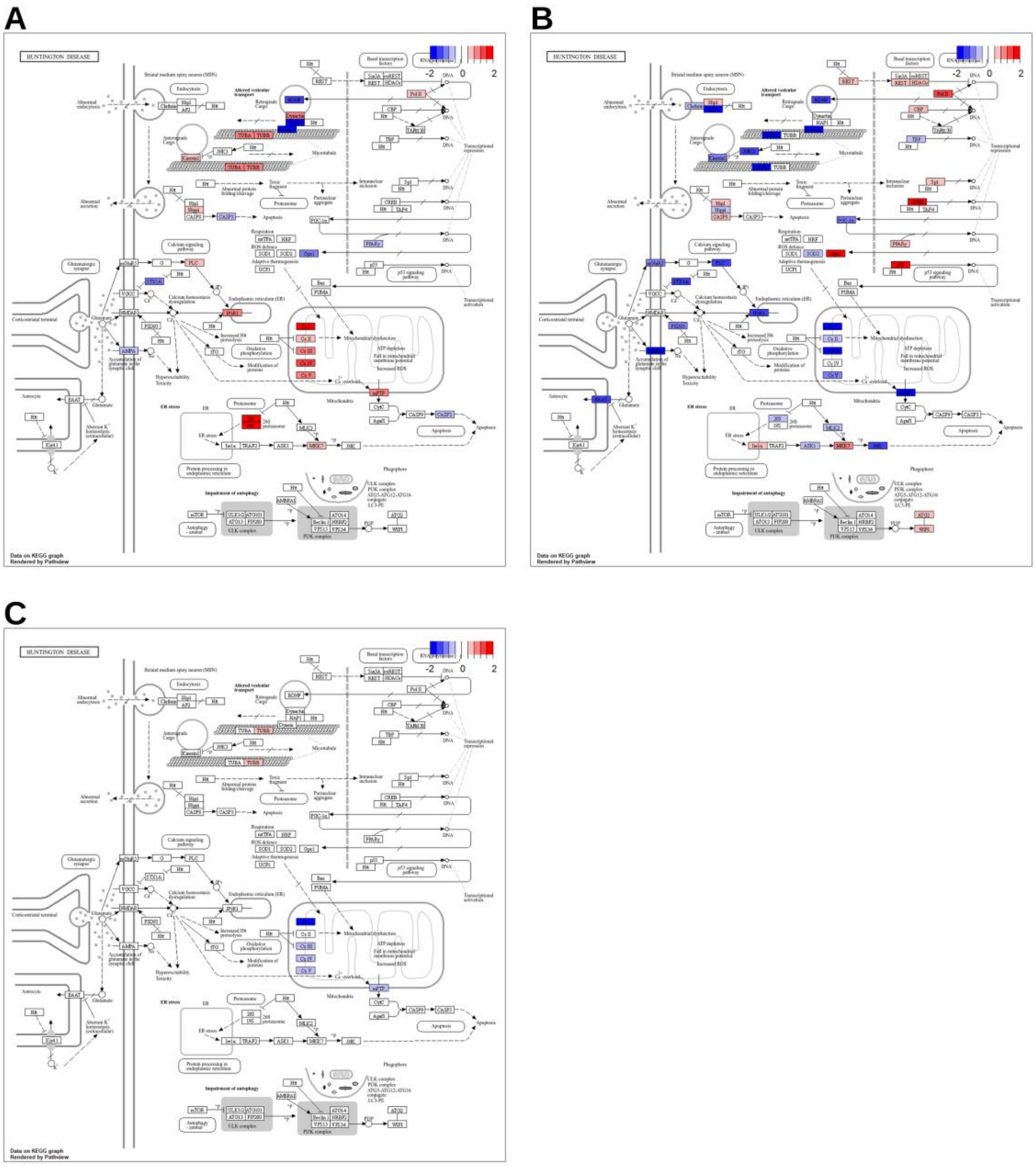

**Supplementary Fig. S6. Gene-level changes mapped onto the KEGG Huntington disease pathway.** Pathview rendering of the KEGG Huntington disease pathway (hsa05016) for **(A)** RNA-seq at 4 years, **(B)** RNA-seq at 8 years and **(C)** age-normalised proteomics at 7 to 8 years, TgHD relative to WT. Node colour is the log2 fold change from blue (negative) through white to red (positive), capped at plus and minus 2 (colour bar). Pathway members not detected in that data set are white. Abbreviations. TgHD, transgenic Huntington's disease minipig. WT, wild type.

#### Supplementary Figure S7

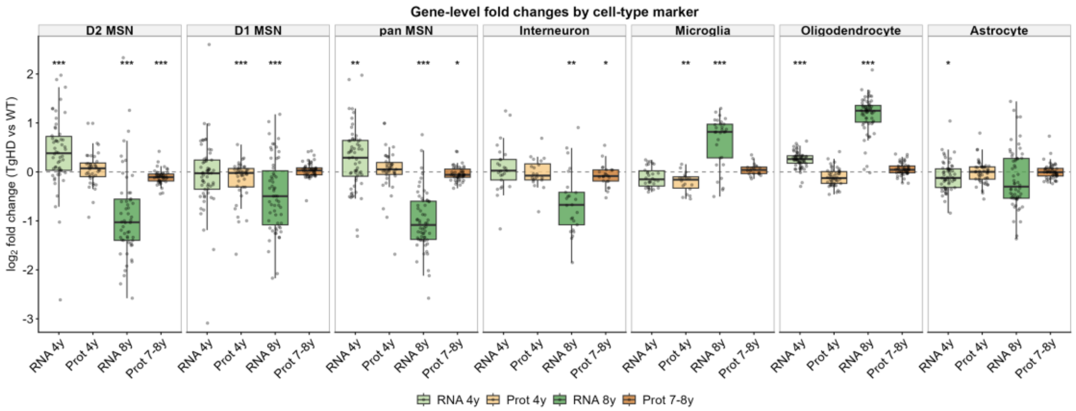


**Supplementary Fig. S7. Distribution of gene-level fold changes within each cell-type marker set.** Box plots of gene-level log2 fold change, TgHD relative to WT, for the curated cell-type marker gene sets of Supplementary Table S8. Boxes show the median and interquartile range, whiskers extend to 1.5 times the interquartile range, and every gene is drawn as a point. Asterisks above a box mark the significance of that gene set in GSEA, * FDR below 0.05, ** below 0.01, *** below 0.001. Abbreviations. FDR, false discovery rate. GSEA, gene set enrichment analysis. MSN, medium spiny neuron. TgHD, transgenic Huntington's disease minipig. WT, wild type.

#### Supplementary Figure S8

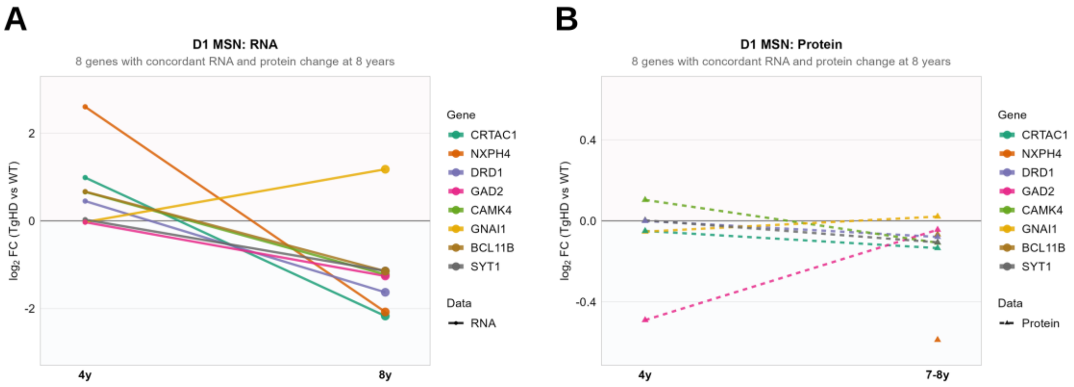

**Supplementary Fig. S8. D1 medium spiny neuron candidate genes between 4 and 8 years.** Log2 FC, TgHD relative to WT, at 4 and 8 years for the genes of the D1 medium spiny neuron (D1 MSN) marker set that change concordantly in RNA and protein at 8 years, in **(A)** RNA-seq and **(B)** age-normalised proteomics. Each line is one gene. Large points mark a FDR below 0.05 in the RNA-seq data. A point without a line (*NXPH4* in B) is a gene with no 4-year protein value. Abbreviations. D1 MSN, dopamine receptor D1 medium spiny neuron. FC, fold change. FDR, false discovery rate. TgHD, transgenic Huntington's disease minipig. WT, wild type.

### Supplementary Figure S9

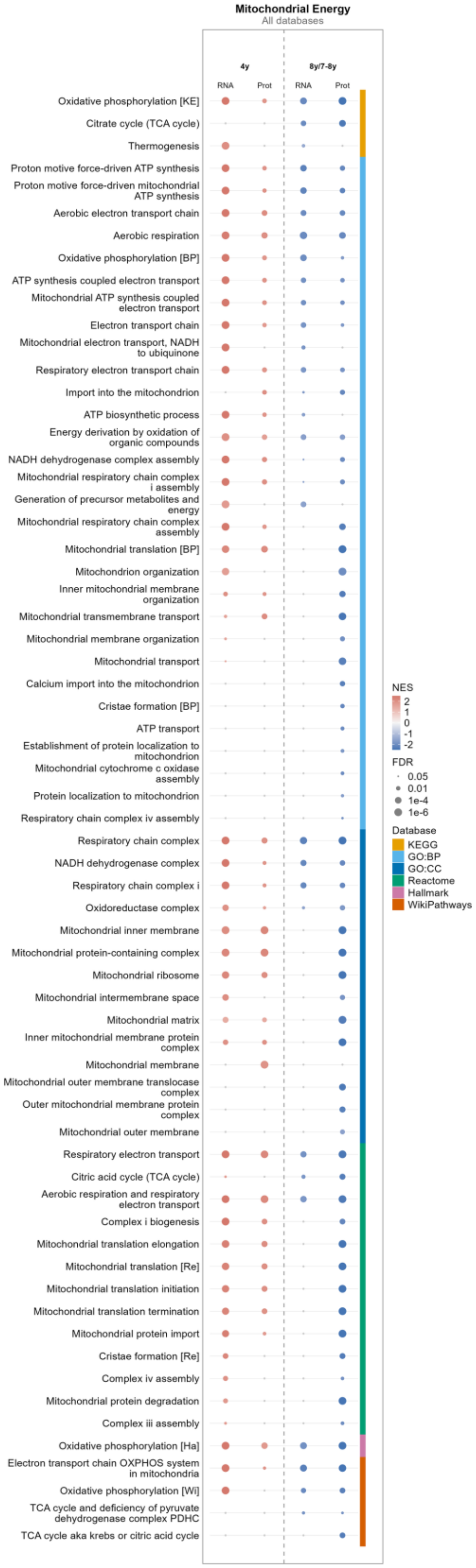

**Supplementary Fig. S9. Mitochondrial pathway enrichment across all pathway databases.** Dot heatmap of GSEA NES for the pathways assigned to the mitochondrial theme by keyword matching (Methods 2.9, Supplementary Table S7), for RNA-seq (RNA) and age-normalised proteomics (Prot) data at 4 years and at 8 years (RNA-seq) or 7 to 8 years (proteomics). Pathways come from KEGG, GO:BP and GO:CC, Reactome, Hallmark and WikiPathways, and the colour strip marks the source database. Pathways reaching a FDR below 0.05 in at least one data set were ranked by the number of data sets in which they reached significance, with pathways significant in either proteomics data set promoted in that ranking, and the 65 highest ranked are shown. Grey dots mark comparisons not reaching a FDR below 0.05. Within GO:BP and GO:CC, redundant parent terms were removed and gene sets with identical membership are shown once (Methods 2.10). A bracketed tag after a name distinguishes gene sets of the same name from different databases (KE, KEGG. BP, Gene Ontology biological process. Re, Reactome. Ha, Hallmark. Wi, WikiPathways). The full pathway list is in Supplementary Table S5. Abbreviations. FDR, false discovery rate. GO:BP, Gene Ontology biological process. GO:CC, Gene Ontology cellular component. GSEA, gene set enrichment analysis. NES, normalised enrichment score. TgHD, transgenic Huntington's disease minipig. WT, wild type.

### Supplementary Figure S10

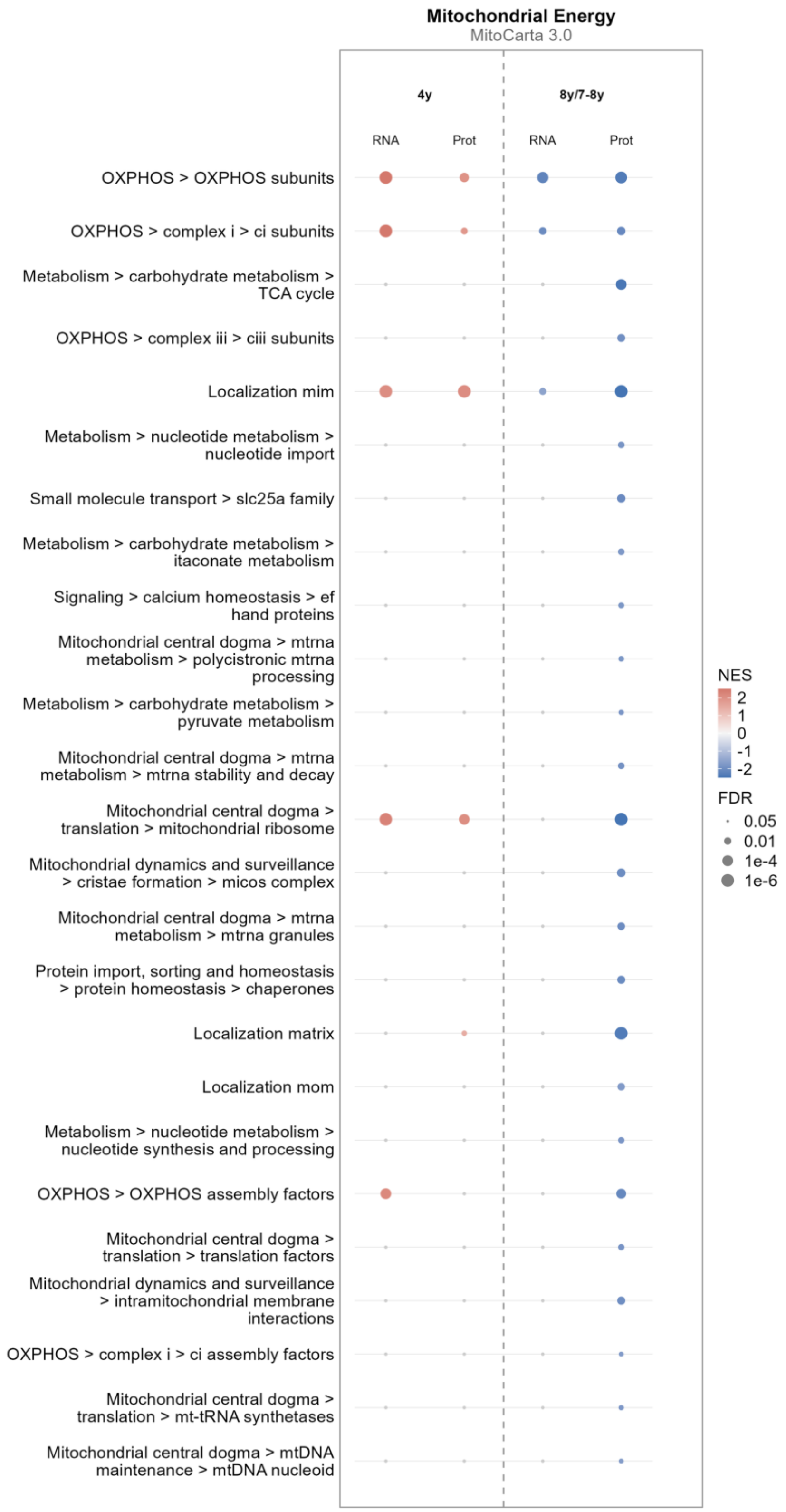

**Supplementary Fig. S10. Mitochondrial pathway enrichment of MitoCarta 3.0 gene sets.** Dot heatmap of GSEA NES for the MitoCarta 3.0 pathway gene sets (Rath et al., 2021), for RNA-seq (RNA) and age-normalised proteomics (Prot) data at 4 years and at 8 years (RNA-seq) or 7 to 8 years (proteomics). Row labels show the MitoCarta pathway hierarchy from the top level down, separated by >, and the rows labelled Localization are the MitoCarta sub-mitochondrial localisation sets (mim, inner membrane, mom, outer membrane). Gene sets reaching a FDR below 0.05 in at least one data set were ranked by the number of data sets in which they reached significance, with gene sets significant in either proteomics data set promoted in that ranking, and the 25 highest ranked are shown. Grey dots mark comparisons not reaching a FDR below 0.05. Abbreviations. FDR, false discovery rate. GSEA, gene set enrichment analysis. NES, normalised enrichment score. TgHD, transgenic Huntington’s disease minipig. WT, wild type.

### Supplementary Figure S11

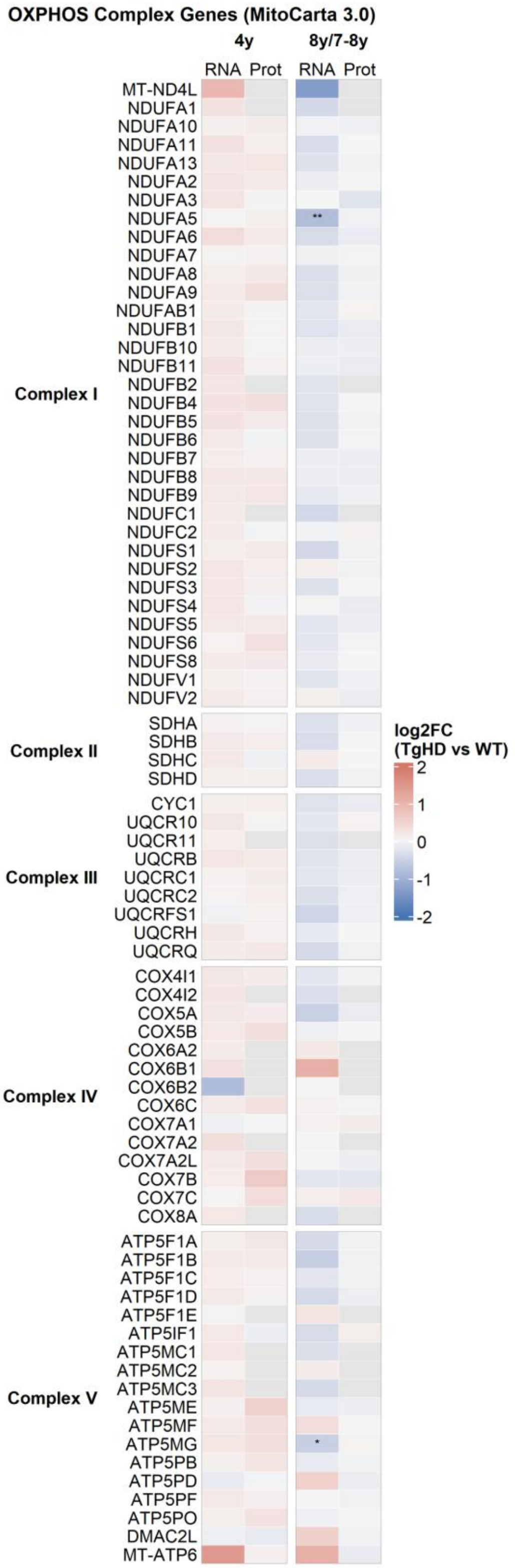

**Supplementary Fig. S11. Changes in the oxidative phosphorylation subunit genes.** Heatmap of log2 fold change, TgHD relative to WT, for the 79 OXPHOS subunit genes of MitoCarta 3.0, for RNA-seq (RNA) and age-normalised proteomics (Prot) data at 4 years and at 8 years (RNA-seq) or 7 to 8 years (proteomics), grouped by respiratory complex. Grey cells mark genes not detected in that data set. Asterisks mark FDR significance in the RNA-seq columns only, * below 0.05, ** below 0.01, *** below 0.001. Abbreviations. FDR, false discovery rate. OXPHOS, oxidative phosphorylation. TgHD, transgenic Huntington's disease minipig. WT, wild type.

### Supplementary Figure S12

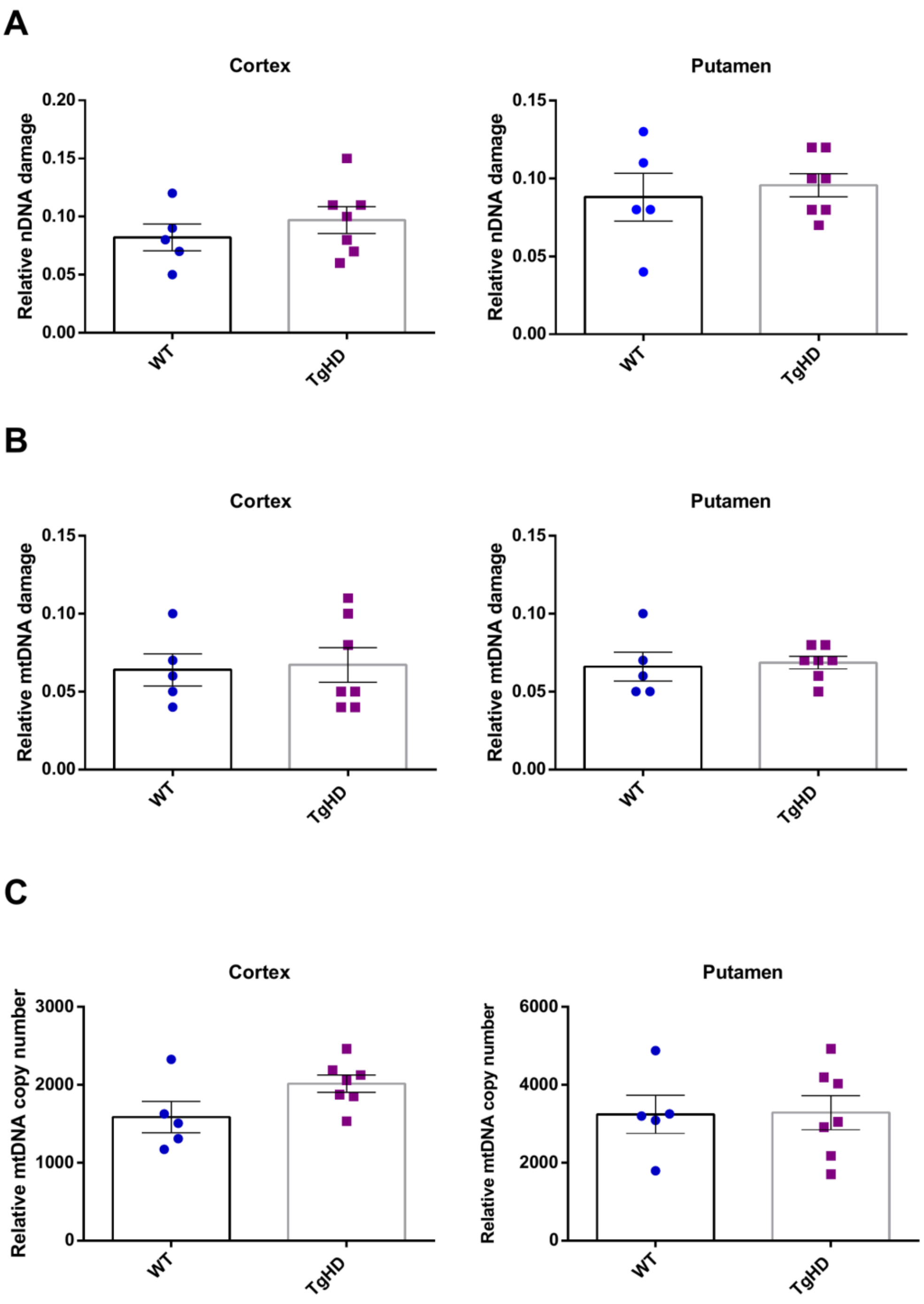


**Supplementary Fig. S12. Nuclear and mitochondrial DNA damage and mitochondrial DNA copy number at 7 years.** DNA damage was measured by RADF and mtDNA copy number by real-time qPCR in cortex and putamen of 7-year WT and TgHD animals. **(A)** Relative nuclear DNA (nDNA) damage in *NDUFA9*. **(B)** Relative mitochondrial DNA (mtDNA) damage in *MT-RNR1*. **(C)** Relative mtDNA copy number. Points are individual animals, bars the mean with SEM, n = 5 WT and 7 TgHD. Abbreviations. mtDNA, mitochondrial DNA. nDNA, nuclear DNA. SEM, standard error of the mean. qPCR, quantitative polymerase chain reaction. RADF, real-time qPCR analysis of damage frequency. TgHD, transgenic Huntington's disease minipig. WT, wild type.

#### Supplementary Figure S13

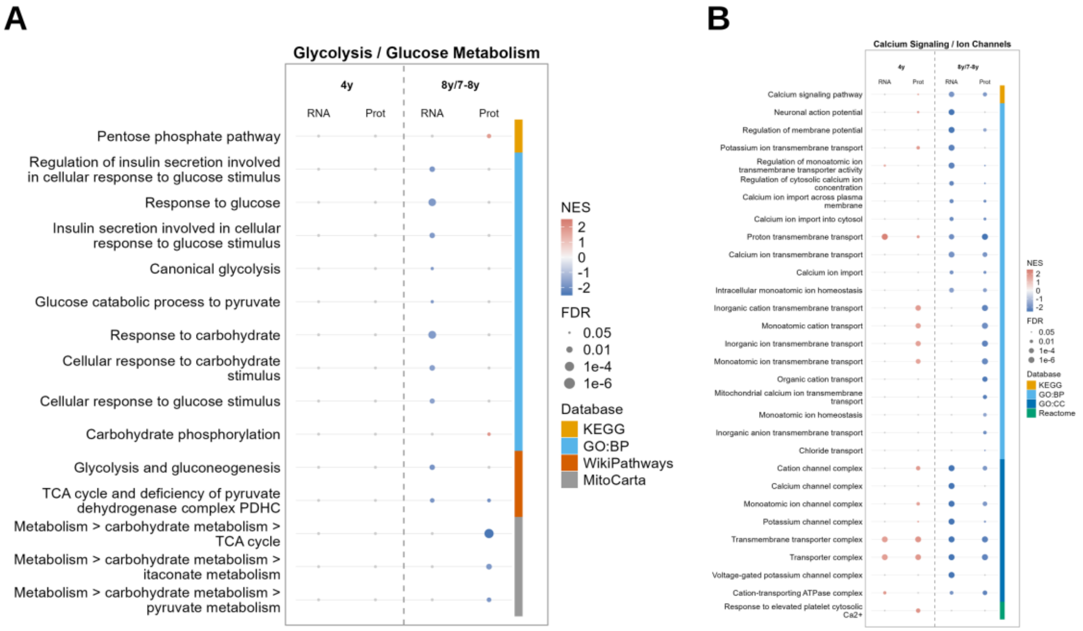


**Supplementary Fig. S13. Carbohydrate metabolism and calcium-handling pathway enrichment.** Dot heatmap of GSEA NES for RNA-seq (RNA) and age-normalised proteomics (Prot) data at 4 years and at 8 years (RNA-seq) or 7 to 8 years (proteomics), for **(A)** glycolysis and carbohydrate metabolism gene sets and **(B)** calcium-handling and ion-channel gene sets, both assigned by keyword matching (Methods 2.9, Supplementary Table S7). The colour strip marks the source database. In each panel, gene sets reaching a FDR below 0.05 in at least one data set were ranked by the number of data sets in which they reached significance, with gene sets significant in either proteomics data set promoted in that ranking, and the highest ranked are shown, up to 25 in (A) and up to 30 in (B). Grey dots mark comparisons not reaching a FDR below 0.05. In (A) the MitoCarta rows are labelled with the MitoCarta pathway hierarchy, separated by >. The full list is in Supplementary Table S5. Abbreviations. FDR, false discovery rate. GSEA, gene set enrichment analysis. NES, normalised enrichment score. TgHD, transgenic Huntington's disease minipig. WT, wild type.

#### Supplementary Figure S14

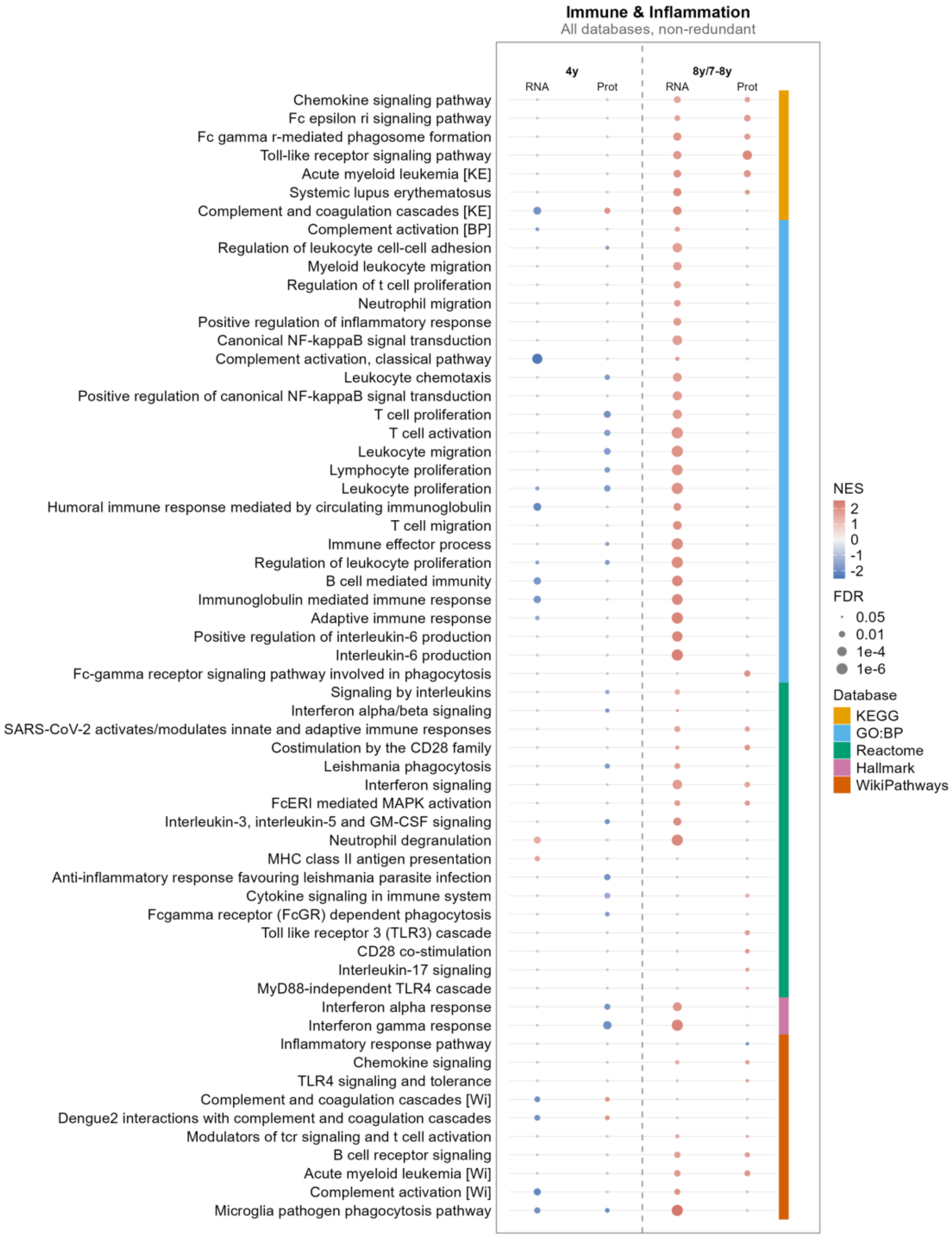


**Supplementary Fig. S14. Immune and inflammatory pathway enrichment across all pathway databases.** Dot heatmap of GSEA NES for the pathways assigned to the immune and inflammation theme by keyword matching (Methods 2.9, Supplementary Table S7), for RNA-seq (RNA) and age-normalised proteomics (Prot) data at 4 years and at 8 years (RNA-seq) or 7 to 8 years (proteomics). The colour strip marks the source database. Pathways reaching a FDR below 0.05 in at least one data set were ranked by the number of data sets in which they reached significance, with pathways significant in either proteomics data set promoted in that ranking, and the 60 highest ranked are shown. Grey dots mark comparisons not reaching a FDR below 0.05. Within GO:BP and GO:CC, redundant parent terms were removed and gene sets with identical membership are shown once (Methods 2.10). A bracketed tag after a name distinguishes gene sets of the same name from different databases (KE, KEGG. BP, Gene Ontology biological process. Wi, WikiPathways). The full pathway list is in Supplementary Table S5. Abbreviations. FDR, false discovery rate. GO:BP, Gene Ontology biological process. GO:CC, Gene Ontology cellular component. GSEA, gene set enrichment analysis. NES, normalised enrichment score. TgHD, transgenic Huntington's disease minipig. WT, wild type.

#### Supplementary Figure S15

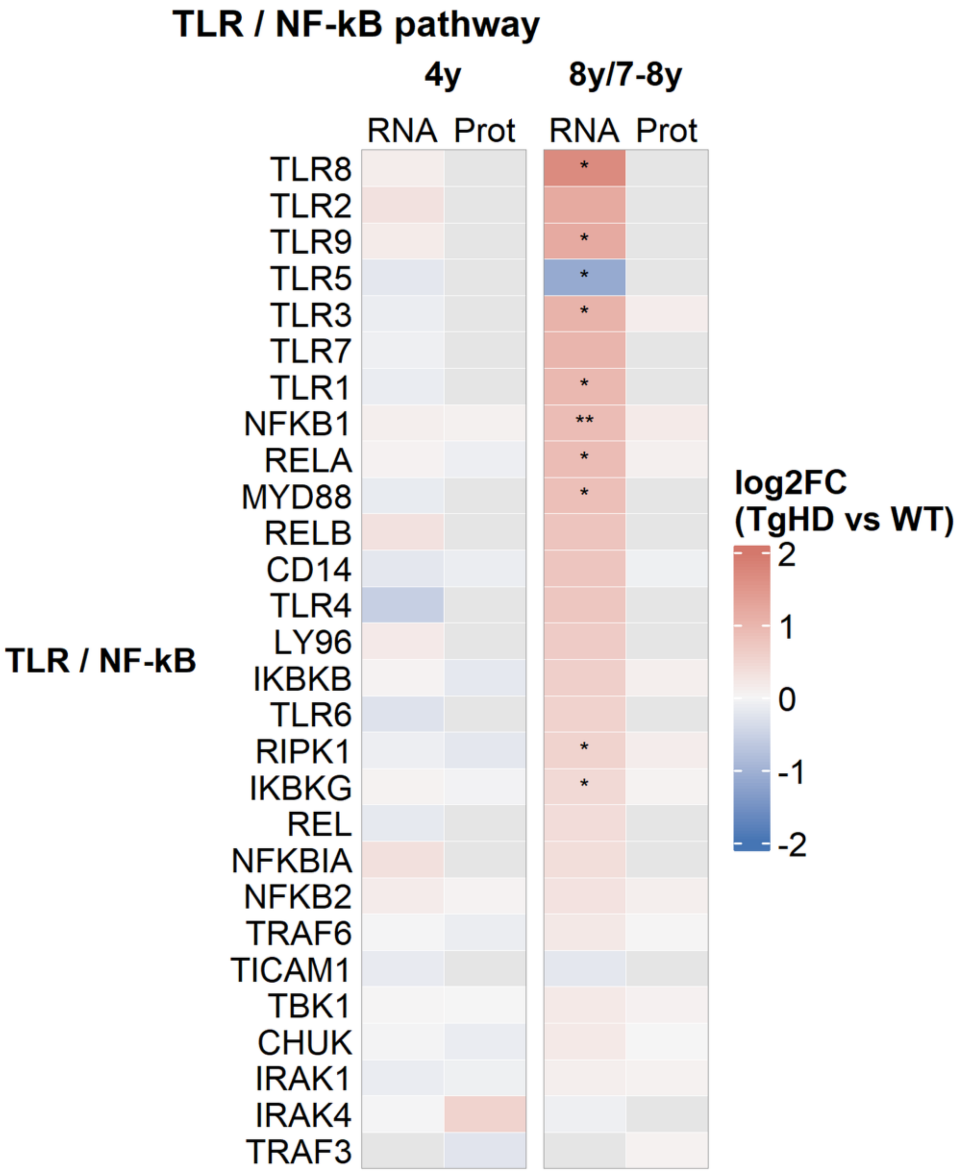


**Supplementary Fig. S15. Toll-like receptor and NF-κB pathway genes.** Heatmap of log2 fold change, TgHD relative to WT, for the TLR and nuclear factor kappa B (NF-κB) signalling gene set (Supplementary Table S8), for RNA-seq (RNA) and age-normalised proteomics (Prot) data at 4 years and at 8 years (RNA-seq) or 7 to 8 years (proteomics). Grey cells mark genes not detected in that data set. Asterisks mark FDR significance in the RNA-seq columns only, * below 0.05, ** below 0.01, *** below 0.001. Abbreviations. FDR, false discovery rate. NF-κB, nuclear factor kappa B. TLR, Toll-like receptor. TgHD, transgenic Huntington's disease minipig. WT, wild type.

#### Supplementary Figure S16

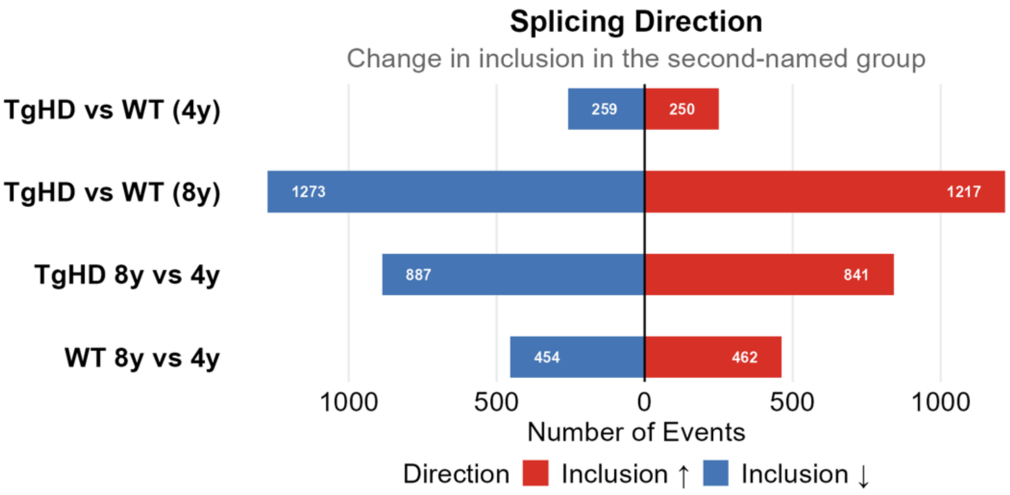


**Supplementary Fig. S16. Direction of the alternative splicing changes.** Diverging bar chart of significant alternative splicing events per comparison, split by the direction of the change in per cent spliced in (ΔPSI) in the second-named group, TgHD in the genotype comparisons and 8 years in the age comparisons. Event counts are printed on the bars. Supplementary Table S9 lists all events with the rMATS sign, first group minus second group. Abbreviations. ΔPSI, change in per cent spliced in. TgHD, transgenic Huntington's disease minipig. WT, wild type.

#### Supplementary Figure S17

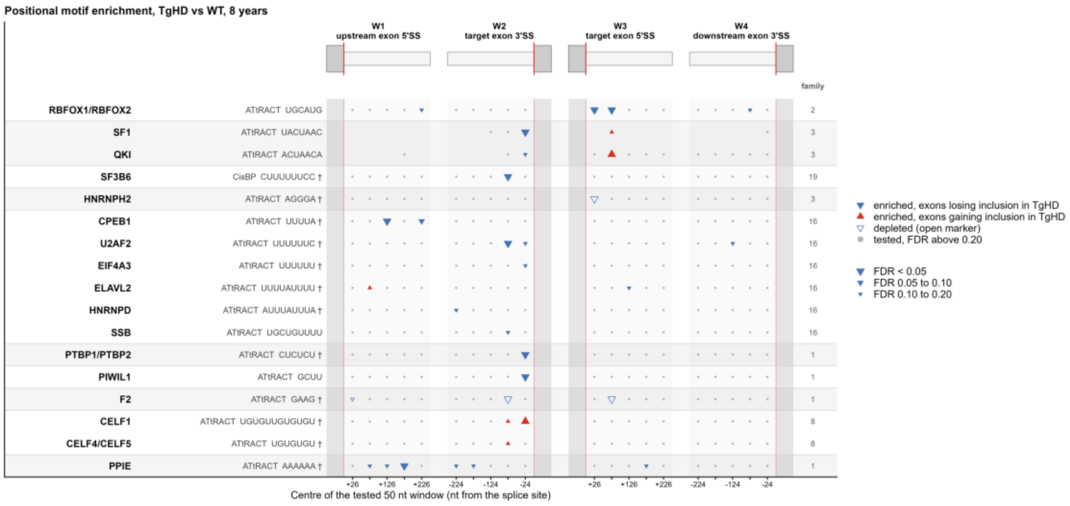


**Supplementary Fig. S17. Positional RNA-binding protein motif enrichment around skipped exons at 8 years.** Each row is one motif, a position weight matrix from the ATtRACT or CisBP-RNA collection (printed as CisBP), named after the protein the collection assigns it to, with the collection and the consensus sequence beside the name. A row appears when its motif family contains a motif with a FDR below 0.05 and the row itself reaches FDR below 0.20, which gives 17 rows in 10 families. Proteins whose motifs have identical values at every tested position share one row. Alternating shading groups the rows of one family, motifs whose matrices correlate at Pearson r of at least 0.85, so one shaded block counts as one observation. The family column gives the number of motifs in that family, including those not shown. A dagger marks a low-complexity consensus (two or fewer distinct bases, or one base at 75 percent or more), which cannot identify a single protein. Columns are the centres of the tested 50 nt windows at the four splice sites of a skipped-exon event, under a schematic of the exon (grey box) and the adjacent intron, with the red line at the splice site. Grey columns are exonic and were not tested. A position without a marker had fewer than 10 exons with the motif and was not tested (Methods 2.12). Fig. 7D shows the RBFOX rows at single-nucleotide resolution. Supplementary Table S10 lists the full results for all four comparisons. Abbreviations. FDR, false discovery rate. TgHD, transgenic Huntington's disease minipig. WT, wild type.

#### Supplementary Figure S18

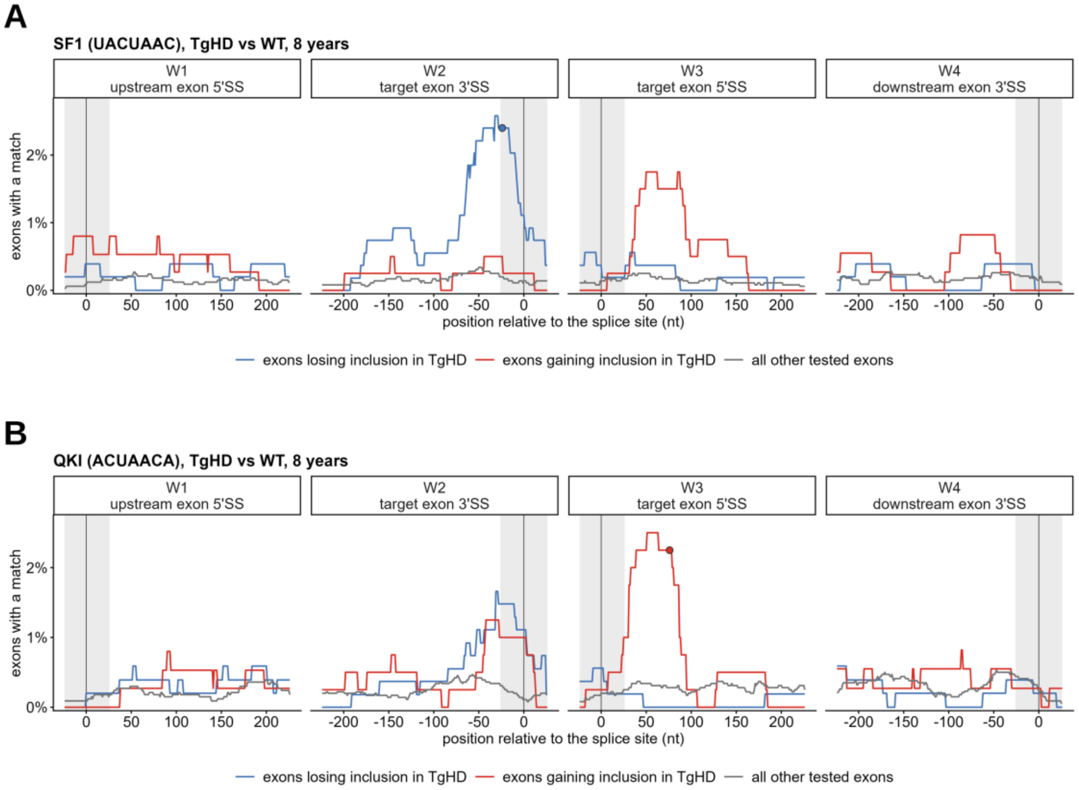


**Supplementary Fig. S18. Positional motif enrichment for SF1 and QKI around skipped exons at 8 years.** Motif maps for **(A)** the SF1 motif UACUAAC, the branch point consensus, and **(B)** the QKI motif ACUAACA, TgHD relative to WT at 8 years. The two motifs overlap in sequence and belong to one motif family, so they appear together. Windows W1 to W4 cover the four splice sites of a skipped-exon event, the 5’ splice site of the upstream exon, the 3’ and 5’ splice sites of the target exon and the 3’ splice site of the downstream exon, with position 0 at the splice site. The y axis gives the percentage of exons with a motif match in the 50 nt window centred at that position, for the three exon groups in the legend. Grey bands cover windows that still contain exonic sequence, which were not tested. The test compares each group with the exons whose inclusion did not change (Methods 2.12), and a filled circle marks a position with a FDR below 0.05. Supplementary Table S10 lists the full results. Abbreviations. FDR, false discovery rate. SS, splice site. TgHD, transgenic Huntington’s disease minipig. WT, wild type.

#### Supplementary Figure S19

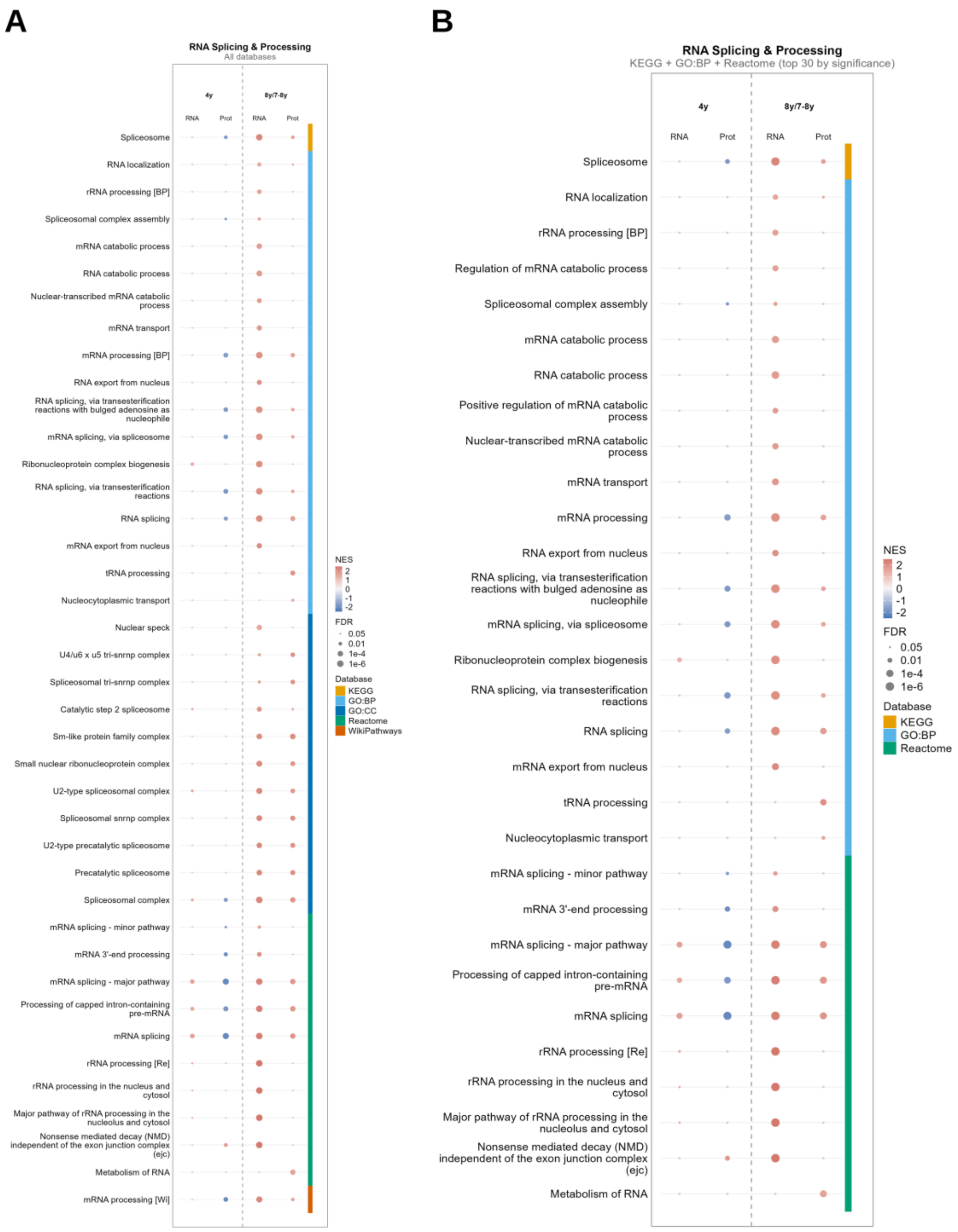


**Supplementary Fig. S19. Splicing and RNA-processing pathway enrichment.** Dot heatmap of GSEA NES for the pathways assigned to the RNA splicing and processing theme by keyword matching (Methods 2.9, Supplementary Table S7), for RNA-seq (RNA) and age-normalised proteomics (Prot) data at 4 years and at 8 years (RNA-seq) or 7 to 8 years (proteomics), from **(A)** all pathway databases and **(B)** KEGG, GO:BP and Reactome only.

The colour strip marks the source database. In each panel, pathways reaching a FDR below 0.05 in at least one data set were ranked by the number of data sets in which they reached significance, with pathways significant in either proteomics data set promoted in that ranking, and the highest ranked are shown, up to 40 in (A) and up to 30 in (B). Grey dots mark comparisons not reaching a FDR below 0.05. Within GO:BP and GO:CC, redundant parent terms were removed and gene sets with identical membership are shown once (Methods 2.10). A bracketed tag after a name distinguishes gene sets of the same name from different databases (BP, Gene Ontology biological process. Re, Reactome. Wi, WikiPathways). The full pathway list is in Supplementary Table S5. Abbreviations. FDR, false discovery rate. GO:BP, Gene Ontology biological process. GO:CC, Gene Ontology cellular component. GSEA, gene set enrichment analysis. NES, normalised enrichment score. TgHD, transgenic Huntington's disease minipig. WT, wild type.

---

#### Supplementary tables

| Table | Content |
|---|---|
| S1 | Sample overview, animals, ages, sex, perfusion |
| S2 | Differential gene expression (DESeq2), the five comparisons |
| S3 | RNA-seq count matrix, raw and variance-stabilised, with sample information |
| S4 | Proteomics differential abundance (limma, age-normalised), 4-year, 8-year and 7- to 8-year comparisons |
| S5 | Gene set enrichment analysis, all databases and comparisons |
| S6 | Cell-type marker set enrichment |
| S7 | Pathway metadata and theme assignment |
| S8 | Cell-type marker gene sets, provenance and scoring |
| S9 | Alternative splicing events (rMATS) |
| S10 | RNA-binding protein positional motif enrichment |
| S11 | Validation correlations, qPCR against RNA-seq and RNA against protein |
| S12 | Gene annotation and provenance |
| S13 | ChEA3 transcription factor enrichment |

| Table | Content |
|---|---|
| S14 | R package versions |
| S15 | Leading-edge gene overlap between pathways |
| S16 | Over-representation analysis of alternatively spliced genes |
| S17 | qPCR primer sequences |
| S18 | Two-way ANOVA statistics for the HTRF assays of Supplementary Fig. S5 |
